# Spatial constraints and cell surface molecule depletion structure a randomly connected learning circuit

**DOI:** 10.1101/2024.07.17.603956

**Authors:** Emma M. Thornton-Kolbe, Maria Ahmed, Finley R. Gordon, Bogdan Sieriebriennikov, Donnell L. Williams, Yerbol Z. Kurmangaliyev, E. Josephine Clowney

## Abstract

The brain can represent almost limitless objects to “categorize an unlabeled world” (Edelman, 1989). This feat is supported by expansion layer circuit architectures, in which neurons carrying information about discrete sensory channels make combinatorial connections onto much larger postsynaptic populations. Combinatorial connections in expansion layers are modeled as randomized sets. The extent to which randomized wiring exists *in vivo* is debated, and how combinatorial connectivity patterns are generated during development is not understood. Non-deterministic wiring algorithms could program such connectivity using minimal genomic information. Here, we investigate anatomic and transcriptional patterns and perturb partner availability to ask how Kenyon cells, the expansion layer neurons of the insect mushroom body, obtain combinatorial input from olfactory projection neurons. Olfactory projection neurons form their presynaptic outputs in an orderly, predictable, and biased fashion. We find that Kenyon cells accept spatially co-located but molecularly heterogeneous inputs from this orderly map, and ask how Kenyon cell surface molecule expression impacts partner choice. Cell surface immunoglobulins are broadly depleted in Kenyon cells, and we propose that this allows them to form connections with molecularly heterogeneous partners. This model can explain how developmentally identical neurons acquire diverse wiring identities.

## Introduction

Expansion layers are a class of circuit in which a large set of neurons receive combinatorial input from a smaller population of presynaptic cells (Albus, 1971; M. Ito, 1970; Marr, 1969).

Typically, the presynaptic cells bring different pieces of sensory information, and the postsynaptic cells act as coincidence detectors, i.e. they fire only when multiple of their inputs are active (Chabrol et al., 2015; Gruntman & Turner, 2013; Huang et al., 2013; Ishikawa et al., 2015). Expansion coding therefore increases the number of experiences that the brain can represent from the set of sensory channels to the much larger set of combinations among them (Cayco-Gajic et al., 2017; Litwin-Kumar et al., 2017). Expansion layers are found in each of the major clades of Bilateria and seem to have evolved multiple times; they include the parallel lobe system of cephalopods, the arthropod mushroom body, and the vertebrate pallium, hippocampus, and cerebellum (Farris, 2011; Hobbs & Young, 1973; Leutgeb et al., 2007; Sawtell, 2010; Srinivasan & Stevens, 2018). These circuit architectures are often used for associative learning, in which previously arbitrary sensory representations become connected with temporally linked events.

From a developmental point of view, two wiring features are critical for expansion coding: density and identity of inputs. For expansion coding to improve pattern separation, expansion layer neurons must receive only some of the presynaptic inputs, not all of them. Principle neurons in each kind of expansion layer receive characteristic densities of presynaptic inputs (Ahmed et al., 2023; Barak et al., 2013; Jortner et al., 2007; A. C. Lin et al., 2014; Litwin-Kumar et al., 2017). In cerebellum-like expansion layers (including the mushroom body, mormyrid fish electric organ, and dorsal cochlear nucleus) each expansion layer neuron receives a small number of discrete inputs, usually less than 10 (Caron et al., 2013; Cayco-Gajic & Silver, 2019; Leiss et al., 2009; Mugnaini et al., 1980; Sawtell, 2010). In our previous work, we found that in the mushroom body, this input density is set during development by the expansion layer neurons themselves, called Kenyon cells, who instruct the input cells to produce the appropriate set of presynaptic structures (Ahmed et al., 2023; Elkahlah et al., 2020). Next, individual expansion layer neurons must receive input from different sets of presynaptic neurons, such that different neurons in the expansion layer are sensitive to different combinations of stimuli (Albus, 1971; Cayco-Gajic et al., 2017; Gruntman & Turner, 2013; M. Ito, 1970; Litwin-Kumar et al., 2017; Marr, 1969). How these diverse, combinatorial connections emerge during development is unknown and is unlikely to mimic the development of wiring observed in motor or sensory structures.

In those better-studied circuits, developmentally diverse neurons connect to one another in precise ways. In contrast, expansion layer neurons tend to be more numerous and developmentally homogenous yet connect to distinct partners. We usually think of the relative “fitness” of a neural circuit architecture only in terms of its marginal improvement of information processing function in the adult. However, given the large number of expansion layer neurons, the compactness of the developmental information required to wire them is also a critical aspect of fitness. For example, the mammalian cerebellum can comprise tens of billions of granule cells; a finite genome cannot deterministically specify unique inputs to each of these billions of cells. Because meaning need not be hard coded in learning circuits and is instead conferred by experience, non-deterministic connectivity is compatible with function (Minsky, 1952, 2016; Rosenblatt, 1958). A non-deterministic developmental algorithm for wiring could require less genomic information and still produce a functional circuit.

The *Drosophila melanogaster* mushroom body provides an excellent model to explore mechanisms that allow expansion layer neurons to receive diverse sets of presynaptic inputs. In this brain region, ∼2000 Kenyon cells each receive 3-10 discrete inputs from the olfactory system (Caron et al., 2013; Leiss et al., 2009). Kenyon cells are developmentally homogenous; they are born from four neural stem cells (neuroblasts) and comprise only seven major anatomic subtypes in the adult (Aso et al., 2009; K. Ito et al., 1997; Kunz et al., 2012). Kenyon cell types are born sequentially, and each of the four neuroblasts makes all types in parallel (T. Lee et al., 1999). >90% of Kenyon cells, including all but two major types, receive almost all of their input from olfactory projection neurons (Aso et al., 2009, 2014; F. Li et al., 2020). The olfactory projection neurons providing presynaptic inputs to Kenyon cells are much more developmentally diverse—these ∼100 neurons are of 51 anatomic types, and arise in a precise and reproducible order from two distinct neuroblasts (S. Lin et al., 2012; H.-H. Yu et al., 2010).

Here, we ask how Kenyon cells with shared specification programs can nevertheless acquire distinct combinations of presynaptic inputs during development. There has been enduring debate in the field about the extent to which Kenyon cell inputs are random (Caron et al., 2013; Eichler et al., 2017; Ellis et al., 2023; Ganguly et al., 2024; Gruntman & Turner, 2013; Hayashi et al., 2022; Jefferis et al., 2004; F. Li et al., 2020; Marin et al., 2002; Murthy et al., 2008; Tanaka et al., 2004; Wong et al., 2002; J.-Y. Yang et al., 2023; Zheng et al., 2022), and a similar debate emerges in the cerebellum literature (Bengtsson & Jörntell, 2009; Gilmer & Person, 2017; Kennedy et al., 2014; Shuster et al., 2021). Therefore, we first analyze electron microscopy connectomic data (Scheffer et al., 2020) and compare it to previous light microscopy studies (Jefferis et al., 2007; T. Lee et al., 1999; H.-H. Lin et al., 2007; Marin et al., 2002; Tanaka et al., 2004; Wong et al., 2002; Zhu et al., 2003) to provide a precise characterization of spatial distributions that bias the sets of inputs to Kenyon cells of different neuroblast clones or types. Unifying previous reports, we find that projection neurons of different types innervate unique and reproducible regions within the mushroom body calyx (Jefferis et al., 2007; H.-H. Lin et al., 2007; Tanaka et al., 2004; Zheng et al., 2022). We find that individual Kenyon cells are spatially constrained depending on their neuroblast origin and subtype, and that these spatial constraints, overlaid on the orderly projection neuron array, set up predictable biases in the boutons that can be reached by individual Kenyon cells. *Within* domains defined by their subtype and neuroblast origin, Kenyon cells sample randomly from the available projection neuron distribution.

Through what molecular mechanisms do Kenyon cells acquire random inputs from diverse presynaptic partners? The classic framing of synaptic partner matching sets up a debate between Sperry, who posited that molecular specificity factors guide matching, and Peters, who argued that neurite spatial targeting provides the primary constraint on connectivity (Peters & Feldman, 1976; Sperry, 1963). Olfactory projection neurons innervating particular glomeruli are born in a reproducible temporal sequence and express distinct repertoires of cell surface molecules (Hongjie Li et al., 2017; S. Lin et al., 2012; Xie et al., 2021; H.-H. Yu et al., 2010). Therefore, we next characterize the transcriptional programs of Kenyon cells across pupal development to ask how neurons of homogenous developmental identity can synapse with molecularly diverse partners to acquire distinct wiring identities. We find that Kenyon cells have a transcriptional depletion of immunoglobulin superfamily proteins. Genes in this family are expressed in diverse and specific patterns in other kinds of neurons, including olfactory projection neurons, and are proposed to regulate a variety of type-specific developmental processes, including synaptic partner choice (Carrillo et al., 2015; Hongjie Li et al., 2017; C. Xu et al., 2019; Yoo et al., 2023). We propose that downregulation of immunoglobulins in Kenyon cells serves to blind them to the molecular differences among incoming olfactory projection neurons. Each Kenyon cell is then free to obtain inputs promiscuously from among the projection neuron boutons available within its domain, generating diverse wiring inputs to developmentally homogenous cells.

## Results

### Projection neuron anatomy in the mushroom body calyx is orderly, biased, and predictable

In the Hemibrain EM connectome, there are 105 cholinergic projection neurons (PNs) that each innervate one of 51 olfactory antennal lobe glomeruli where they synapse with one type of olfactory sensory neuron (Figure 1A-B). In the Hemibrain calyx, these PNs provide input to 1728 olfactory Kenyon cells (Figure 1A). PN axonal termini in the calyx, called boutons, are bulbous structures containing multiple presynaptic sites (Leiss et al., 2009; K. Yang et al., 2022; Yusuyama et al., 2002). Each Kenyon cell produces between 3 and 10 claw-shaped dendrites that each enwrap a bouton (Caron et al., 2013; Leiss et al., 2009). Thus, each Kenyon cell receives information from 3- 10 of the 51 possible antennal lobe glomeruli (Figure 1C).

**Figure 1.**
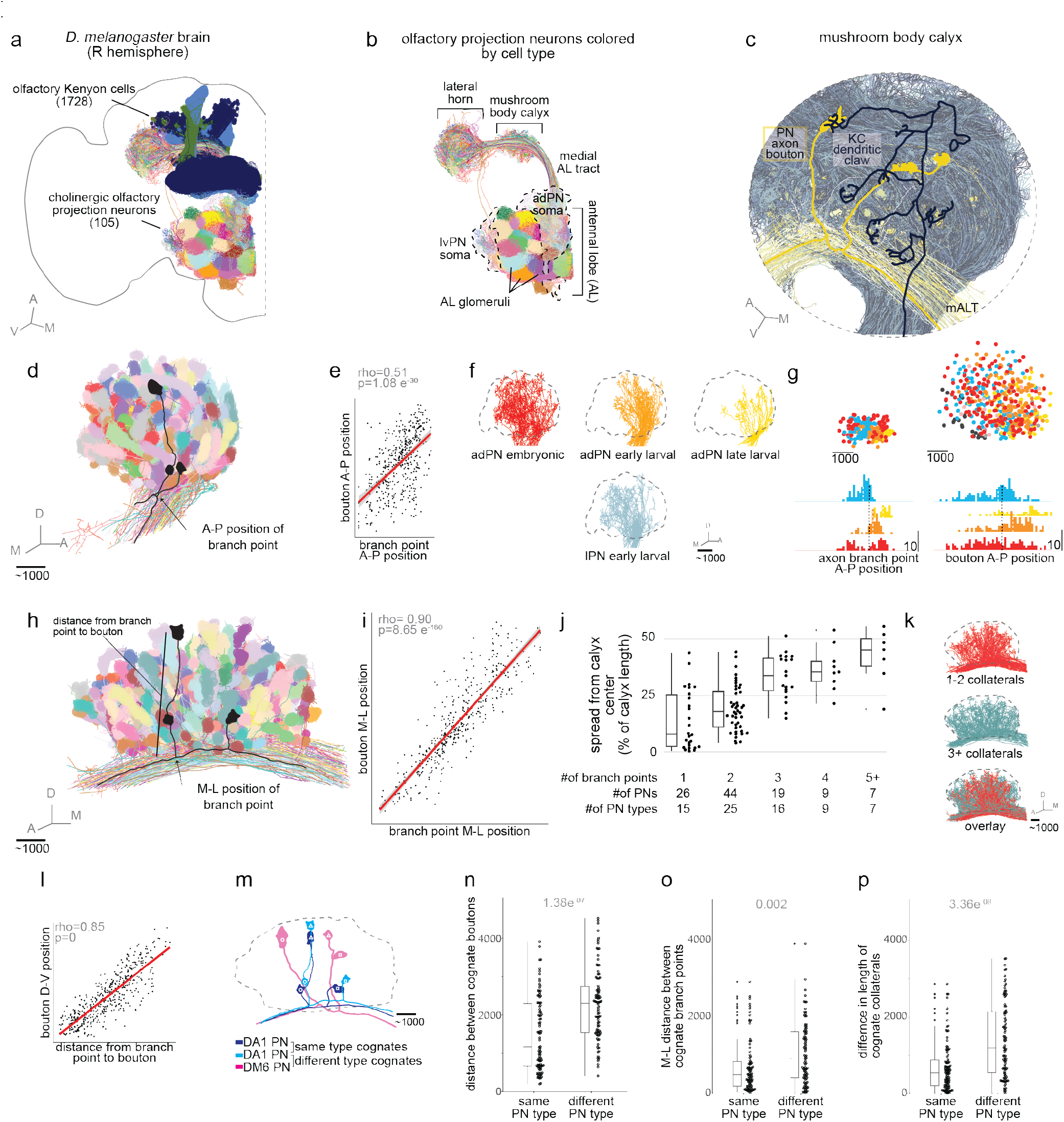
**(a,b)** Skeletons of olfactory Kenyon cells (a) and uniglomerular olfactory projection neurons (a,b) from the Hemibrain EM connectome, colored by type. Anatomic axes used to orient the calyx are shown in (a) (see methods). In (b), Kenyon cells are removed to allow visualization of the calyx, and dotted lines outline the locations of PN soma. PN types segregate in the antennal lobe and mix in the calyx. **(c)** Skeletons of KCs (blue shades) and PNs (yellow shades) in the calyx with a single γ KC and DM6 projection neuron highlighted. Each dendritic claw of a KC enwraps a single PN bouton and each bouton is enwrapped by an average of 21 dendritic claws. **(d)** A-P view of olfactory PNs skeletons in the calyx, colored by type. Here and throughout, bouton shapes are approximated from centroids of enwrapping KC claws. A single DA1 PN is traced in black. **(e)** The A-P coordinate of each PN bouton centroid is well correlated with the A-P coordinate of the branch point collateral on which it resides (Spearman’s rank correlation, rho=0.51, p = 1.08e-30, n=436 boutons). Here and throughout, red line is a linear model and shaded grey represents 95% confidence interval. **(f)**A-P view of olfactory PN skeletons colored and separated by neuroblast origin and birth stage. Dotted outline represents the bounds of the calyx. **(g)** Axon branch point position (left) or bouton position (right) along the A-P (x-axis) and D-V (y-axis) for all collateral branch points colored by neuroblast origin and birth stage. Histograms of branch point (n=246) and bouton centroid (n=436) positions are displayed below with a bin width of 100 pixels. **(h)** M-L view of olfactory PN skeletons, plotted as in (d). The black PN is the same DA1 PN as in (d). A single DA1 PN is traced in black along with example branch-point-to-bouton measurement. **(i)** The M-L coordinate of each PN bouton centroid is well correlated with the M-L coordinate of the branch point collateral on which it resides (Spearman’s rank correlation, rho=0.90, p=8.65e-160, n=436 boutons). **(j)** For each PN, maximum distance of a collateral from the center of the calyx along the M-L axis, expressed as percent of calyx length. M-L center is defined as the average M-L branch point of all collaterals. Most PNs with 1-2 collaterals branch within the central half of the calyx (a quarter of the length to either side), while PNs with more than 2 collaterals can place branches more distantly. In box plots: center line, median; box limits, upper and lower quartiles; whiskers, 1.5x interquartile range; points, outliers. **(k)** M-L view of PN skeletons colored by number of collaterals. Dotted outline represents the bounds of the calyx. Branches of PNs with 1-2 collaterals cluster in the central calyx, while PNs making more branches can distribute them more broadly. **(l)** The D-V coordinate of each PN bouton centroid is well correlated with its linear distance from the collateral branch point of which it resides (Spearman’s rank correlation, rho=0.85, p=0.00, n=436 boutons). **(m)** Example of cognate PNs, branches, and boutons: We matched PNs to cognates with the same number of branch points and boutons. Within cognate groups, cognate branch points and boutons are those that occupy the same M-L ordinal position. DA1 PNs (grey and black; black PN is the PN highlight in (d) and (h)) represent cognate PNs of the same type. The DM6 PN (purple) is a cognate of a different type. Cognate boutons on each PN are labeled with the same shape. **(n)** Linear distance between cognate boutons on PNs of the same type (n=98, median=1172.79) and those on different types (n=98 randomly sampled from 2263, median =2310.27). (Wilcoxon rank sum test; U=2710, p=1.38e^-07^, Cohen’s D=0.81). In box plots: center line, median; box limits, upper and lower quartiles; whiskers, 1.5x interquartile range; points, outliers. **(o)** Distance along the M-L axis between cognate branch points of PNs of the same type (n=88, median=490.25) and those on a different type (n=88 randomly sampled from 2818, median = 910.27). (Wilcoxon rank sum test, U=2821, p=0.002, Cohen’s D=0.35) **(p)** Distance along the M-L axis between cognate branch points of PNs of the same type (n=98, median=538.12) and those on a different type (n=98 randomly sampled from 2185, median=1186.99). (Wilcoxon rank sum test, U=2609, p=3.36e^-08^, Cohen’s D=0.86) Even among matched cognates, structures of PNs of the same type are more alike than those of different types.

Uniglomerular olfactory PNs are produced by invariant neural lineages and innervate predictable antennal lobe glomeruli depending on their birth order (Alexander S. Bates et al., 2020; Grabe et al., 2016; S. Lin et al., 2012; H.-H. Yu et al., 2010). Each glomerulus in the antennal lobe is innervated by between 1 and 7 PNs, which is reproducible from animal to animal (Grabe et al., 2016). PNs of particular types make between 1 and 15 boutons in the calyx; while the total number of boutons in the calyx is influenced by the number of Kenyon cells and PNs in a particular brain, the relative contributions of different PN types is predictable and consistent (Alexander S. Bates et al., 2020; Caron et al., 2013; Elkahlah et al., 2020; Hayashi et al., 2022; Jefferis et al., 2007; H.-H. Lin et al., 2007; Marin et al., 2002; Tanaka et al., 2004; Wong et al., 2002; Zheng et al., 2022). Many early light microscopy studies identified clear and predictable differences in the positioning of neurites and boutons of different PN types (Jefferis et al., 2007; H.-H. Lin et al., 2007; Marin et al., 2002; Tanaka et al., 2004; Wong et al., 2002). Using the Hemibrain connectome, we sought (1) to situate the anatomies of individual PN types within a holistic description of bouton placement; (2) to identify developmental variables producing these anatomies; and (3) to build a developmental framework that reconciles predictable PN development with variation in Kenyon cell connectivity.

To describe the anatomic organization of PNs in the calyx, we identified 24,791 synapses from olfactory PNs to olfactory Kenyon cells within the neuropil defined as “calyx” and then clustered these synapses into boutons (see Methods). To simplify further analyses, we rotated coordinates, which are tilted relative to the whole brain, to align with Cartesian axes (Figure S1A-B, Methods).

First we looked at the positioning of boutons across the 20 micron anterior-posterior (A-P) span of the calyx (Figure 1D). The PN axons in the medial antennal lobe tract (mALT) are perpendicular to this axis along the ventral side of the calyx such that each axon occupies a distinct A-P position. The position of the axon from which a bouton arises predicts its A-P position (Figure 1E). Previous work noted that the axons are segregated in the mALT based on the developmental stage of their birth and neuroblast origin (Jefferis et al., 2007). We found this pattern to be consistent across the 51 types of olfactory PNs in the Hemibrain dataset: axons and boutons of neurons born embryonically are broadly distributed along the A-P axis, while axons and boutons of larvally-born PNs from the anterodorsal PN (adPN) and lateral PN (lPN) lineages are biased toward anterior versus posterior domains, respectively (Figure 1F,G). These patterns are sharper for axons than for boutons, partly due to the posterior lean of the collaterals on which the boutons are placed which results in more intermixing of boutons from different lineages and birth stages.

We next looked at positioning of boutons along the 50 micron medial-lateral (M-L) axis of the calyx (Figure 1H). While some boutons form directly on the axon, most form at the end of collaterals which branch perpendicularly from the mALT and continue in a straight line dorsally such that the M-L position of a bouton is strongly predicted by the M-L position of the branch point from the main axon (Figure 1I). Individual PNs make type-specific numbers of collaterals and place them in type-specific places across the calyx. While some neurons give rise to several major collaterals evenly distributed across the calyx, others biasedly innervated certain M-L compartments (H.-H. Lin et al., 2007). PNs in the Hemibrain have between 1 and 7 collaterals; neurons and neuron types are almost evenly split among those making one, two, or ≥ three collaterals (Figure 1J). We found that most neurons with just one or two collaterals placed those collaterals toward the center of the calyx while those with 3 or more collaterals spread them further along the M-L axis (Figure 1J,K). These highly branched PNs populating the periphery of the Hemibrain calyx are the same PN types observed as the outermost bouton shell in Tanaka 2004 and as the “overconvergent fovea” in Zheng 2022 (Figure S1C).

Finally, we described the positioning of boutons along the 30 micron dorsal-ventral axis of the calyx. While boutons are most densely packed in the ventral calyx closer to the axons, some are placed more dorsally at the end of long collaterals. Indeed, distance between the collateral branch point and bouton (a proxy for collateral length) predicts the dorsal-ventral positioning of the bouton (Figure 1L). Collateral length varies equally across both the A-P and M-L axis.

In the Hemibrain, 29 of 51 PN types have multiple member neurons who innervate the same AL glomeruli and whose anatomy closely mirror one another in the calyx. To compare the locations of boutons between individual PNs of the same type we measured the distance between cognate boutons on different neurons. We defined cognate boutons as ones which occupy the same M-L ordinal position, and only compared neurons with the same number of boutons. As a control, we also measure distances between cognate boutons on PNs of different types (Figure 1M). We found that cognate boutons on PNs of the same type were placed much closer together than cognate boutons on PNs of different types (Figure 1N). Bouton position is the result of collateral placement and length. Therefore, we next compared collateral branch point location and height between cognate collaterals. While the extent of the peripheral spread of collaterals is correlated with their number, precise M-L location is best predicted by cell type: We found that the M-L distance between cognate collaterals on PNs of the same type was significantly less than that between cognate collaterals on PNs of a different type with the same number of collaterals (Figure 1O).

Similarly, cognate collaterals on PNs of the same type tend to be more similar in height than cognate collaterals on PNs of different types (Figure 1P). Overall similarity in axonal arbor shape across sister PNs of the same type was also observed in the FAFB dataset (Zheng et al., 2018).

While the Hemibrain represents just a single brain instance, we observe remarkable concordance of PN type behavior between the Hemibrain dataset and many previous light and EM analyses. This demonstrates a high degree of consistency and predictability in bouton placement in the calyx, such that boutons carrying information about different odors are distributed in a mixed but developmentally non-random manner across brains (H.-H. Lin et al., 2007; Tanaka et al., 2004; Zheng et al., 2018, 2022) (Figure S1C). Moreover, A-P bouton position is predicted by neuroblast origin and birth time and M-L bouton position by collateral number. We did not identify a developmental correlate of D-V bouton position, although this too is predictable by PN type. We note that the arborizations of these same neurons are also characteristic and predictable in their dendritic locations in the antennal lobe and in their second axonal target site in the lateral horn (Jefferis et al., 2007).

### Individual Kenyon cells innervate restricted regions of the calyx

We next sought to describe the calyx innervation patterns of Kenyon cells and the PN repertoires that they have access to. Of the seven developmental Kenyon cell types, those receiving olfactory input fall into three major anatomic classes based on their axonal lobe innervation patterns: γ, α’β’, and αβ. Four identical mushroom body neuroblasts per hemisphere produce each of the Kenyon cell types: γ Kenyon cells are born during the embryonic and early larval stages, α’β’ during late larval stages, and αβ during pupation (Figure 2A) (T. Lee et al., 1999). To examine how these developmental traits influence calyx organization, we ascertained the clonal origin of each Kenyon cell by measuring its axonal position outside the calyx, as it traverses the pedunculus on its way to the lobes (see Methods); for γ, α’β’, and αβ type assignment, we used Hemibrain annotations (Scheffer et al., 2020).

**Figure 2.**
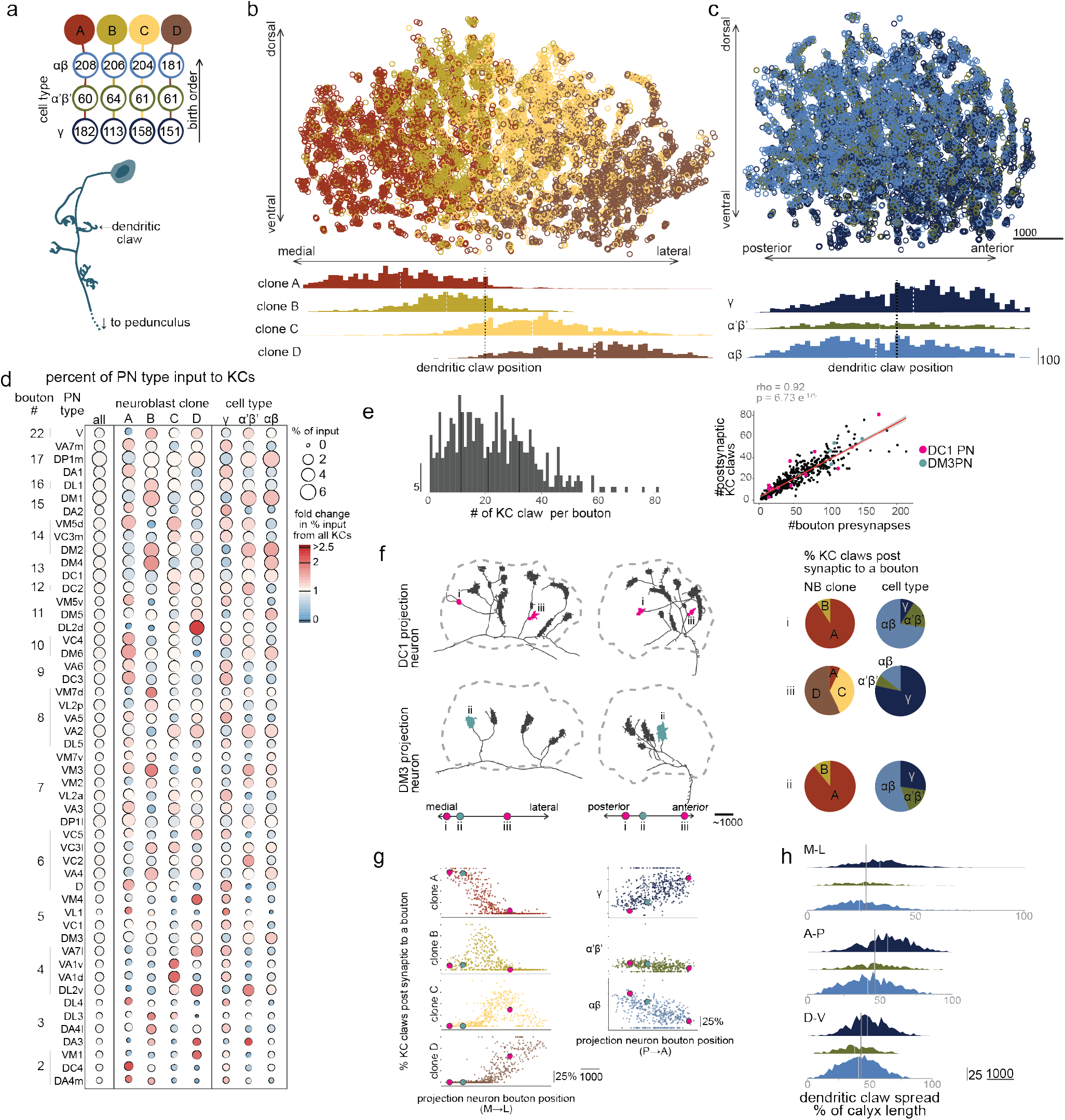
**(a)** top: Schematic of developmental history of Kenyon cells with numbers of olfactory KCs assigned to each neuroblast and type in the Hemibrain. Bottom: Anatomy of a KC in the calyx. **(b- c)** Centroid of KC claws (n=10,195) colored by the predicted neuroblast origin along the M-L axis (b) and colored by annotated KC type along the A-P axis(c). Histograms of claw center locations separated by NB origin or KC type is below with a bin width of 100 pixels. **(d)** Distribution of input from PNs of different types to different groups of Kenyon cells. Size of the circle represents the percent of claws of KCs in that group connected to PNs of each type. Each column sums to ∼100%. Color of the circle represents the fold change in percent representation from the value for all KCs (leftmost column). **(e)** left: histogram of the number of claws enwrapping each bouton (n=432) in the Hemibrain. Bin width is 1. right: Boutons with more presynaptic sites have more KC claws enwrapping them (Spearman’s rank correlation, rho=0.91, p=6.73e-175). DC1 and DM3 boutons are highlighted in pink and cyan respectively. **(f)** left: Hemibrain skeletons of a DC1 PN (top) and a DM3 PN (bottom) in the M-L (left) and A-P (right) views. Dotted line: calyx outline. Right: Pie charts of the clonal origin or type of KCs postsynaptic to each labeled bouton. Boutons i and iii are from the same PN, but located far apart in the calyx, while i and ii are from different cells, but located nearby. The boutons closer in space have more similar postsynaptic partner makeup than distant boutons of the same PN. **(g)** For each bouton, percent of KC claws from each of the 4 clones (left) and each of the three KC types (right). Boutons are organized by their M-L position (left) or their A-P position (right). Boutons labeled in 2f are shown in pink and cyan. **(h)** Histograms of the maximum distance between claws on a KC, along each axis, separated by type. Bin width is 100. All graphs are scaled to the same absolute spatial scale, with percent of that dimension labeled separately.

Consistent with previous light microscopy analyses, Kenyon cell dendrites in the adult calyx retain vestiges of clonal and birth order (type) segregation (H.-H. Lin et al., 2007; Zhu et al., 2003). The clones of Kenyon cells deriving from the four neuroblasts are arrayed across the calyx along the medial-lateral axis. We named these four clones A-D from medial to lateral. Kenyon cell primary neurites retain a rough concentric organization within each clone, with the primary neurites of early-born γ cells located at the periphery of each clonal unit and primary neurites of late-born αβ cells at the interior. Individual Kenyon cells extend 3-10 dendritic claws from the primary neurite within the calyx, each of which enwraps a single PN bouton (Caron et al., 2013; Leiss et al., 2009). We asked whether claw locations of a Kenyon cell are constrained by the position of its primary neurite. By plotting the location of each claw in the calyx, we found that claws from Kenyon cells sharing a neuroblast origin are clustered together along the medial-lateral axis, though claws from Kenyon cells derived from adjacent clones overlap and there are no sharp boundaries (Figure 2B). Claws from Kenyon cells of each of the three types are found throughout the calyx though their density varies across the anterior-posterior axis: claws from γ Kenyon cells tend to be in the more anterior part of the calyx while claws from the later born α’β’ and αβ Kenyon cells are denser in the posterior end of the calyx (Figure 2C).

Our analysis from Figure 1 predicts that different domains of the calyx will contain regionally unique mixtures of PN bouton types. To test this, we described the olfactory inputs to each Kenyon cell clonal or type group by calculating the percent of claws receiving input from each of the 51 olfactory projection neuron types. The proportion of overall input contributed by each PN type was heterogeneous and influenced by both the number of boutons of that type and the number of claws enwrapping them (Figure 2 D). In the Hemibrain, the average bouton is enwrapped by 22 claws (Range: 2-81, middle 50%: 11-32 claws). The exact number of claws postsynaptic to a bouton is highly correlated with the number of presynapses it contains (Figure 2E). The number of presynapses in a bouton varies across the calyx as well as across boutons of the same PN (Figure 2E, Figure S2).

All projection neuron types connect to Kenyon cells born from all four neuroblast clones in the Hemibrain (with the exception of the DL3 PN to clone D Kenyon cells) and to all Kenyon cell types. However, consistent with the PN and Kenyon cell type overlaps described in Lin 2007 and our analysis of the varying distribution of PN bouton types in Figure 1, Kenyon cells of different clonal origins and of different types received inputs from PN types at different rates. For example, in Lin 2007, DM2 PNs elaborate boutons in all four clones with a slight preference for the middle two, as is also the case here in Hemibrain (Figure 2D).

To disentangle spatial from molecular influences on connectivity, we compared the groups of Kenyon cells postsynaptic to different PN boutons. While each Kenyon cell claw grasps only a single PN bouton, each bouton is enwrapped by an average of 22 Kenyon cell claws. We compared the make-up of Kenyon cells postsynaptic to boutons that shared molecular similarities (from the same neuron) or spatial similarities (neighbors from different neurons). We found that the percentage of claws from Kenyon cells of each clone and type was variable across boutons of the same PN and that nearby boutons from different PNs were more similar to one another (Figure 2F). We plotted the percent of claws postsynaptic to each bouton from each clone and type by the bouton’s M-L or A-P location (Figure 2G). We found that across the calyx, the M-L location of a bouton predicted the percent of its postsynaptic partners from each of the four neuroblast clones, while the A-P position predicted the percent of its postsynaptic partners from each of the three types.

In summary, Kenyon cells make dendritic claws whose placement is dependent on their neuroblast origin and cell type. Because they are limited to a region of the calyx, they receive input from PNs at different rates. Connectivity between PNs and Kenyon cells is better predicted by the proximity of their neurites than by their cell types, as distant boutons from the same cell have different postsynaptic partner profiles. Kenyon cells are particularly restricted along the long M-L axis of the calyx, as dendrites of individual cells only reach a quarter of its extent (Figure 2H).

### Within a Kenyon cell’s restricted region of the calyx, its inputs are randomized

Previous analyses have debated the extent to which PN inputs to Kenyon cells are random or structured (Caron et al., 2013; Ellis et al., 2023; Ganguly et al., 2024; Hayashi et al., 2022; Jefferis et al., 2004; F. Li et al., 2020; Marin et al., 2002; Murthy et al., 2008; Tanaka et al., 2004; Wong et al., 2002; J.-Y. Yang et al., 2023; Zheng et al., 2022). Based on these previous reports, we hypothesized that Kenyon cells use a stochastic developmental mechanism to acquire their inputs, but because individual Kenyon cells are spatially restricted, the structure inherent in the placement of different PN boutons produces correlations in the patterns of inputs to Kenyon cells innervating different regions. To test this, we generated models of connectivity between Kenyon cells and projection neuron input types: We first started with a model similar to the “random bouton model” in Zheng 2022 in which each simulated Kenyon cell in the calyx randomly picked inputs, with the probability of selecting a given input type proportional to the number of PN boutons of that type in the Hemibrain data. Claw numbers for simulated Kenyon cells were kept consistent with their Hemibrain counterparts. To account for the effect of different spatial domains, we restricted this random bouton model to separately consider Kenyon cells from each of the four clones (medial to lateral spatial domains); each of the three cell types (anterior to posterior spatial domains); and the intersection of clonal origins and cell type (12 sub-domains). In these spatially restricted models, the probability of a modeled Kenyon cell selecting any PN input type was proportional to the number of boutons reachable by Kenyon cells of that type and/or clone in the Hemibrain data (Figure 3A).

**Figure 3.**
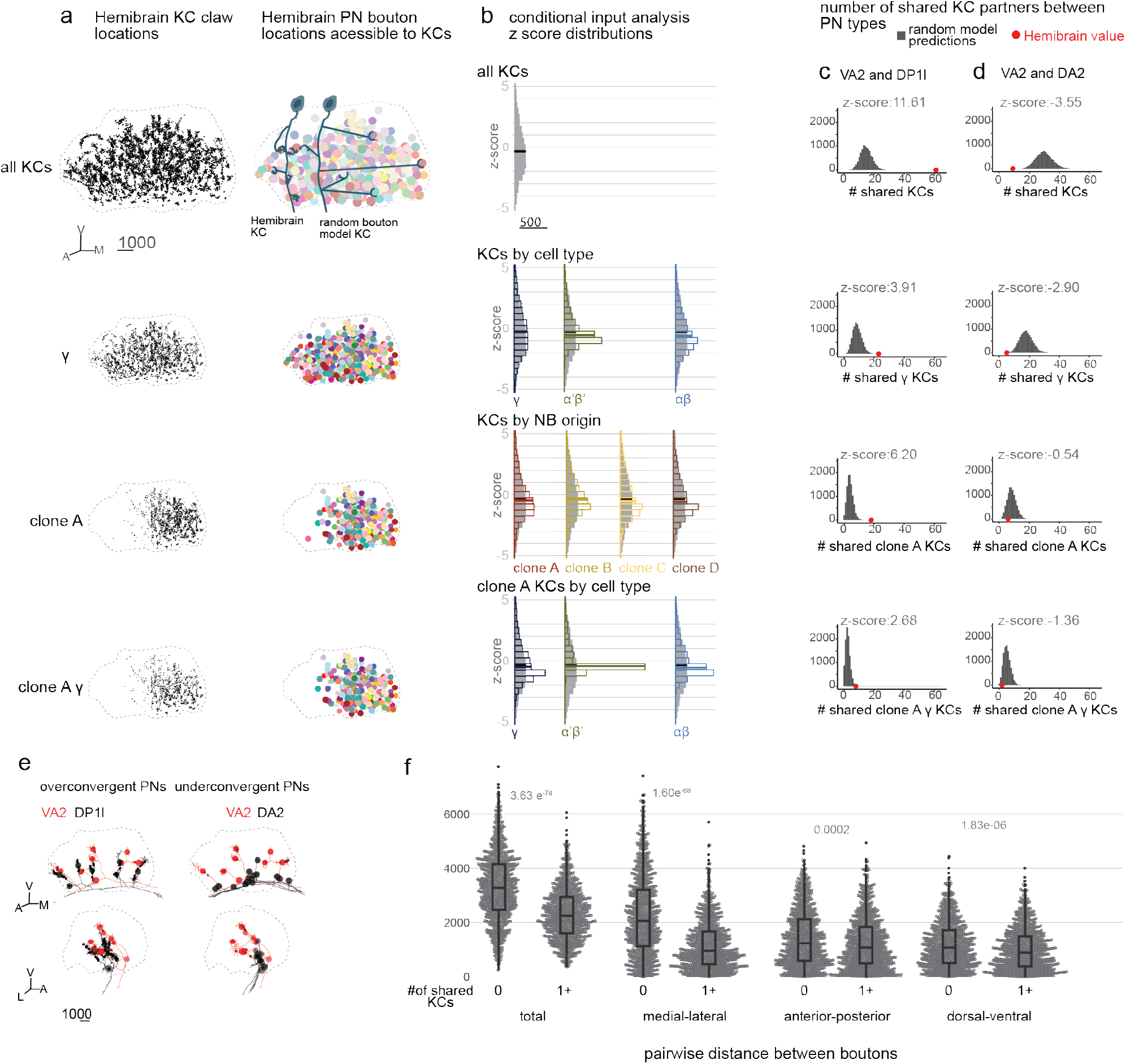
**(a)** left: centroids of claws of different KC groups in the Hemibrain calyx (dotted outline). Right: centroids of PN boutons accessible by KC groups in the Hemibrain calyx. Random bouton model KCs have the same number of claws as Hemibrain KCs but pick bouton inputs randomly from those reachable by any Hemibrain KCs of their type. **(b)** Histograms of conditional input analysis z scores for all PN types for each group of KCs. Conditional input z score is the number of standard deviations from the mean prediction of 10,000 random models that the probability of a KC receiving one PN type input given another observed in the Hemibrain. Bin width is 0.5. Thicker bars represent the median. **(c)** Number of KCs connected to DP1l as well as to VA2 PNs in the Hemibrain data (red dot) and predicted by 10,000 random models (grey bars) for models based on all KCs, γ KCs, clone A KCs, and clone A γ KCs. DP1l and VA2 are considered overconvergent as they share more KC partners than would be predicted by the all KC random model. **(d)** Number of KCs connected to DA2 as well as to VA2 PNs in the Hemibrain data (red dot) and predicted by 10,000 random models (grey bars) for models based on all KCs, γ KCs, clone A KCs, and clone A γ KCs. DA2 and VA2 are considered underconvergent as they share fewer KC partners than would be predicted by the all KC random model. **(e)** Hemibrain skeletons in the calyx of VA2 (red) with DP1l (black,left) or DA2 (black, right). Overconvergent pair VA2 and DP1l have boutons closer together than underconvergent pair VA2 and DA2. **(f)** Pairwise distances between boutons that share no (1,000 randomly sampled from 75,331) or at least 1 KC partner (1,000 randomly sampled from 19,499). Boutons which share at least 1 KC partner are on average 31% closer together (medians = 3297.77, 2264.38, Wilcoxon rank sum test, U =735269, p=3.63e^-74^, Cohen’s D=0.90). Boutons that share at least 1 KC partner are 53% closer together on the M-L axis than boutons who do not share KC partners and are slightly closer along the A-P (13%), and D-V (17%)axes than boutons who share no KC partners. (M-L: medians =2068.51, 962.51; Wilcoxon rank sum test, U=725899, p=1.60e^-68^; Cohen’s D=0.89. A-P: medians =1245.68, 1083.88; Wilcoxon rank sum test, U=548588, p=0.0002; Cohen’s D=0.19. D-V: Medians =1084.92, 899.61; Wilcoxon rank sum test, U=561616, p=1.83e^-06^; Cohen’s D=0.23). In box plots: center line, median; box limits, upper and lower quartiles; whiskers, 1.5x interquartile range; points, outliers.

For each model type, we generated 10,000 random connectivity matrices to compare to the connectivity observed in the Hemibrain using the conditional input analysis method described in Zheng 2022. This method calculates a z-score for each pair of PNs which describes how distinct the number of KCs which connect to both types in the actual data is from the numbers produced in the 10,000 randomly generated connectivity matrices. Using this method, observed connectivity in the Hemibrain calyx statistically diverged from that predicted by the full calyx random models. The pairs of PN types that most diverged in the observed data from the full calyx random model were a similar set of PN types that were reported as “over convergent” in Zheng 2022, which analyzed a different brain, the FAFB volume. These PNs correspond to the outer group of PNs in Tanaka 2004 and tend to have collaterals in more peripheral regions of the calyx, as we show in Figure 1J.

As we moved from modeling the whole Hemibrain calyx to Kenyon cells in spatially restricted subdomains, observed connectivity more closely recapitulated that predicted by random models (Figure 3B). To explicate the origin of this pattern, we picked the pair of PN types most over-convergent in the Hemibrain calyx compared to the full calyx models: VA2 and DP1l (Figure 3 C). 61 Hemibrain Kenyon cells receive inputs from both VA2 and DP1l, which significantly diverges from the average of ∼15 predicted by the whole calyx random model. For comparison, we selected a third PN type, DA2, which is least frequently co-sampled with VA2 (Figure 3D,E). Only 9 Hemibrain Kenyon cells receive inputs from both VA2 and DA2, in contrast to the average of 30 KCs predicted by the whole calyx random model. While the observed co-sampling of these pairs differs dramatically from the prediction based on the full calyx models, it differs minimally from our spatial domain-based models.

We next investigated the spatial relationships between boutons presynaptic to the same Kenyon cell. Remarkably, we found that pairs of boutons co-innervating at least one Kenyon cell were closer on the medial-lateral axis than boutons not sharing Kenyon cells, but had minimal unique proximity along the dorsal-ventral or anterior-posterior axes (Figure 3F). This suggests that the spatially constrained dendritic development of Kenyon cells from different neuroblast clones is the primary determinant of correlation structure in PN inputs to individual cells. While Zheng et al 2022, and Li et al 2020 found that imposing spherical spatial constraints on random models of connectivity improved model performance, our model identifies a single spatial axis that constrains KC choices and that is produced by a key feature of Kenyon cell development: neuroblast origin.

Combining the structured patterning of PN bouton types in the calyx with random input acquisition by Kenyon cells restricted along the M-L axis can explain all the anatomic and functional observations to date: precise combinations of KC inputs are unpredictable, but correlations are observed in PN sampling by Kenyon cells distributed in different calyx regions.

### Kenyon cells express a reduced set of cell surface molecules

Our anatomical observations appear to be in conflict: from the projection neuron point of view, the calyx is stereotyped and orderly. On the other hand, while Kenyon cells have their own spatial constraints, they don’t appear to discriminate among the PNs they encounter. Different PN types express numerous and diverse cell surface molecules (Hongjie Li et al., 2017; Xie et al., 2021). While some PN-type-specific factors could be trafficked exclusively to antennal lobe dendrites to govern glomerular specificity, PNs also make precise and stereotyped connections onto downstream partners in the lateral horn (Marin et al., 2002; Wong et al., 2002). Kenyon cells are thus likely faced with cell surface molecular diversity on the presynaptic processes of potential PN partners. This led us to ask what tools they might have on their cell surface to help them acquire varied PN inputs.

To answer this, we analyzed Kenyon cell transcription across developmental time (Figure 4A). First, we performed RNA-seq on bulk FACS-sorted projection neurons and Kenyon cells at 45-46h APF (after puparium formation) (one replicate) and in adulthood (2-7 days after eclosion, two replicates) (Figure 4B). We also sorted and sequenced cells that were neither PNs nor Kenyon cells (“double negatives”) for comparison. We characterized the expression of 932 putative cell surface and secreted (CSS) molecules compiled by the Zinn lab based on protein domain architectures (Kurusu et al., 2008). In projection neurons, similar numbers of CSS’s were up- and down-regulated compared to double negative cells at both time points, as expected (Figure 4B). In contrast, most CSS’s were downregulated in Kenyon cells compared to double negatives at both time points (Figure 4B). The downregulation could reflect either global reduction of these transcripts or selective expression of transcripts in a subset of Kenyon cells.

**Figure 4.**
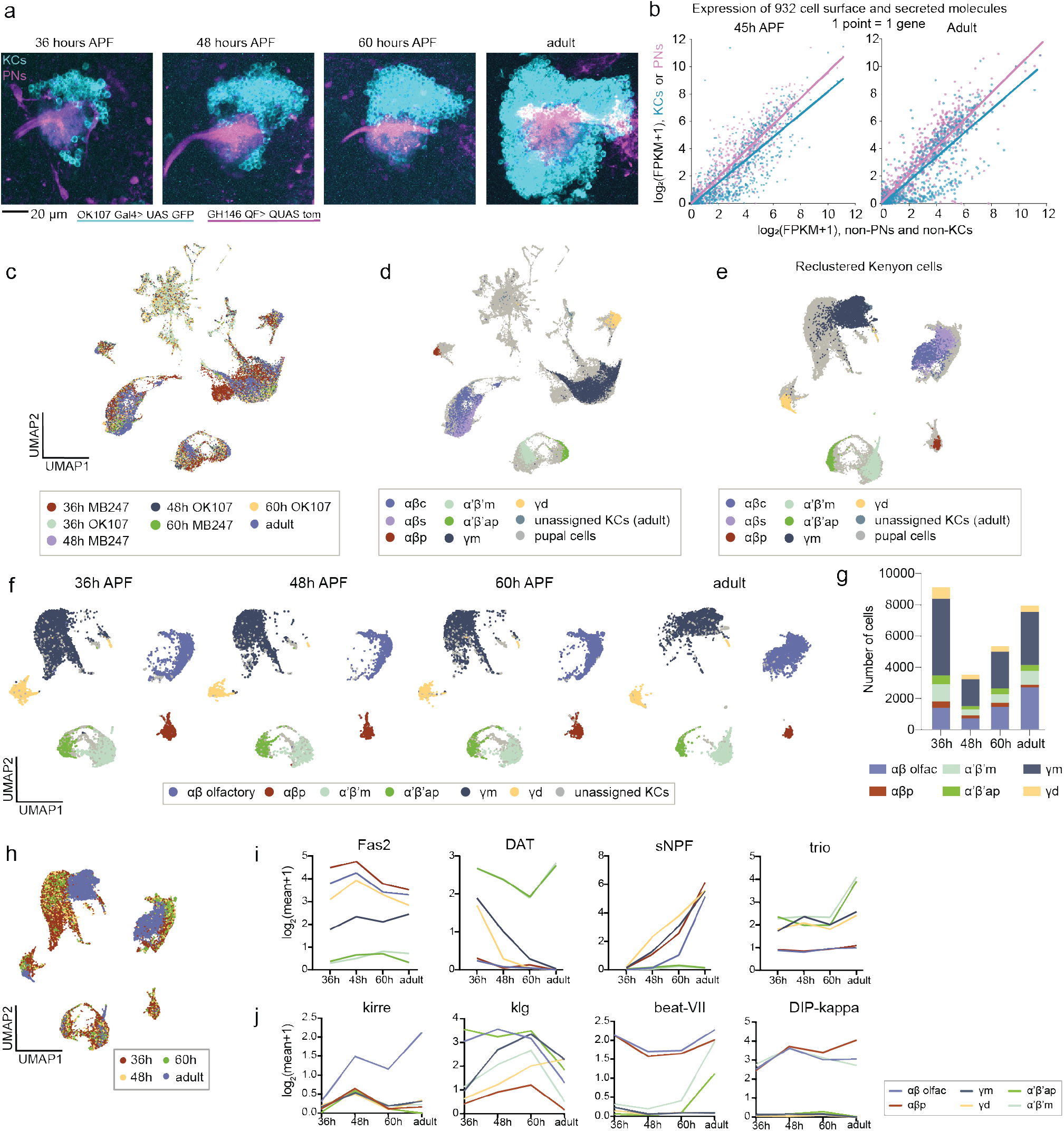
**(a)** KCs (cyan, OK107-Gal4) and PNs (magenta, GH146-QF) innervating the calyx at 36hAPF, 48hAPF, 60hAPF, and 2-3 day old adults. **(b)** Expression of 932 cell surface and secreted (CSS) molecules in pooled 45h APF and adult KCs or PNs plotted against expression in unlabeled (non- PN, non-KC) brain cells. Each point is one CSS gene. Lines represent linear regression of KC data (cyan) and PN data (pink). The downward shift of the KC regression line shows that there is a lower expression of CSS genes KCs relative to non-KCs, and non-PNs. **(c)** UMAP of scRNAseq data from Kenyon cells at all time points (36hAPF, 48hAPF, 60hAPF, adult) along with the driver lines used to FAC-sort them (MB247 or OK107). **(d)** UMAP showing adult pre-assigned KC identities mapped onto the data shown in c. Adult data is from multiple split-Gal4 lines labeling the 7 Kenyon cell classes. KC class-assigned adult cells are labeled in different colors. “Pupal cells” refers to cells from the 36h, 48h, and 60h timepoints. The clusters that do not match any of the adult KC identities are the non-Kenyon cells in the dataset, which were sorted as false positives.**(e)** UMAP of Kenyon cells subset from the data in c, d. Colors and corresponding identities of clusters are shown from the adult data. **(f)** Distribution of KC classes across the different stages. This is the same UMAP in e, split by the developmental stage. Cell types were assigned by matching to adult KC data, and by verifying expression of known markers (a subset of these are shown in panel i). **(g)** Number of Kenyon cells belonging to each class for each time point.**(h)** UMAP of KCs from 36h, 48h, 60h and adult where KCs from each stage are labeled a different color. **(i-j)** Log-normalized average expression of known KC type marker genes (i) and pan-KC differentially expressed cell adhesion molecules (j) at the different time points. x axes show the stages, and colors indicate the KC class. Each line is the normalized average expression of that gene in that KC class at the 4 time points. (i) Based on adult Kenyon cell data in the literature, Fas2 is highest in αβ, lower in γ, and not reported to be expressed in α’β’ KCs. DAT is specific to α’β’ KCs. γ KCs express trio and sNPF, α’β’ KCs express trio and DAT, and αβ KCs express sNPF. (j) Kirre is highest at 48h. For most KC types, klg expression increases at 60h, which corresponds to the stage of synaptogenesis. Beat-VII expression increases from 60h to adult, and DIP-kappa is specific to αβ and α’β’ and changes over time in these cell types but is not expressed in γ KCs.

We reasoned that investigating transcriptional patterns in individual cells could point to cellular mechanisms of partner choice: If individual Kenyon cells express stochastic combinations of cell surface molecules, they could have the potential to actively choose inputs from distinct projection neuron partners. If individual Kenyon cells weakly express cell surface molecules, they might acquire inputs promiscuously because they lack the molecular tools to tell those inputs apart. We therefore performed single-cell RNA sequencing (scRNAseq) on Kenyon cells across pupal development. We sequenced Kenyon cells at three different time points: 48h APF, when PN:KC contacts first appear; 60h APF, during a wave of synaptogenesis across the central brain (Muthukumar et al., 2014); and 36h APF, to capture any important transcriptional changes that may occur prior to initial contacts. We also sequenced adult Kenyon cells in which the seven developmental types were labeled *a priori* (see Methods).

Transcriptomic clusters at all three developmental time points aligned well with the cell types assigned in the adult data and expressed known type-specific markers (Croset et al., 2018; G. Lee et al., 2000; Shih et al., 2019). Pupal Kenyon cells clustered into 6 major groups with the following identities, listed in order of their birth: γd, γm, α’β’m, α’β’ap, αβp, αβ (Figure 4C-H)(T. Lee et al., 1999). (Pupal αβ cells did not cluster neatly onto earlier-born “shell” and later-born “core” subtypes that are observed in the adult because Kenyon cell neurogenesis continues until eclosion, with αβ core cells born only at late pupal stages.) Two Kenyon cell subsets, γd and αβp, innervate accessory calyx regions and receive visual, rather than olfactory, inputs (Aso et al., 2009; Ganguly et al., 2024; F. Li et al., 2020; Vogt et al., 2016). These visual Kenyon cell subtypes clustered separately from olfactory γ and αβ Kenyon cells at all time points (Figure 4D-F). We have previously shown that visual γd Kenyon cells cannot support olfactory PN bouton formation in the calyx, and that individual γd cells receive unpredictable combinations of visual inputs from specific visual projection neuron types innervating diverse optic lobe neuropils (Elkahlah et al., 2020; Ganguly et al., 2024).

We were able to identify markers that consistently marked Kenyon cell subtypes throughout development (e.g. sNPF, Fas2, DAT, trio, Figure 4I). We also identified a set of genes with dynamic expression across Kenyon cell subtypes, including the Immunoglobulin superfamily (Ig-SF) molecules kirrel, beat-VII, and DIP-kappa (Figure 4J). While genes whose expression varies across developmental time in all Kenyon cells are unlikely to contribute to their acquisition of varied projection neuron inputs, these genes could be used globally by all Kenyon cells at important stages of synapse formation with PNs generally.

Having identified the Kenyon cell classes at each time point, we then compared expression of cell surface and secreted molecules (CSS’s) in our bulk RNAseq data to scRNAseq data, and cross-checked these expression patterns with MIMIC-Gal4 reporters for Ig-SF protein families called dprs (defective proboscis response) and their binding partners, DIPs (dpr-interacting proteins). The reporter expression, bulk RNAseq and single-cell RNA-seq expression matched well, with certain DIPs and dprs expressed ubiquitously in all Kenyon cells while others were expressed in a subset of Kenyon cells (Supplementary Figure 4). Isolated Kenyon cells expressing a particular Ig- SF gene could be Kenyon cells of a specific developmental type.

To test if the bulk depletion of CSS transcripts in Kenyon cells (Figure 4B) reflects heterogeneous expression across cells or low expression in each cell, we investigated CSS expression in individual Kenyon cells by summing up the normalized numbers of transcripts (i.e. ln(TP10K+1)) of all CSS genes expressed in each cell. We found that there were less CSS transcripts overall in individual visual and olfactory Kenyon cells relative to non-Kenyon cells (Figure 5A). This analysis supports the model that cell surface molecules are depleted in individual Kenyon cells.

**Figure 5.**
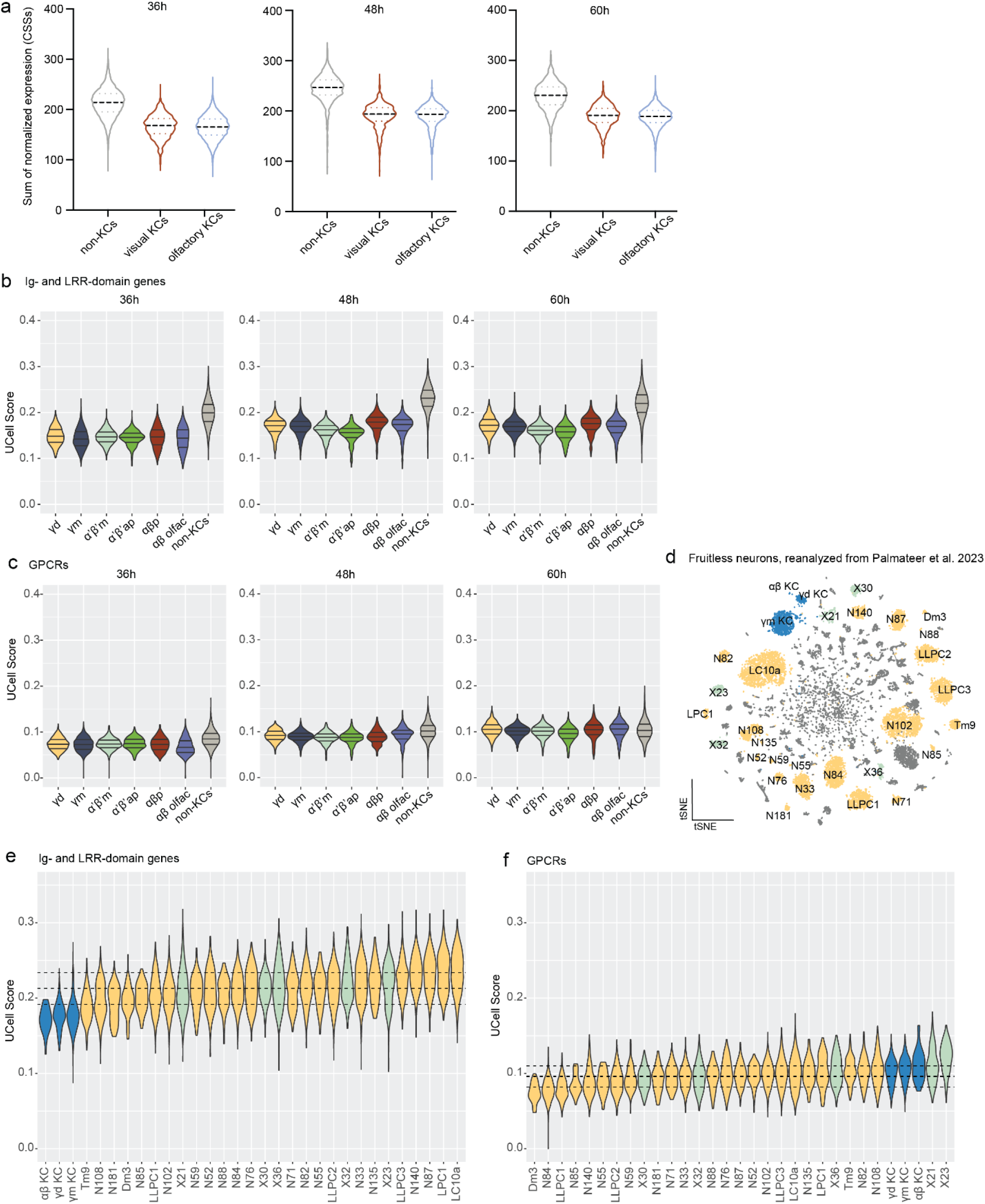
**(a)** Summed expression of 932 cell surface and secreted molecules in individual Kenyon cells and non-Kenyon cells sorted as false positives. Violins show distributions of values for individual cells. Black dashed line: median, colored dotted lines: 25% and 75% quartiles. **(b,c)** Gene set enrichment scores (UCell Scores) for Ig- and LRR-superfamilies of CSS (b) and GPCRs (c). Violins show distributions of UCell values for individual cells. Solid lines: 25%, 50% (median), and 75% quartiles for each violin. N for each category shown in Figure 4g. Additional gene sets are shown in Supplementary Figure 5-1. **(d)** tSNE of *fruitless*-expressing neurons at 48h APF from Palmateer et al. Labeled clusters are shown in (e,f). Blue: Kenyon cells. Yellow: optic lobe types matched to Yoo et al. Transcriptional types with known anatomical matches in connectomes are named, transcriptional types that are not yet matched to cell types in connectomes are tagged with “N.” Green: selected types from central brain, tagged as “X.” Additional analyses shown in Supplementary Figures 5-2 to 5-4. **(e,f)** UCell scores of Ig-SF+LRR and GPCR gene sets for select cell types and clusters from (d). Violins show distributions of UCell values for individual cells. Dashed lines: 25%, 50% (median), and 75% quartiles across the entire dataset . Clusters are ordered by UCell median. Additional gene sets are shown in Supplementary Figure 5-5.

Next, we narrowed down the CSS list to focus on immunoglobulin (Ig) and leucine-rich repeat (LRR) domain-containing molecules. Genes in these families are expressed in diverse and specific patterns across different types of neurons in the fly brain and engage in complex biochemical interactions (Barish et al., 2018; Brovero et al., 2021; Carrillo et al., 2015; Kurmangaliyev et al., 2020; Hanqing Li et al., 2017; Hongjie Li et al., 2017; Özkan et al., 2013; Tan et al., 2015; Yoo et al., 2023). They are thus predicted to regulate synaptic partner choice or other type-specific processes during neuronal development (Hassan & Hiesinger, 2015; Hiesinger, 2021; Sanes & Zipursky, 2020). We quantified the enrichment of these specificity molecules in each cell using UCell, a gene ranking-based approach for quantifying gene signature scores (Andreatta & Carmona, 2023). These specificity molecules were strikingly low-ranked in Kenyon cells of all types compared to non-Kenyon cells, and this relationship was true across time points (Figure 5B). We note that Kenyon cells are small cells with lower amounts of RNA per cell than other neurons in the central brain; in accordance with this, we obtain fewer UMI’s per Kenyon cell than for other cells in the central brain (Supplementary Figures 5-1). The rank-based computation used by UCell successfully accounts for this difference in overall RNA content. For example, UCell enrichment of transcription factors and GPCRs was similar between Kenyon cells and non-Kenyon cells (Figure 5C, Supplementary Figure 5-1). Reduced expression of specificity molecules in Kenyon cells was driven by Immuoglobulin superfamily (Ig-SF) proteins, while expression of LRR-domain transcripts was more consistent with that in non-Kenyon cells (Supplementary Figure 5-1).

The non-Kenyon cells in our datasets represent a random and likely very heterogeneous sample of various cell types. Therefore, we decided to repeat our analysis in other datasets that include Kenyon cells and other distinct, well-defined neuronal cell types. The Arbeitman lab has recently published a scRNAseq dataset that covers *fruitless*-expressing neurons, including a few Kenyon cell types (Palmateer et al., 2023). The initial analysis of this dataset revealed many transcriptionally distinct populations of neurons and mapped select gene expression programs to morphologically defined cell types (Palmateer et al., 2023). Most cell types in the central brain are ultra-rare and unlikely to be resolved into separate clusters at the given resolution of the dataset (Scheffer et al., 2020). The most abundant *fruitless* cell types include γ Kenyon cells and cell types in the visual system, which are represented by dozens to hundreds of copies per brain. Transcriptomes of many visual system neurons have been well characterized in the comprehensive transcriptional atlases of the *Drosophila* optic lobes (Kurmangaliyev et al., 2020; Özel et al., 2021).

We re-analyzed this *fruitless* dataset and mapped the identities of Kenyon cells and visual system neurons (Figure 5D, see Methods for details and Supplementary Figures 5-2, 5-3, 5-4). In total, we identified 132 clusters of neurons, including three types of Kenyon cells and 23 clusters of visual neurons previously identified by transcriptomics (7 of these are matched to morphologically defined connectomic cell types). We repeated the enrichment analysis of Ig and LRR domain molecules for *fruitless* neurons (Figure 5E-F, Supplementary Figure 5-5). All three types of Kenyon cells had significantly less of these specificity factors than all other *fruitless* clusters, including the 23 cell types from the visual system. Again, no differences were observed for transcription factors or GPCRs (Supplementary Figure 5-5). Together, enrichment analyses of individual Kenyon cells from both our data and from the *fruitless* dataset support a model in which each Kenyon cell is generally depleted of Ig-SF proteins, rather than different cells robustly expressing different family members.

Interestingly, among the Ig-SF molecules depleted in KCs is the entire family of Beats (Supplementary Figures 5-2, 5-4). 14 Beats form complex heterophilic receptor-ligand networks with eight Sides (Hanqing Li et al., 2017). Two pairs of Beats and Sides have been recently shown to be involved in synaptic target choice in the visual circuits (Osaka et al., 2024; Yoo et al., 2023). In both cases, Beats were expressed by postsynaptic neurons. We speculate that the depletion of Beats and other Ig-SF molecules can lead to increased promiscuity at postsynaptic sites of Kenyon cells.

Essentially, projection neurons of different types express different combinations of surface molecules, but if Kenyon cells lack the proteins that would bind these molecules, they may not be able to select specific PN partners.

Of course, Kenyon cells do not lack partner specificity entirely: Kenyon cells of different types receive input from precise sensory modalities (Ganguly et al., 2024; F. Li et al., 2020; Yagi et al., 2016), and indeed some Ig-SF molecules are differentially expressed in olfactory versus visual Kenyon cells, e.g. dpr1 is expressed higher in γd (visual) compared to γm (olfactory) Kenyon cells (Supplementary Figure 5-2). Olfactory Kenyon cell subtypes innervate distinct axonal lobes and connect with lobe-specific dopaminergic neurons and mushroom body output neurons. As described previously, DIP and dpr Ig-SF proteins are required for these processes (Bornstein et al., 2021; Morano et al., 2024). Consistent with this, we find that certain cell surface genes are selective for distinct Kenyon cell olfactory subtypes, e.g. Lac was selectively expressed in α’β’m Kenyon cells (Supplementary Figure 5-2). As Kenyon cells of every olfactory subtype can receive inputs from every PN type (Figure 2) (Caron et al., 2013), we do not expect differences in transcription *between* olfactory Kenyon cell types to explain the diversity of inputs from PNs.

### When deprived of typical partners, Kenyon cells still synapse

Our connectomic and transcriptomic analyses suggest that Kenyon cells partner randomly with nearby PN boutons rather than using cell surface molecules to structure connectivity. We sought to test Kenyon cell promiscuity *in vivo* by changing the composition of PN boutons. To accomplish this, we killed half the PNs during development by driving expression of diptheria toxin CBβ with VT033008-Gal4 which, as we described previously, labels a defined PN subset which includes most of the PNs from the anterodorsal lineage (Elkahlah et al., 2020) (Figure 6A,B). This manipulation produced a shift in the PN repertoire represented across boutons. VT033008+ PNs contribute an average of 234 boutons in control and are lost when these neurons are ablated (Figure 6C-E). Moreover, remaining PNs, labeled by VT033006-LexA driver and/or ChAT immunostaining, increase their bouton production, such that ablated calyces lack only ∼170 boutons on average (Figure 6E). We next decided to look at the behavior of smaller subsets of PNs in the context of VT033008 PN ablation. We labeled ∼12 PNs from 3 glomeruli labeled by MZ19 QF (Figure 6F-I). In the ablation condition, an average of 2 MZ19 PNs were lost, but we observed a 60% increase in MZ19 boutons in the calyx (Figure 6G,I). These analyses show that in addition to loss of VT033008-labeled types, this condition also involved PN-type-specific rebalancing of calyx composition. For example, MZ19+ boutons go from contributing 6% of boutons in wild type to 14% of boutons in the ablation condition, and while the absolute number of boutons not labeled by VT033008 does not change significantly, these go from 55% to 100%.

**Figure 6.**
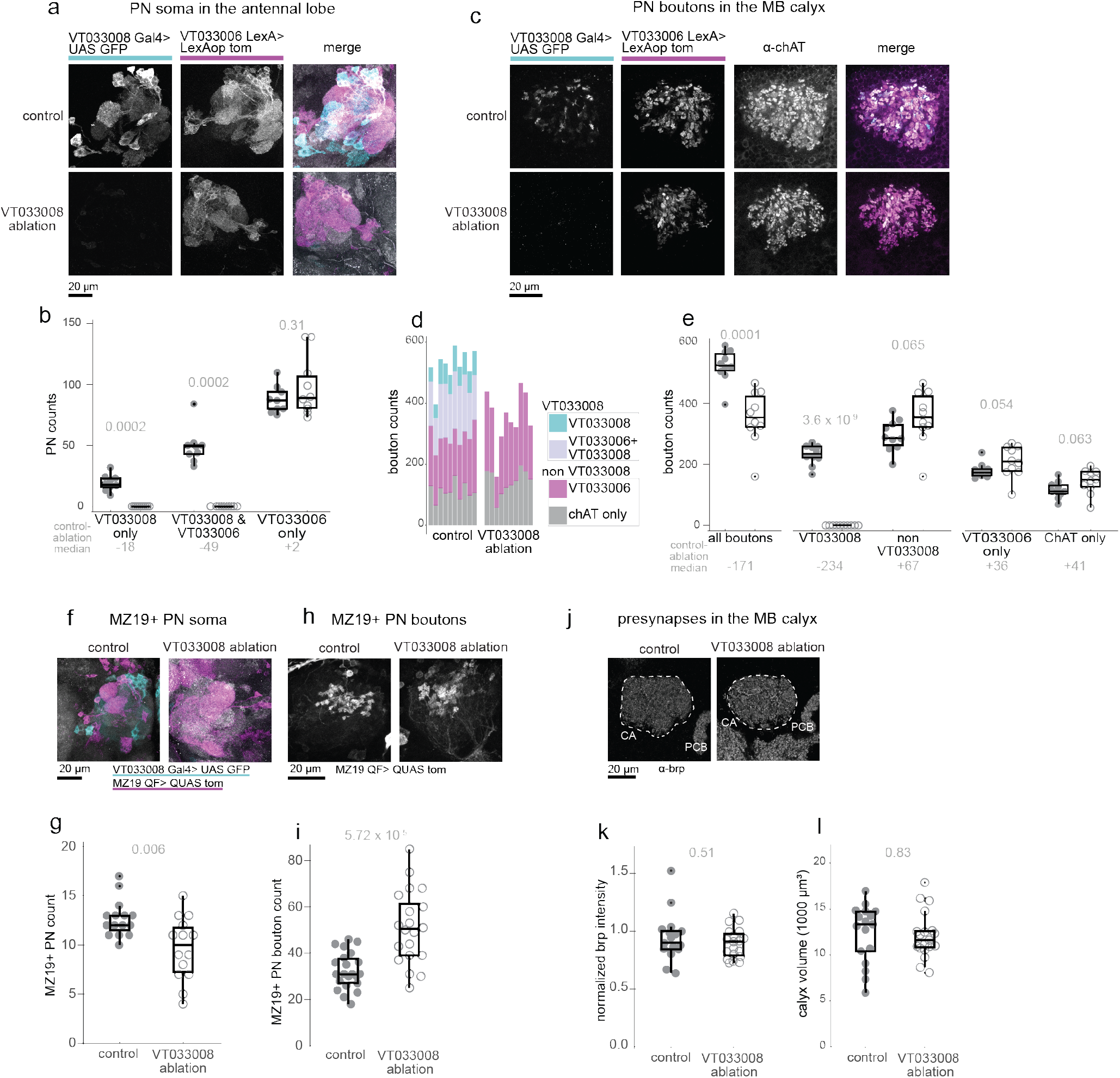
**(a)** Antennal lobes in control animals and animals subjected to ablation of half of PNs via VT033008-Gal4 driven expression of diphtheria toxin, shown in maximum intensity Z projection of confocal image. Cyan: VT033008+ PNs; magenta: VT033006+ PNs; grey: Immunostaining against choline acetyltransferase (ChAT) which marks all uniglomerular PNs. **(b)** Cell counts for PNs labeled by VT033008, VT033006, or both in control (9 hemispheres from 5 brains) or VT033008 ablation (10 hemispheres from 5 brains). Throughout this figure, grey bars depict medians and P values from Wilcoxon rank sum or t-tests are written above. The difference between control and VT033008 ablation median cell counts is written in grey below the bouton type label on the x axis. Here and throughout control hemisphere values are represented with filled dots on the left and VT033008 ablation values are open dots on the right. (VT033008 only: Wilcoxon rank sum test, U=95, p=0.00023; Cohen’s D=2.78.VT033006 & VT033008: Wilcoxon rank sum test, U=95, p=0.0023; Cohen’s D=3.02. VT033006 only: Wilcoxon rank sum test, U=36, p=0.31; Cohen’s D=0.69.) In box plots: center line, median; box limits, upper and lower quartiles; whiskers, 1.5x interquartile range; points, outliers. **(c)** Control and VT033008 Ablation calyces, single confocal slice shown. Cyan: VT033008+ PNs; magenta: VT033006+ PNs; grey: ChAT. **(d)** Distribution of boutons of each type for each of the 10 calyces quantified per condition. **(e)** Bouton counts separated by type (10 hemispheres from 5 brains each for control and VT033008 ablation). The difference between control and VT033008 ablation median bouton counts is written in grey below the bouton type label on the x axis. Saturating ablation of VT033008 cells reduces the number of boutons by a median of only 171, though 234 are contributed by cells expressing this driver in controls. The difference can be explained by an increase in boutons labeled only by VT033006 (median gain of 36 boutons) and those labeled by ChAT alone (median gain of 41 boutons).(all boutons: Student’s T-test, t=5.15, p=0.0001; Cohen’s D=2.3. VT033008 boutons: Student’s T -test, t=22.21, p=3.60e-09; Cohen’s D=9.93. Non VT033008 boutons: Student’s T -test, t=-2.04, p=0.06; Cohen’s D=0.91.VT033006 only boutons: Wilcoxon rank sum test, U=24, p=0.05; Cohen’s D=0.74.ChAT only boutons: Student’s T -test, t=-2.00, p=0.06; Cohen’s D=0.90.) **(f)** Control and VT033008-Ablated Antennal Lobes, maximum intensity confocal Z projection. Cyan: VT033008+ PNs; magenta: MZ19+ PNs; grey: ChAT. **(g)** Cell counts for PNs labeled by MZ19 in control (15 hemispheres from 11 brains) and ablation (14 hemispheres from 9 brains). (Student’s T -test, t = 3.08, p =0.006; Cohen’s D = 1.15) **(h)** Control and VT033008-Ablated calyces, single confocal slice of MZ19+ boutons. **(i)** MZ19+ bouton counts in control (22 hemispheres from 14 brains) and ablation (20 hemispheres from 11 brains). (Student’s T -test, t = -4.75, p =5.72e-05; Cohen’s D = 1.49) **(j)** Control and VT033008 ablation calyces labeled with an antibody against presynaptic marker bruchpilot (brp). Calyx (CA) and Protocerebral Bridge (PCB) are labeled. The PCB is unmanipulated region and used for normalization. **(k)** Normalized Brp intensity for control (16 hemispheres from 11 brains) and ablation (19 hemispheres from 11 brains) calyces. Normalized intensity was calculated as follows- mean gray value for a 250×250 pixel region of the calyx divided by mean grey value for 75×75 pixel region of the protocerebral bridge. (Wilcoxon rank sum test, U=183, p = 0.51; Cohen’s D = 0.27) **(l)** Calyx volume measurements from control (16 hemispheres from 11 brains) and ablation (19 hemispheres from 11 brains) calyces. (Student’s T-test, t = 0.22, p = 0.83; Cohen’s D = 0.07)

Even so, Kenyon cells appear able to obtain a normal number of inputs in this condition, suggesting that PN subtype identity does not restrict partnerships: We have previously reported that Kenyon cell number is unchanged by the loss of VT033008 PNs, and that even extreme olfactory PN loss does not affect Kenyon cell claw number (Elkahlah et al., 2020). Here, we find that overall calyx volume as well as the intensity of labeling of the presynaptic marker Bruchpilot do not change on loss of VT033008 PNs (Figure 6 J,K,L). As individual boutons can have widely varying numbers of synapses and connect with widely varying numbers of Kenyon cells (Figure 2E), our data is consistent with a model in which spared PNs modestly increase their bouton number while also increasing the number of synapses on each bouton and connected claws to preserve the overall synaptic repertoire of the calyx.

### Ectopic Ig-SF expression redistributes Kenyon cell claws onto boutons of PNs expressing cognate molecules

Previous work on the role of Ig-SF molecules in the development of connectivity, including our own, suggested that when these proteins are experimentally depleted, neurons synapse promiscuously (C. Xu et al., 2019; Yoo et al., 2023). Our analyses here suggest that Kenyon cells accept partners promiscuously due to their natural depletion of Ig-SF proteins. We thus sought to perform the opposite experiment: To ask if adding an Ig-SF protein to Kenyon cells could bias their partner choices. We identified the Ig-SF repertoires of a set of PNs that we could reliably label, using the genetic marker MZ19 (DA1, DC3, and VA1d) (Figure 7A, B). MZ19+ PNs express *DIP-eta* and *DIP-theta* (Figure 7 B). We examined expression of the partner molecules of these DIPs in Kenyon cells, and found that Kenyon cells expressed little to none of their partner, *dpr4* (Supplementary Figure 5-2)(Özkan et al., 2013).

**Figure 7.**
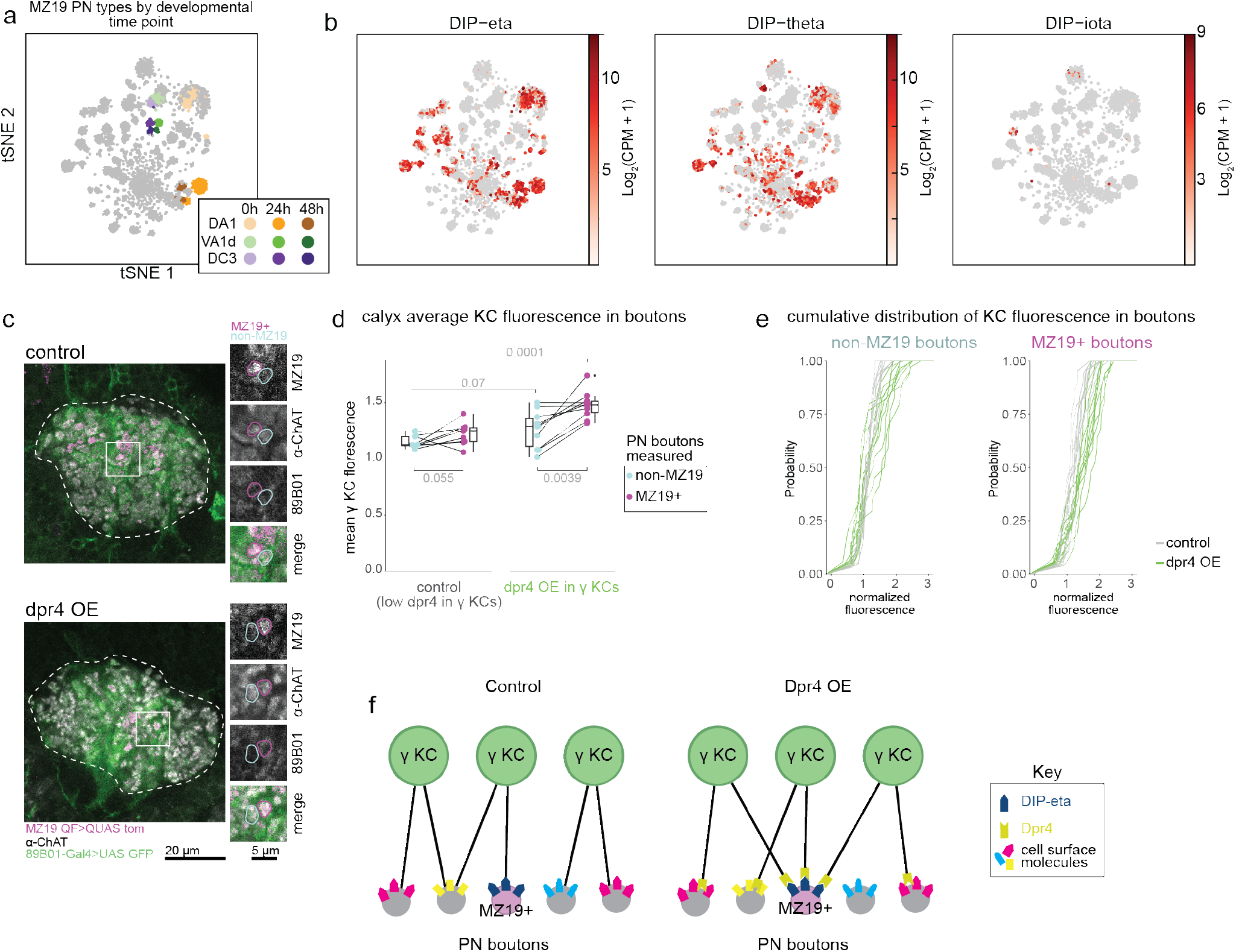
**(a)** tSNE plots of single cell RNA sequencing of projection neurons throughout development done by Xie and colleagues(Xie et al., 2021). Clusters of cells belonging to PN types labeled by genetic driver MZ19 are colored according to their type and developmental stage.**(b)** tSNE plots of log2 normalized expression of dpr4 partners Dip-eta, DIP-theta, and DIP-iota in projections neurons. These partners, particularly DIP-eta and DIP-theta are highly and relatively specifically expressed in DA1 PNs throughout development. **(c)** Control (top) and Dpr4 OE (bottom) calyces single z slices. Immunostaining against chat is in grey and shows PN boutons, MZ19-QF labeled boutons are in magenta, γ KCs labeled by 89B01-Gal4 are shown in green. The dotted outline circles the entire calyx and the box highlights a region with individual boutons which is magnified and shown in all channels to the right. Pink outlined bouton was also labeled by MZ19-QF. Cyan outlined bouton was not labeled by MZ19-QF. **(d)** Average normalized fluorescence values for non-MZ19 (cyan) and MZ19+ (pink) boutons of each calyx for the control (9 hemispheres from 6 brains) and dpr4 overexpression (10 hemispheres from 6 brains) samples. Average fluorescence values from the same calyx are connected by a line. Normalized KC fluorescence values are only slightly different between non-MZ19 and MZ19+ boutons in control calyces (Wilcoxon signed rank exact test for paired samples, U = 39, p = 0.055). MZ19+ boutons have higher normalized KC fluorescence values than non-MZ19 boutons in dpr4OE calyces (Wilcoxon signed rank exact test for paired samples, U = 54, p = 0.0039). MZ19+ boutons in calyces where dpr4 is overexpressed by γ KCs have significantly higher normalized KC fluorescence than MZ19+ boutons in control calyces (Student’s T-test, t = 5.02, p = 0.0001; Cohen’s D = 2.30). Normalized KC fluorescence in non MZ19 boutons is not significantly different in calyces where dpr4 is overexpressed compared to control (Student’s T-test, t = 2.02, p = 0.07, Cohen’s D = 0.91). In box plots: center line, median; box limits, upper and lower quartiles; whiskers, 1.5x interquartile range; points, outliers. **(e)** Cumulative density plot showing the normalized KC fluorescence values by calyx for MZ19+ (left) and non-MZ19 boutons in dpr4 overexpression and control conditions. **(f)** Model for overexpression experiment results. In control conditions, γ Kenyon cells do not express dpr4 and select inputs from nearby olfactory projection neurons. When γ Kenyon cells overexpress dpr4 they are able to recognize partner molecules like DIP-eta on PN partner boutons and select these over other nearby boutons.

We drove expression of the missing partner molecule in a subset of Kenyon cells and asked whether this could influence their PN selection. We chose to target the broadly-ramifying γ-Kenyon cells to nullify spatial limitations that we highlight above. We found that overlap between γ Kenyon cell claws and MZ19 PN boutons increased when γ Kenyon cells ectopically expressed dpr4 (Figure 7 C-E). We compared γ Kenyon cell claw fluorescence on MZ19 boutons to neighboring non-MZ19 boutons, which are less likely to express *DIP-eta* or *DIP-theta* compared to MZ19 cells (Figure 7B-E). Increases in γ Kenyon cell fluorescence on these boutons was modest compared to on MZ19 boutons, with the fluorescence intensity distribution of overexpression and control calyces intermingled. Together, these results suggest that while Kenyon cells in wild type animals cannot sense molecular differences between PN types, PN type selectivity can be induced if Kenyon cells are given the proper molecular tools (Figure 7F).

## Discussion

In 1958, Frank Rosenblatt sought to “understand the capability of higher organisms for perceptual recognition, generalization, recall, and thinking.” He formulated a model learning circuit, the Perceptron, which bears an uncanny resemblance to the mushroom body. Rosenblatt assumed that the Kenyon-cell-like “A units” of the Perceptron would receive random input combinations from the sensory apparatus (Rosenblatt, 1958). Though modern neural network AI has diverged from its biological inspiration in the brain’s *actual* intelligence, the utility (or simply sufficiency) of randomized initial conditions remains a key feature of the computational method. Of course, we now know that there is tremendous specificity in the distribution of neuronal cell types and connections in many parts of the brain, and while Rosenblatt assumed that inputs to neurons engaged in associative learning were randomized, he lacked the data to do so. Whether the inputs to biological learning circuits are in fact randomly wired has therefore been the subject of much debate. The question actually contains two different principles within it: (1) Whether randomized connectivity can be found in the adult brain and be sufficient for perceptual function when used as a substrate for reinforcement learning, and (2) whether the development of circuit structure is explicitly coded or develops with a degree of random-ness. Here, we address both principles in the expansion layer of insects, the mushroom body, using *Drosophila melanogaster* as our model.

### Principle 1: Is the adult circuit random?

Associative learning circuits like the mushroom body allow animals to attach meanings to stimuli whose valence wasn’t known in advance. While the odorant perception and olfactory associative learning functions of the mushroom body have been well established (De Belle & Heisenberg, 1994; Laurent & Naraghi, 1994; Masse et al., 2009; Murthy et al., 2008), debate about the random or structured nature of the connectivity continues (Caron et al., 2013; Eichler et al., 2017; Ellis et al., 2023; Ganguly et al., 2024; Gruntman & Turner, 2013; Hayashi et al., 2022; Jefferis et al., 2004; F. Li et al., 2020; Marin et al., 2002; Murthy et al., 2008; Tanaka et al., 2004; Wong et al., 2002; J.-Y. Yang et al., 2023; Zheng et al., 2022). Because learning can select experience-relevant representations from an arbitrary map, random developmental wiring can be a sufficient, or even a theoretically superior, substrate (Albus, 1971; Cayco-Gajic et al., 2017; Jortner et al., 2007; Litwin-Kumar et al., 2017; Minsky, 1952, 2016; Rosenblatt, 1958). Kenyon cells encode odor identity by acting as coincidence detectors: They fire only when multiple of their inputs are active (Gruntman & Turner, 2013). Thus, the number and identity of the scents that can be encoded is directly determined by the diversity of the combinations of inputs (Cayco-Gajic et al., 2017; Litwin-Kumar et al., 2017). If we think of KCs as picking PN inputs as if from a bag of marbles, random choice makes it unlikely that two KCs would have the same combination of inputs. Random sampling also has the benefit of spreading the combinations chosen by the 2000 KCs equally among the millions of combinations that are possible to draw. Both minimizing the chance of overlapping combinations and distributing combinations evenly over what is possible are suited for the mushroom body’s role in representing the diverse and evolutionarily unpredictable scents the animal may encounter.

Despite the theoretical benefit of complete randomness, consistent functional correlations have been observed in Kenyon cell odor responses, and spatial correlations have been observed in projection neuron processes (Gruntman & Turner, 2013; Jefferis et al., 2007; H.-H. Lin et al., 2007; Tanaka et al., 2004; J.-Y. Yang et al., 2023; Zheng et al., 2022). These biases and correlations are also hypothesized to improve fitness by prioritizing specific odor representations and relationships (Ellis et al., 2023; J.-Y. Yang et al., 2023; Zheng et al., 2022). Stochastic patterns and processes can seem inconsistent with the robust molecular mechanisms in biology and are out of step with deterministic mechanisms that have been discovered in the precise and repetitive circuits favored in developmental studies. These patterns and points of view were used to argue that the calyx is structured. Nevertheless, multiple lines of evidence supported the model that *connectivity* onto Kenyon cells is random: Analysis of the inputs to hundreds of KCs across different brains failed to find structure in input combinations, and connectomic analyses of inputs to larval olfactory Kenyon cells, adult olfactory Kenyon cells, and adult visual Kenyon cells each showed that structure in inputs to individual Kenyon cells was mostly lacking, and that input patterns found in one animal or hemisphere were not found again in other animals or hemispheres (Caron et al., 2013; Eichler et al., 2017; Ganguly et al., 2024; Zheng et al., 2022).

To gauge the extent of randomness in the actual connectivity between PNs and KCs, we unify analyses of an electron microscopy reconstruction of the full adult calyx (Scheffer et al., 2020) with numerous light level analyses performed in the last twenty years. Our analyses bring together what appeared to be competing findings: From the point of view of cell types, the calyx is in fact orderly, whereas the connectivity between those cell types is less predictable. In the calyx, each PN type makes different and predictable numbers of axonal boutons and places them in predictable locations; A-P location is largely determined by a cell’s birth order and neuroblast, M-L positioning by the number of collaterals made by that neuron, and D-V by the length of that collateral. Indeed, relative positions of projection neuron outputs in the calyx and the predictably wired lateral horn mirror one another (Jefferis et al., 2007; Tanaka et al., 2004). The dendrites of Kenyon cells are also organized in the calyx according to cell type. Principle neurites and dendrites of KCs born from different neuroblasts occupy four overlapping medial-lateral wedges, as described previously, and dendrites of different major types are differentially distributed along the anterior-posterior axis (T. Lee et al., 1999; H.-H. Lin et al., 2007; Zhu et al., 2003). Similarly, lineally related “sister” neurons structure columns in the mammalian cortex and have correlated activity, while sets of horizontally arrayed sisters have been proposed to function as processing units in the hippocampus (Y. Li et al., 2012; H.-T. Xu et al., 2014; Y.-C. Yu et al., 2009).

We find that the structure of cell type overlaps in the calyx, combined with the spatially constrained anatomy of individual Kenyon cells, produces correlations in which types of olfactory PNs are available to Kenyon cells born from different clones or of different types. This likely produces the observed anatomic and some of the functional correlations in odor tuning exhibited by some subsets of Kenyon cells (J.-Y. Yang et al., 2023; Zheng et al., 2022). In our developmentally-based model, we find that previously competing theories are both right—yes, cell types are structured in particular locations in the calyx, and this can give rise to regional correlations in Kenyon cell inputs or odor representations; and yes, Kenyon cells appear to choose randomly from the particular set of projection neurons spatially available to them.

This model of spatially constrained but stochastic input choice could also explain patterns of inputs to cerebellar granule cells which vary across individual cells, but exhibit regional correlations (Gilmer & Person, 2017; Huang et al., 2013). Our model could also describe the regionalization of sensory inputs to the cortex even if cortical neurogenesis programs are themselves homogenous (Di Bella et al., 2021). However, this mechanism may not describe all connections in all expansion layers: A connectomic analysis of neocortex from an adult mouse found that some axons synapsed with more than one dendritic spine from the same postsynaptic neuron. The frequency of these redundant connections could not be recapitulated in a model where axons randomly synapsed with nearby spines, suggesting these neurons instead have some method of choosing partners beyond spatial proximity (Kasthuri et al., 2015).

### Principle 2: How random is development?

PNs make the calyx a bag of predictably positioned marbles that are not equally mixed. Each Kenyon cell selects marbles/inputs without regard for their identity but only from a small region of the bag. In our ablation experiments above and previously, we removed half the colors of marbles from the bag and found that Kenyon cells continued to make the same number of selections. This suggests that Kenyon cells default to forming partnerships, rather than spurning unexpected input distributions. Indeed, KCs seem eager to form synapses with diverse PN inputs. Previously, we observed that when we increased the number of dendrites made by KCs, each cell could respond to more odors suggesting they connected with more PN types. This behavior is consistent with many examples in the literature of neurons synapsing promiscuously (Wolterhoff & Hiesinger, 2024).

Neurons in culture will synapse with whoever they touch, or with themselves (Bekkers & Stevens, 1991; Van der Loos & Glaser, 1972). The existence of “neuronal self-avoidance” systems (Dscams, clustered protocadherins) implies that neurons will form synapses unless told not to (Grueber & Sagasti, 2010; Williams et al., 2021). In fact, even in normal development, there is some degree of promiscuity in wildtype processes such as synaptic pruning where neurons initially form abundant synapses that later get refined to allow “correct” synaptic partnerships (Lieberman et al., 2019; Wilton et al., 2019). Additionally, by simply slowing down filopodial kinetics and allowing more time for synapse formation, the pool of synaptic partners for a neuron can be drastically altered (Kiral et al., 2020). This suggests that neurons can form synapses with even incorrect partners, if given the opportunity. Most striking of all, we were inspired by work from Matthew Pecot and others which showed that in the *Drosophila* visual system, removing Ig-SF molecules results in promiscuous synaptogenesis with partners that are not otherwise preferred (C. Xu et al., 2019), as we have also observed (Yoo et al., 2023). In this model, neuronal specificity factors are not required for synapse formation, but allow neurons to rank potential partners. While Kenyon cells still use Dscams for self-repulsion, we reasoned that their natural state might otherwise be akin to this underlying promiscuity (Wang et al., 2002). Just as “choosy neurons” can become more promiscuous when Ig- SFs are experimentally removed, Kenyon cells can be made choosy by Ig-SF addition.

At first blush, this appears to be a classic example of “Peters’s rule,” where guidance processes lead a neuron to a precise location and it synapses promiscuously with whoever is there (Peters & Feldman, 1976). However, the mechanism we propose is not free of molecular constraint. Rather, as olfactory projection neurons must make specific partnerships in the antennal lobe and lateral horn, they produce a diversity of cell surface molecules which then likely must be dealt with by the Kenyon cells. As randomized mechanisms vary surface molecule expression in other contexts (Aresta-Branco et al., 2019; Magklara & Lomvardas, 2013; Williams et al., 2021), we initially embarked on transcriptional analysis of Kenyon cells expecting to find a stochastic expression system that would allow Kenyon cells to actively but randomly choose presynaptic partners. Our data lead instead to a hypothesis we call “see-no-Sperry,” in which a natural and general depletion of specificity molecules allows Kenyon cells to ignore molecular variation among their potential inputs and draw blindly from the bag of marbles.

There is likely a continuum of partner choice stringency across the brain, with Kenyon cells at one end of the spectrum not only tolerating partner choice variation, but using it for their circuit function. While the Ig-SF depletion of Kenyon cells is correlated with Kenyon cell promiscuity, other mechanisms may also contribute. First, there may be sources of cell autonomous molecular variation among Kenyon cells that we have not discerned here. Second, it is possible that the Ig-SF molecules produced in projection neurons do not reach their axons in the calyx, and that PN surfaces are more similar than their transcriptional profiles would suggest. However, individual projection neurons make deterministic partnerships in the lateral horn, and plausible mechanisms that would differentiate these two output sites have not been discovered. Finally, biophysical characteristics of the growth and stabilization of Kenyon cell processes could allow for unique modes of input acquisition, even surmounting molecular constraints. In this model, the depletion of specificity molecules in Kenyon cells could serve other unique biophysical characteristics of their development.

Though individual Kenyon cells are able to connect promiscuously, the proximity to potential partners is still constrained by a few different mechanisms. Kenyon cell neurites seem to inherit their position from their neuroblast mother and timing of their birth. While there don’t seem to be physical barriers dividing the calyx, typical Kenyon cells do not reach their dendrites along the whole medial-lateral axis. We don’t yet know what kind of process constrains dendritic spread of Kenyon cells; they could have limited membrane or are unlikely to reach far without encountering a viable partner. As KCs are most restricted along the M-L axis, the M-L positioning of PN boutons determines the mix of inputs across calyx quadrants (H.-H. Lin et al., 2007). The M-L positioning of PN boutons seems to be linked to the specific number of collaterals made by that cell. We don’t yet know what determines each PN type’s propensity to make collaterals.

While olfactory Kenyon cells and visual Kenyon cells both accept promiscuous inputs, they each do so only from a precise group of partners (Ganguly et al., 2024; Marin et al., 2002; Wong et al., 2002). Overall, we hypothesize that while Kenyon cells ignore the diversity among their olfactory or visual projection neuron partners, they could have molecular determinants for their presynaptic partners as a group, such as an “olfactory universal glue” or “visual universal glue.” Alternatively, visual and olfactory subdivisions of the calyx could arise secondarily from spatial separation of these neuronal pools. Of course, many additional aspects of Kenyon cell anatomy and connectivity are developmentally constrained, from the quantity of PN inputs they receive, to the targeting of Kenyon cell axons to specific mushroom body lobes.

### Evolutionary perspective

We find it interesting that in the anatomic and developmental system we describe here, it is up to projection neurons to determine how well-mixed the calyx is. We do not know from what circuit substrate the system evolved, but because Kenyon cells seem to be outliers versus the other cell types we analyzed in their lack of specificity molecules, we guess that the ancestral cells from which Kenyon cells evolved were more molecularly constrained. Once the Kenyon cells acquired suppression of surface molecules, projection neuron developmental programs could cause the calyx to be functionally organized, such that information about related odors is stratified across it. The inferior colliculous could be an example of this type of organization, where signals are tonotopically mapped along one axis of the tissue but mixed within tonotopic domains (Drotos & Roberts, 2024). Or, along other evolutionary paths, projection neuron processes could become mixed such that the mushroom body functions as a mathematically ideal Marr-Albus expansion layer (Albus, 1971; M. Ito, 1970; Marr, 1969). Across extant insects, which have extremely diverse mushroom bodies, there may be a wide spectrum of functional correlation versus mixing in the inputs to the calyx (Farris, 2015). Kenyon cells in any species could use the same mechanism—spatially constrained promiscuity—and end up with very different functional responses. The fruit fly mushroom body provides just one example, in which inputs from different sensory modalities are strictly separated; the projection neurons exhibit diverse weights; and the projection neuron processes from different glomeruli are fairly well mixed in space in the calyx.

While inherent promiscuity in synapse formation seems antithetical to the specificity needed to wire functional neuronal circuits, we argue that the relative fitness of an adult structure is only meaningful with respect to how difficult it is to build. Here, the “type-agnostic” system through which we hypothesize Kenyon cells acquire their olfactory presynaptic partners can provide variation in odor representation across cells while having ancillary benefits: (1) It can flexibly incorporate evolution of projection neuron odor representations, providing both evolutionary robustness and evolvability. (2) It is compact in terms of the amount of information it requires to be built during development. (3) It is scalable across different sized Kenyon cell networks and indeed should require minimal increase in genomic information across variation in odorant receptor gene family, number of PN types, or number of Kenyon cells. Similar principles could underlie the development of vastly larger expansion layers in other organisms.

## Acknowledgements

We thank Bloomington Drosophila Stock Center, Liqun Luo, Yoshi Aso, and Gerry Rubin and Heather Dionne for sharing fly strains; and Kamlai Saiya-Cork (University of Michigan Flow Cytometry Core) and Gregg Sobocinski (MCDB Imaging Core) for technical assistance. Audrey Drotos and Margarita Brovkina provided conceptual insight. Vanessa Ruta and Claude Desplan provided support for the project, and Ruta lab members assisted in generation of bulk sequencing libraries. Robin Hiesinger and Clowney lab members provided input on the manuscript. EMT-K was supported by NIH T32 DC00011, MA by the University of Michigan Rackham Predoctoral Fellowship and Rackham Graduate Student Research Grant, and FRG by the MNI Magnificent Michigan Fellowship. B.S. was supported by a Long-Term Fellowship LT000010/2020-L from the Human Frontier Science Program. EJC is a McKnight Scholar, Pew Biomedical Scholar, and Rita Allen Milton Cassell Scholar. Funding was provided by NIDCD R01DC018032 (to EJC)

## Author Contributions

MA, EMT-K, and EJC conceived of the project, designed and managed experiments and analyses, procured funding, and prepared figures and wrote the manuscript with contributions of all authors. EMT-K contributed connectomic analyses and conducted projection neuron and Kenyon cell manipulation experiments with FRG. MA molecularly profiled developing Kenyon cells with the assistance of DLW and performed transcriptional analyses. BS generated scRNAseq datasets of adult Kenyon cell types labeled *a priori* and shared them prior to publication. YZK conceived of and led transcriptomic comparison of Kenyon cells with other pure cell types.

## Methods

### Key Resources Tables

#### Fly alleles

| Reagent | Source | Citation |
| --- | --- | --- |
| GH146 Gal4 (II) | BDSC 30026 | (Stocker et al., 1997) |
| VT033008-Gal4 (attp2) | Yoshi Aso, Janelia Farms Research Campus | (Kvon et al., 2014) |
| 89B01-Gal4 (attp2) | BDSC 40541 | (Jenett et al., 2012) |
| OK107Gal4 (IV) | BDSC 854 | (Connolly et al., 1996) |
| 10XUAS-IVS-myr::tdTomato (attp40) | BDSC 32222 | (Pfeiffer et al., 2012) |
| UAS-CD8GFP (III) | BDSC 5130 | (T. Lee et al., 1999) |
| 10XUAS-IVS-myr::GFP (attP2) | BDSC 32197 | (Pfeiffer et al., 2012) |
| UAS-Cbβ/DT-A.I[18] (II) | BDSC 25039 | (Han et al., 2000) |
| UAS luc RNAi (III) | BDSC 35788 | (Zirin et al., 2020) |
| UAS DPR4 OE (86Fb) | FlyORF F002762 | (Bischof et al., 2013) |
| VT033006:: LexAp65 (JK22C) | Heather Dionne and Gerry Rubin, Janelia Farms Research Campus | (Cachero et al., 2020; Tirian & Dickson, 2017) |
| 91G04-LexA, LexAopTom (attp40) | Liqun Luo, Stanford University | (Xie et al., 2019) |
| LexAopTomato (suHw attp5) | FBti0160868 |  |
| MB247-LexAVP16 (III) | Scott Waddell, University of Oxford | (Pitman et al., 2011) |
| LexAop CD2-GFP (III) | BDSC 66544 | (Lai & Lee, 2006) |
| GH146QF, QUAS-mtdTomato-3xHA (II) | BDSC 30037 | (Potter et al., 2010) |
| MZ19-QF (II) | BDSC 41573 | (Hong et al., 2012) |
| QUAS-mtdTomato-3xHA (II) | BDSC 30004 | (Potter et al., 2010) |
| P(caryP) attp40 | BDSC 36304 | (Markstein et al., 2008) |
| Empty landing site in 86FB | Fly ORF/ FBti0076525<br>(removed 3XP3-RFP<br>with CRE) | (Bischof et al., 2013) |
| OK107-QF2>QUAS-GFP, UAS-mtdTomato | BDSC 66472 |  |
| P{y[+t7.7] w[+mC]=R13F02-<br>p65.AD}attP40/CyO; P{y[+t7.7]<br>w[+mC]=R89B01-GAL4.DBD}attP2 | BDSC 68265 | (Aso et al., 2014) |
| P{y[+t7.7] w[+mC]=R26E07-<br>p65.AD}attP40/CyO; P{y[+t7.7]<br>w[+mC]=R30F02-GAL4.DBD}attP2 | BDSC 68322 | (Aso et al., 2014) |
| P{y[+t7.7] w[+mC]=R35B12-<br>p65.AD}attP40; P{y[+t7.7]<br>w[+mC]=R34A03-<br>GAL4.DBD}attP2/TM6B, Tb[1] | BDSC 68370 | (Aso et al., 2014) |
| P {y[+t7.7] w[+mC]=R13F02-<br>p65.AD}attP40; P{y[+t7.7]<br>w[+mC]=R85D07-GAL4.DBD}attP2 | BDSC 68383 | (Aso et al., 2014) |
| P{y[+t7.7] w[+mC]=R13F02-<br>p65.AD}attP40; P{y[+t7.7]<br>w[+mC]=R58F02-GAL4.DBD}attP2 | BDSC 68255 | (Aso et al., 2014) |
| P{y[+t7.7] w[+mC]=R52H09-<br>p65.AD}attP40; P{y[+t7.7]<br>w[+mC]=R18F09-GAL4.DBD}attP2 | BDSC 68267 | (Aso et al., 2014) |
| GAL4<sup>dpr1-MI02201-TG4.1</sup> | FBal0349605 | Zinn and Zipursky labs,<br>via Pelin Volkan |
| GAL4<sup>dpr3-MI05963-TG4.1</sup> | FBal0349598 | Zinn and Zipursky labs,<br>via Pelin Volkan |
| GAL4<sup>dpr5-MI11085-TG4.1</sup> | FBal0349594 | Zinn and Zipursky labs,<br>via Pelin Volkan |
| GAL4<sup>dpr6-MI01358-TG4.1</sup> | FBal0349602 | Zinn and Zipursky labs,<br>via Pelin Volkan |
| GAL4<sup>dpr8-MI06778-TG4.0</sup> | FBal0349599 | Zinn and Zipursky labs,<br>via Pelin Volkan |
| GAL4<sup>dpr9-MI03594-TG4.1</sup> | FBal0349603 | Zinn and Zipursky labs,<br>via Pelin Volkan |
| GAL4<sup>dpr10-MI03557-TG4.1</sup> | FBal0349595 | Zinn and Zipursky labs,<br>via Pelin Volkan |
| GAL4<sup>dpr11-MI01743-TG4.1</sup> | FBal0349596 | Zinn and Zipursky labs,<br>via Pelin Volkan |
| GAL4<sup>dpr12-MI01695-TG4.1</sup> | FBal0349600 | Zinn and Zipursky labs,<br>via Pelin Volkan |
| GAL4<sup>dpr15-MI01408-TG4.1</sup> | FBal0328010 | Zinn and Zipursky labs,<br>via Pelin Volkan |
| GAL4<sup>DIP-α-MI02031-TG4.1</sup> | FBal0345530 | Zinn and Zipursky labs,<br>via Pelin Volkan |
| GAL4<sup>DIP-δ-MI08287-TG4.1</sup> | FBal0345531 | Zinn and Zipursky labs,<br>via Pelin Volkan |
| GAL4<sup>DIP-η-MI07948-TG4.1</sup> | FBal0345533 | Zinn and Zipursky labs,<br>via Pelin Volkan |
| DIP-γ<sup>MI03222-GAL4</sup> | FBal0319064 | Zinn and Zipursky labs,<br>via Pelin Volkan |
| DIP-κ<sup>MI01295-Gal4</sup> | No record, generated<br>from FBal0264429 by<br>Carillo et al | Zinn and Zipursky labs,<br>via Pelin Volkan |
| GAL4<sup>DIP-θ-MI03191-TG4.1</sup> | FBal0345534 | Zinn and Zipursky labs,<br>via Pelin Volkan |
| GAL4<sup>DIP-ζ-MI03838-TG4.0</sup> | FBal0345532 | Zinn and Zipursky labs,<br>via Pelin Volkan |

#### Antibodies

| Reagent | Source | Identifier |
| --- | --- | --- |
| anti-ChAT (mouse monoclonal) | Developmental Studies Hybridoma Bank | Cat# chat4b1; RRID: AB_528122 |
| anti-DsRed (rabbit polyclonal) | Clontech/Takara Bio | Cat# 632496; RRID: AB_10013483 |
| anti-GFP (chicken polyclonal) | Cai Lab | N/A |
| anti-brp (mouse monoclonal) | Developmental Studies Hybridoma Bank | Cat# nc82; RRID: AB_2314866 |
| Anti-GFP (rabbit polyclonal) | Fisher | Cat# A11122 |
| Anti-RFP (rat monoclonal) | Chromotek | Cat# 5f8 |
| Alexa 488 anti-chicken (goat polyclonal) | Fisher | Cat# A32931; RRID: AB_2762843 |
| Alexa 488 anti-rabbit (goat polyclonal) | Fisher | Cat# A11008 |
| Alexa 568 anti-rabbit (goat polyclonal) | Fisher | Cat# A-11036; RRID: AB_10563566 |
| Alexa 568 anti-rat (goat polyclonal) | Fisher | Cat# A11077 |
| Alexa 647 anti-mouse (goat polyclonal) | Fisher | Cat# A-21236; RRID: AB_2535805 |

#### Chemicals, peptides, and recombinant proteins

| Reagent | Source | identifier |
| --- | --- | --- |
| Paraformaldehyde | EMS | 15710 |
| DAPI (used in stainings though not shown in figures) | Sigma | D9542 |
| 10X PBS Gibco | Fisher | 70-011-044 |
| Normal Goat Serum | MP biomedical | 8642921 |
| Triton | Sigma | X100 |
| Sodium chloride NaCl | Sigma | S7653 |
| Potassium chloride KCl | Fisher | P217-500 |
| Calcium chloride<br>CaCl<sub>2</sub>*2H<sub>2</sub>O | Fisher | C70-500 |
| Magnesium chloride<br>MgCl<sub>2</sub>*6H<sub>2</sub>O | Sigma | M2670 |
| HEPES sodium salt | Sigma | H7006 |
| Trehalose | Sigma | T9531 |
| Sucrose | Sigma | 84097 |
| Sodium bicarbonate NaHCO<sub>3</sub> | Fisher | S233-500 |
| Sodium phosphate<br>NaH<sub>2</sub>PO<sub>4</sub>*H<sub>2</sub>O | Fisher | S369-500 |
| Propyl gallate | Sigma | 02370 |
| Schneider's *Drosophila* Medium | Gibco | 21720024 |
| Collagenase from *Clostridium histolyticum* | Sigma | C0130 |
| Dispase II | Sigma | D4693 |
| Actinomycin D | Sigma | A1410 |
| 1X Dulbecco's PBS | Corning | 21-031-CV |
| 50 mg/mL UltraPure BSA | Invitrogen | AM2618 |
| 5 mM DRAQ5 Fluorescent Probe Solution | Thermo Scientific | 62254 |

#### Software and algorithms

| Reagent | Source | Identifier |
| --- | --- | --- |
| FIJI | https://imagej.net/software/fiji/downloads | (Schindelin et al., 2012) |
| R | https://www.R-project.org/ | (R Foundation for Statistical Computing, 2023) |
| Neuprint: An open access tool for EM connectomics | https://neuprint.janelia.org/ | (Clements et al., 2020) |
| R package natverse | https://natverse.org/ | (Alexander Shakeel Bates et al., 2020) |
| R package Seurat | https://satijalab.org/seurat/ | (Hao et al., 2021; Satija et al., 2015; Stuart et al., 2019) |
| R package Clustree | https://lazappi.github.io/clustree/ | (Zappia & Oshlack, 2018) |
| R package UCell | https://github.com/carmonalab/UCell | (Andreatta & Carmona, 2021) |
| R package RColorBrewer | https://CRAN.R-project.org/package=RColorBrewer | (Neuwirth, 2022) |
| python | http://www.python.org | Python Software Foundation |
| python package scanpy | https://github.com/scverse/scanpy | (Wolf et al., 2018) |
| python package scvi-tools | https://github.com/scverse/scvi-tools | (Gayoso et al., 2022) |

#### Other

| Reagent | Source | Identifier |
| --- | --- | --- |
| Fly food | Lab Express, Ann Arbor MI | “B” Food |
| Cell strainer caps (used as colanders for stainings) | Fisher | BD 352235 |
| Binder reinforcement stickers (for mounting brains) | Office Depot | Avery 5722 |
| pluriStrainer Mini 20 µm | pluriSelect | 43-10020-40 |

#### Fly genotype by figure

| Figure | Genotype |
| --- | --- |
| 4A | UAS-CD8GFP/CyO; GH146QF, QUAS-Tomato/TM6B; OK107-Gal4/OK107-Gal4 |
| 4B | GH146 Gal4, UAS-mtdTomato/CyO; MB247-LexA, LexAop CD2-GFP/TM2 |
| 4C-J | Developmental time points: UAS-CD8GFP/CyO; GH146QF, QUAS-Tomato/TM6B; OK107-Gal4/OK107-Gal4 and GH146 Gal4, UAS-mtdTomato/CyO; MB247-LexA, LexAop CD2-GFP/TM2 |
| 4C-J | Adult time point:<br>P{y[+t7.7] w[+mC]=R13F02-p65.AD}attP40/CyO; P{y[+t7.7] w[+mC]=R89B01-GAL4.DBD}attP2; OK107-QF2>QUAS-GFP, UAS-mtdTomato<br>P{y[+t7.7] w[+mC]=R26E07-p65.AD}attP40/CyO; P{y[+t7.7] w[+mC]=R30F02-GAL4.DBD}attP2; OK107-QF2>QUAS-GFP, UAS-mtdTomato<br>P{y[+t7.7] w[+mC]=R35B12-p65.AD}attP40; P{y[+t7.7] w[+mC]=R34A03-GAL4.DBD}attP2/TM6B, Tb[1]; OK107-QF2>QUAS-GFP, UAS-mtdTomato<br>P{y[+t7.7] w[+mC]=R13F02-p65.AD}attP40; P{y[+t7.7] w[+mC]=R85D07-GAL4.DBD}attP2; OK107-QF2>QUAS-GFP, UAS-mtdTomato<br>P{y[+t7.7] w[+mC]=R13F02-p65.AD}attP40; P{y[+t7.7] w[+mC]=R58F02-GAL4.DBD}attP2; OK107-QF2>QUAS-GFP, UAS-mtdTomato<br>P{y[+t7.7] w[+mC]=R52H09-p65.AD}attP40; P{y[+t7.7] w[+mC]=R18F09-GAL4.DBD}attP2; OK107-QF2>QUAS-GFP, UAS-mtdTomato |
| Supplementary 4C-E | DIP and dpr Gal4 insertions as indicated (described in table of fly alleles) were crossed with UAS-GFP alleles |
| 6A-E | Control: VT033006-LexA, LexAop tom/caryPattp40; VT033008-Gal4, UAS-GFP/+<br>Experimental: VT033006-LexA, LexAop tom/UAS-DTA; VT033008-Gal4, UAS-GFP/+ |
| 6F-I | Control: MZ19QF, QUAS tomato/+; VT033008-Gal4, UAS GFP/ UAS luc RNAi<br>OR<br>MZ19QF, QUAS tomato/ caryP attp40; VT033008-Gal4, UAS GFP/ +<br>Experimental: MZ19QF, QUAS tomato/UAS DTA; VT033008-Gal4, UAS GFP/+ |
| 6 J-L | Control:<br>MZ19QF, QUAS tomato/ caryP attp40; VT033008-Gal4, UAS GFP/ +<br>91G04-LexA, LexAopTom/ caryP attP40; VT033008-Gal4, UAS GFP/+<br>Experimental:<br>MZ19QF, QUAS tomato/UAS DTA; VT033008-Gal4, UAS GFP/+<br>91G04-LexA, LexAopTom/ UAS-DTA; VT033008-Gal4, UAS GFP/+ |
| 7 | Control: MZ19QF, QUAS tom/CyO or sp; 89B01 gal4, UAS GFP/Empty landing site in 86FB<br>Experimental: MZ19QF, QUAS tom/CyO or sp; 89B01 gal4, UAS GFP/UAS DPR4 OE |

## EM analysis

### Downloading synapse locations from neuprint+

Cartesian coordinate locations of synapses between uniglomerular projection neurons and Kenyon cells from the Hemibrain: v1.2.1 were downloaded from neuprint+ using a custom query which retrieved location of synapses between 115 uniglomerular projection neurons and any cell with ‘KC’ in its instance (name). The unique Hemibrain cell body Id for both pre and postsynaptic partners was also included in the output. The custom query was run 6x with batches of about ∼20 PN cell body Ids at a time.

Example query:

MATCH(pn:Neuron)-[:Contains]->(:‘SynapseSet’) -[:Contains]->(pns:Synapse)- [:SynapsesTo]->(kcs:Synapse)<-[:Contains]-(:‘SynapseSet’)<-[:Contains]-(kc:Neuron) WHERE pn.bodyId IN [1639234609, 818983130, 1796818119, 1734350788, 1788300760, 5813054697, 5813024710, 5813024601, 754534424, 1670934213, 1734350908, 5813039315, 722817260, 1888572074, 5813055184, 1609542060, 5813039465, 850375847, 1914140664, 5813024712] AND kc.instance=∼’KC.*’ AND kc.status = "Traced" AND pns.type = "pre" AND pns.‘CA(R)’ RETURN pn.bodyId, pn.instance, kc.bodyId, kc.instance, pns.location.x, pns.location.y, pns.location.z, pns.type

The list of 115 cell body Ids for uniglomerular projection neurons was obtained by using the “find neurons” function in neuprint+ to search for neurons with inputs in AL(R) and outputs in CA(R). The output of 136 neurons with ‘PN’ in their instance was further filtered to exclude 21 multiglomerular PNs using a list of Hemibrain counterparts of PNs reconstructed from the FAFB by Bates et al. In the Hemibrain, only 112 of these PNs made synapses with KCs in the calyx. For all analyses the 112 PNs were filtered to 105 olfactory PNs by removing 7 thermo- and hygro-sensory PNs innervating the ventral posterior glomeruli- VP1d, VP1m, VP2, VP3, VP4, and VP5.

25,718 unique presynapses from uniglomerular olfactory projection neurons to KCs were found. Monosynaptic connections between PNs and KCs were disproportionally present in the data and so filtered out as false positives down to a total of 25,394 presynapses. For all analyses only synapses between olfactory PNs and main calyx KCs were considered. These amounted to a total of 24,791 synapses.

### Downloading, and trimming neuron skeletons

All PN and KC skeletons were downloaded using the “neuprint_read_neurons” function in the neuprintR R package from our list of body Ids from PNs and KCs. Skeletons were roughly trimmed to a cube surrounding the calyx which was 5% larger on each end of the x and z axis and 10% larger on the y axis than the bounds of the CA(R) mesh (neuprint_ROI_mesh(‘CA(R)’)).

### Cell types and developmental feature annotation

Each PN was annotated with an AL glomerular target/ cell type and neuroblast origin in neuprint+ and in Bates 2020 (Alexander S. Bates et al., 2020; Scheffer et al., 2020). We further added developmental stage of birth and birth order information from Yu et al 2010 and Lin et al 2012 (S. Lin et al., 2012; H.-H. Yu et al., 2010).

We used KC type annotations from neuprint+ (Scheffer et al., 2020). γ KCs were KCs with instances including ‘KCg-m’,’KCg-d’, ‘KCg-t’, ‘KCg-s’, and ‘KCy(half). α’β’ KCs were KCs with instances including ’KCa’b’-m’, ’KCa’b’-ap1’, and ’KCa’b’-ap2’. αβ KCs were those with instances including ’KCab-s’,’KCab-m’,’KCab-c’,’KCab-p’. Unknown KCs included instances of ‘unknown’, ‘KC part due to gap’, ‘KC (incomplete?)’. Kenyon cells were filtered to only the 1728/ 1874 that innervate the olfactory calyx (not γ-d, α’β’ ap1, nor αβp).

We assigned each KC to one of 4 neuroblasts of origin by using the location of its axon in the pedunculus where axons are known to segregate in a clonal fashion (T. Lee et al., 1999). We started by assigning putative neuroblast origin to the pupally born αβ KCs. As the latest born KC subtype, their axons are surrounded by α’β’ and γ KCs born from their same neuroblast and so are most separated from αβ cells born from different neuroblasts. We used k-means clustering to sort points of intersection between each KC and each of three planes perpendicular to the pedunculus into 4 groups. 806/809 αβ KCs were assigned the same cluster at each of the three planes and so were sorted into neuroblasts named A,B,C,D from medial to lateral.

We next assigned α’β’ KCs to neuroblasts by again calculating the points of intersection between axons and planes perpendicular to the pedunculus at 3 different levels. Rather than using k-means clustering of points of intersection; each unassigned KC was assigned to the clone to which it had the lowest mean distance from each of the axons already assigned. 246/336 α’β’ KCs were automatically sorted into neuroblasts in this manner. Finally, the earliest born γ KCs were sorted using the same method proximity based method to sorted αβ and α’β’ KCs. 264/593 γ KCs were automatically assigned using this method. KCs of all three types which were not able to be computationally assigned to clones using axon intersections in the pedunculus were manually assigned to clones based on their soma position which is also known to be sorted by clones (T. Lee et al., 1999) and further proofread with pedunculus position. The number of cells of each type assigned to each clone is roughly equivalent, as expected (Figure 2A).

### Annotating subcellular features

Using the locations of synaptic contacts, we obtained spatial information about sub-cellular features of both projection neurons and Kenyon cells in the calyx. For projection neurons, we estimated coordinate centers of boutons and of collateral branch points from the main axon. For Kenyon cells, we estimated coordinate centers of claws.

Bouton centers were calculated as the coordinate center of the presynapses that made up the bouton. Presynapses clustered together most often at the ends of collaterals into putative boutons. We used k-means clustering to quickly assign groups of presynapses to individual boutons using the number of putative boutons as the number of clusters. We next manually ‘proof-read’ bouton assignments reassigning clear errors, e.g. presynapses clearly on other collateral branches or not immediately next to other presynapses assigned to that bouton. In sum we found 538 boutons which matches with our counts in Figure 5D (median = 524.5 boutons). We limited analyses to 436 olfactory PN boutons that were made up more than 3 presynapses.

Boutons most often are at the ends of collaterals which branch perpendicularly from the PN axon in the medial Antennal Lobe Tract (mALT). We extracted the locations of these collateral branch points by filtering branch points in the PN skeleton to those close to the mALT and then manually assigned them by plotting putative branch points onto the skeleton with bouton centers.

Collateral branch points were branch points at the base of collaterals that bore boutons. In sum we annotated 265 collateral branch points.

Each Kenyon cell claw center was calculated as the coordinate center of the presynapses of a single PN bouton opposed by a single Kenyon cell. In sum we annotated 10,970 Kenyon cell claws. We limited our analyses to 10,674 claws belonging to main calyx Kenyon cells. Kenyon cell claws that opposed fewer than 3 presynapses were also filtered out reducing their number to 10,195 in the olfactory calyx.

### Rotating coordinate space to match up with relevant calyx axes

To simplify analyses of the calyx we rotated all coordinates such that the longest extent of the calyx along which the mALT runs (M-L) was parallel to the X axis and the shortest axis (A-P) perpendicular to the mALT was parallel to the Y axis. Coordinates were rotated 285 degrees about the X axis and 200 degrees about the Y axis.

### Generation of random models of KC:PN connectivity

In order to ask if patterns of non-random connectivity between PNs and KCs could be explained by developmentally defined spatial limits on the neurites of both PNs and KCs, we generated a series of null models of connectivity to compare with observed connectivity in the Hemibrain. We considered 20 different developmentally defined groups of Kenyon cells: all olfactory Kenyon cells, Kenyon cells belonging to each of the 4 clones, Kenyon cells of each of the 3 types, and Kenyon cells belonging to each of the 12 groups of clone and type. For each group we generated 10,000 null models of connectivity in which each claw of each Kenyon cell is randomly assigned to a PN type. The probability of selecting any PN type was proportional to the number of boutons accessible to KCs of that group in the Hemibrain dataset.

### Conditional Input analysis

We performed conditional input analysis as described in Zheng 2022 (Zheng et al., 2022) on connectivity matrixes for each of our 20 developmentally defined KC groups. The output of this analysis is a matrix of PN types (51x51), which indicates the number of KCs connected to a column PN type given they also are connected to a row PN type. By comparing values computed using observed Hemibrain connectivity to distributions of values computed using the 10,000 random models we can assess how similar observed connectivity is to that of each of the random models.

## Fly husbandry

Flies were maintained on Bloomington food with a yeast sprinkle (“B” recipe, Lab Express, Ann Arbor, MI) at 25C on a 12:12 light-dark cycle with at least 60% humidity (provided by a beaker of water in the incubator). Flies for bulk RNAseq were maintained on Rockefeller University molasses food. For staging developmental time points, pupae were marked at the 0h APF “white pupae” stage. Animals for scRNAseq were collected +/-2 hours from the stated developmental time points (e.g. 48h APF were collected from 46-50h APF). Pupae for bulk RNAseq were collected at 44-46h APF. Flies for adult scRNAseq were maintained on Desplan lab standard cornmeal food at 25C on a 12:12 light-dark cycle without controlling for humidity. Females for adult scRNAseq were collected <2 weeks after eclosion.

## Flow cytometry

Flow cytometry was performed largely as described in (Brovkina et al., 2021). Brains were dissected for up to 60 minutes in Schneider’s medium supplemented with 1% BSA and placed on ice. Optic lobes were removed during dissection. After all dissections were completed, collagenase was added to a final concentration of 2mg/mL and samples were incubated at 37C for 20 minutes (adults) or 12 minutes (pupae), without agitation. Samples were dissociated by trituration and spun down at 300g, 4C, for 5 minutes, in a swing-out rotor. Collagenase solution was removed and replaced with PBS+0.1% BSA, and cells were passed through a cell strainer cap and supplemented with 50ng/mL DAPI before being subjected to flow cytometry on a FACS Aria II (University of Michigan Flow Cytometry Core for scRNAseq, Rockefeller University Flow Cytometry Core for bulk RNAseq). Plasticware for cell dissociation and collection was pre-treated by rinsing with PBS+1% BSA to prevent cells from sticking to bare plastic.

During flow cytometry, dead and dying cells were excluded using DAPI signal, and forward scatter and side scatter measurements were used to gate single cells. Using our dissociation methods, 50–90% of singlets appeared viable (DAPI-low). Fluorophore-positive cell rate varied with the prevalence of the cell population. During sorting, we made two adjustments to protect the fly primary cells, which were very delicate—we disabled agitation of the sample tube, and sorted using the “large nozzle,” e.g. 100μm, i.e. using larger droplet size and lower pressure. For bulk RNAseq, we sorted cells directly into Trizol-LS. For scRNAseq, we sorted cells into PBS+1% BSA.

Samples for adult scRNA-seq were prepared largely as described in (Özel et al., 2021). Brains were dissected for up to 60 minutes in Schneider’s medium and placed on ice. Optic lobes were removed during dissection. After all dissections were completed, collagenase and dispase were added to a final concentration of 2mg/mL each along with 45 uM Actinomycin D, and samples were incubated at 26C for 1 h without agitation. Samples were washed 2-3 times with DPBS+0.04% BSA, dissociated by trituration and passed through a 20 um PluriStrainer. Cells were stained with 5 uM DR before being subjected to flow cytometry on a FACS Aria II (New York University Genomics Core). Forward scatter area and width were used to gate single cells, and cells were sorted into DPBS+0.04% BSA.

## RNA isolation, library preparation, and sequencing for low-input bulk RNAseq on FAC-sorted neurons

To collect olfactory projection neurons and Kenyon cells for bulk RNAseq, we dissected, removed optic lobes, dissociated, and FAC-sorted about 100 brains (adult) or 25 brains (45h APF) for each replicate in which Kenyon cells were labeled by MB247LexA>LexAopGFP, and olfactory projection neurons by GH146-Gal4>UAS-mTdTomato. GFP-positive cells were 5-8% of viable singlets, and Tomato-positive were 0.4-1% of viable singlets. These numbers are consistent with the rate at which these cell types appear in the central brain (MB247+ Kenyon cells: ∼4000 out of ∼50,000; GH146+ PNs: 200-300 cells out of 50,000).

RNA isolation was performed as we described in detail previously (Brovkina et al., 2021). Briefly, we FAC-sorted cells directly into Trizol-LS and stored at -80C prior to RNA preparation.

Cells obtained in each replicate are shown in the table below. We followed the standard Trizol-LS protocol until the aqueous phase was isolated. We then passed the aqueous phase over Arcturus Picopure columns, including a DNAse treatment on the column, using modifications of the standard Picopure protocol described in (Brovkina et al., 2021). We quantified total RNA using a BioAnalyzer and aimed for 1ng starting material per library. As insect processing of rRNA makes the “RNA Integrity Number” irrelevant, we visually inspected total RNA profile to assess RNA quality.

For each replicate, we matched the amount of total RNA used for library prep across the three cell types, i.e. we matched Kenyon cell and “double negative” RNA amounts to the amount we obtained for olfactory PNs. Total rRNA was poly-A selected and libraries were prepared using SmartSeq V2 at the Rockefeller University Genomics Core and sequenced to a depth of 25 million reads on Illumina platforms as described in the table.

| Sample | Cells | RNA obtained | RNA used for library | Platform | Reads | Mapped reads | Uniquely mapped reads |
| --- | --- | --- | --- | --- | --- | --- | --- |
| Double Negative, adult, Rep 1 | 513,000 used | 9ng | 1.9ng | HiSeq 50 SR | 24,386,406 | 21,563,377 | 20,506,772 |
| Kenyon cells, adult, Rep 1 | 100,174 | 6.8ng | 1.9ng | HiSeq 50 SR | 19,339,250 | 16,809,729 | 12,977,111 |
| Olfactory PNs, adult, Rep 1 | 7,770 | 1.9ng | 1.9ng | HiSeq 50 SR | 25,139,441 | 22,665,097 | 20,327,411 |
| Double Negative, adult, Rep 2 | 250,000 used | 14ng | 1.8ng | HiSeq 50 SR | 24,687,509 | 20,938,375 | 20,938,375 |
| Kenyon cells, adult, Rep 2 | 98,458 | 6.6ng | 1.8ng | HiSeq 50 SR | 25,618,833 | 22,375,536 | 18,728,324 |
| Olfactory PNs, adult, Rep 2 | 5,139 | 1.8ng | 1.8ng | HiSeq 50 SR |  | 21,803,823 |  |
| Double negative, 45h APF, rep 1 | 150,000 used | n/a, pooled with rep 2 |  |  |  |  |  |
| Kenyon cells, 45h APF, rep 1 | 28,022 | n/a, pooled with rep 2 |  |  |  |  |  |
| Olfactory PNs, 45h APF, rep 1 | 3,735 | n/a, pooled with rep 2 |  |  |  |  |  |
| Double negative, 45h APF, rep 2 | 115,607 | Pooled with 1: 51ng | 1ng | NextSeq High Output 75SR | 42,126,450 | 32,437,525 | 30,623,495 |
| Kenyon cells, 45h APF, rep 1 | 26,076 | Pooled with 1: 5ng | 1ng | NextSeq High Output 75SR | 36,222,854 | 28,086,686 | 25,250,765 |
| Olfactory PNs, 45h APF, rep 2 | 3,127 | Pooled with 1: 1ng | 1ng | NextSeq High Output 75SR | 40,062,033 | 30,676,723 | 28,591,020 |

## Analysis of bulk RNAseq datasets

Analyses were performed in Galaxy. FASTQ files were groomed using FASTQ Groomer and aligned to the dm6 assembly of the *D. melanogaster* genome using TopHat. 76-90% of reads for each library mapped to the fly genome, with 3-22% of mapped reads aligning multiple times. We used Cufflinks to quantify the amount of transcription from each gene and assembled a table of FPKMs. We matched gene names to the Refseq “name2” annotations and provide additional gene-level information in the table. Enrichments of canonical markers of KCs (*ey*) and PNs (*Oaz*) are shown below, as well as the pan-neuronal marker nsyb, and *pros*, which is enriched in mature neurons. Adult data is shown as average of two replicates; 45h APF data is from a single library for each cell type. Numbers here are FPKM generated by Cufflinks/Cuttdiff. Nsyb is a pan-neuronal marker.

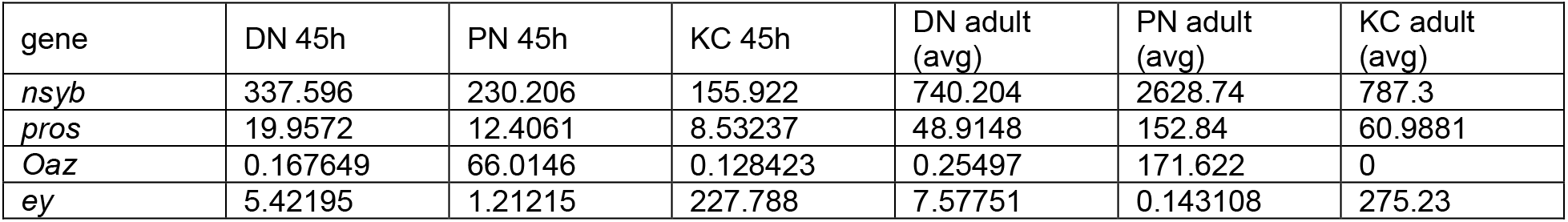

## Single-cell RNA-sequencing

### Flow cytometry for single-cell RNA-sequencing of developing Kenyon cells

To compensate for the effects of different strain backgrounds and driver lines, we labeled Kenyon cells with MB247-LexA (a transgene containing the *Mef2* promoter) or with OK107-Gal4 (a Gal4 insertion into the *eyeless* locus). We then FAC-sorted and performed scRNAseq at 36h, 48h, and 60h APF (one OK107 and one MB247 sample at each time point). Preparation for FACS was as described above in the flow cytometry methods section. Each time point was dissected on a separate day. We collected 40-80 brains per dataset.

To make sure all Kenyon cells were collected, regardless of fluorescence intensity, we used a loose gating protocol in FACS prior to scRNAseq. This was especially important for the MB247 driver, which labels different Kenyon cells with different intensity. The sorted populations therefore also contained small proportions of “non-Kenyon cells” that were sorted as false positives; we made use of these cells to compare expression of various groups of genes to that of Kenyon cells. Prior to library prep for sequencing, cells were centrifuged at 4°C to increase the cell concentration per microliter.

### Labeling of adult Kenyon cells and preparation for scRNAseq

To generate an adult Kenyon cell data set and annotate the seven molecularly defined cell subtypes (Aso et al., 2014), we single-cell sequenced seven genotypes that simultaneously expressed GFP in all Kenyon cells and mtdTomato in a defined Kenyon cell subtype(s). To generate such flies, the OK107-QF2>QUAS-GFP, UAS-mtdTomato line (BDSC# 66472) was crossed with each of the following subtype-specific split-GAL4 lines (Aso et al., 2014; Shih et al., 2019): γm and γd (BDSC #68265), γd (BDSC #68256), α′/β′m (BDSC #68322), α′/β′ap (BDSC #68370), α/βp (BDSC #68383), α/βc (BDSC #68255), α/βs (BDSC #68267).

The brains of adult female F1 progeny with the respective OK107-QF2>QUAS-GFP, GAL4-AD∩GAL4-DBD>UAS-mtdTomato genotype were dissected, visually checked for mtdTomato fluorescence in the mushroom body, dissociated into single cells, and stained with the DNA dye DR. DR +,GFP+ cells were FAC-sorted to enrich the samples for Kenyon cells. mtdTomato expression was not used for sorting. Given the prior knowledge that different classes of Kenyon cells separate well during the analysis of single-cell data (Croset et al., 2018), the brains of flies where the subtypes of different classes were labeled (e.g. γd, α′/β′ap, and α/βs) were pooled in different combinations before the dissociation to minimize batch effects. Each brain pool corresponds to one library and includes at least 8 brains. In total, 8 single-cell libraries were prepared using the 10X Chromium Next GEM Single Cell 3’ Kit v3.1. The libraries were sequenced on either Illumina NextSeq500 or NovaSeq6000 to the depth of at least 28k reads/cell. All the analyses were performed using scanpy and scvi-tools. After the initial round of clustering, clusters containing Kenyon cells were selected based on the expression of *dac*, *ey*, and *Pka-C1*. Then, Kenyon cells were re-clustered and class identities were assigned based on the expression of the γ marker ab, the α′/β′ marker *CG8641*, and the α/β marker *Ca-alpha1T*. Next, subtype identities were assigned using the following criteria: 1) expression of mtdTomato transcripts (for example, if library 1 contained brains expressing mtdTomato under a γd-specific split-GAL4 driver, the γ cluster expressing mtdTomato in this library was designated as γd), 2) gene expression correlation between pseudo-bulk data generated from each cluster and bulk data from sorted cells expressing fluorescent marker driven by subtype-specific split-GAL4s (Shih et al., 2019), and 3) relative size of each cluster as compared to the known relative size of each subtype population (Aso et al., 2014).

### scRNAseq and quality control metrics for pupal Kenyon cells

ScRNAseq data was generated using gel bead microfluidics (10x Chromium 3’ system) on a NovaSeq 6000 instrument, from FAC-sorted cells. Cell viability and aggregation was checked and number of cells to load was selected to ensure a low multiplet rate (∼3%) and ample expected cell recovery (at least 3000 cells). Raw fastq read files were processed using Cell Ranger (v6.1.0 or v7.1.0) with default parameters. A custom reference was built from the *Drosophila* FlyBase reference genome and transgene sequences (Gal4, LexA, QF, GFP, mtdTomato). For each sample, the sequencing was effective in recovering between ∼3,000-14,000 cells, with ∼60-80% sequencing saturation, ∼40,000- 100,000 mean reads per cell and ∼1600-2800 median genes per cell. UMI and cell barcode filtering was done to remove duplicate reads and non-cell-associated barcodes. A summary of metrics of acquired datasets is given below. In the sample name, MB247 or OK107 refers to the driver used for sorting for Kenyon cells. 36h, 48h, and 60h indicate the pupal stage.

| Sample | Estimated # of Cells | Mean Reads per Cell | Median Genes per Cell | # of Reads | Valid Barcodes | Sequencing Saturation | Reads Mapped to Genome | Fraction Reads in Cells | Total Genes Detected | Median UMI Counts per Cell |
| --- | --- | --- | --- | --- | --- | --- | --- | --- | --- | --- |
| MB247_36h | 11,655 | 40,896 | 2,374 | 476,637,400 | 97.70 % | 60.40% | 80.60% | 90.70% | 15,245 | 8,505 |
| OK107_36h | 13,312 | 42,628 | 2,251 | 567,466,841 | 97.10 % | 58.60% | 95.60% | 87.90% | 16,734 | 8,178 |
| MB247_48h | 3,614 | 88,528 | 1,648 | 319,939,146 | 97.60 % | 74.60% | 79.90% | 72.60% | 10,505 | 5,040 |
| OK107_48h | 4,911 | 99,944 | 2,748 | 490,826,737 | 98.00 % | 78.30% | 95.70% | 94.40% | 15,896 | 11,872 |
| MB247_60h | 3,405 | 64,026 | 2,321 | 218,007,780 | 98.70 % | 72.40% | 89.10% | 95.40% | 13,734 | 9,746 |
| OK107_60h | 13,708 | 53,719 | 1,903 | 736,379,055 | 97.90 % | 68.60% | 94.80% | 88.00% | 16,301 | 6,967 |

In the resulting gene-barcode matrix, each element gives UMI count associated with a gene (row) and a barcode (column). All steps of single-cell RNA-seq analysis were performed using Seurat (v4.3.0 and above). Cells were included if they had at least 1,000 but no more than 5,000 genes detected, and less than 5% mitochondrial gene content. Cells from all time points and driver lines (OK107 and MB247) were merged together with data from a-priori-labeled adult Kenyon cells into one Seurat object. Gene counts per cell were normalized to total read counts per cell (NormalizeData function, LogNormalize method). Variable genes were identified for each dataset independently (FindVariableFeatures function), followed by selection of 2000 genes that were repeatedly variable across datasets (SelectIntegrationFeatures function). To perform integration (IntegrateData function), this gene set was first filtered to exclude mitochondrial genes, heat shock proteins, transgenes (e.g. Gal4, GFP etc. – this was done to prevent transgenic variation driving the clustering), and ribosomal protein genes. Integrated data was then scaled and PCA was run with 50 principle components (ScaleData and RunPCA functions). Based on the ElbowPlot function, 40 dimensions were selected to cluster cells (RunUMAP, FindNeighbors, FindClusters functions). In FindClusters, clustering resolution was selected as 0.4 after testing a range of resolutions and using the R package “clustree” to assess stability scores of clusters at different resolutions (Zappia & Oshlack, 2018). Neurons, glia, and Kenyon cells were identified based on expression of known markers in the literature (Croset et al., 2018; G. Lee et al., 2000; Shih et al., 2019). As adult Kenyon cell classes were defined by *a priori* Gal4 classification of cells (described above), we could observe that the integrated data was coherent with the adult classification of clusters. In addition, libraries from the two drivers, MB247 and OK107, mixed well among each other and the transgenic backgrounds were not the key factor influencing the clustering (Figure 4C).

### Transcriptional analysis of cell surface and secreted molecules in Kenyon cells

To compare expression of CSS’s in olfactory Kenyon cells to visual Kenyon cells and non-Kenyon cell brain cells (Figure 5A), log-normalized read counts for each CSS in every cell were obtained by the Seurat function GetAssayData(slot="data", assays ="RNA"). The counts were then summed up for all the CSS’s to get a single value for each cell. This was done to take into account any difference in total number of reads across the 3 types of cell populations.

### Expression of Ig-SF and LRR molecules

Expression was measured by computing average expression of each gene in the Kenyon cell population and non-Kenyon cell populations using Seurat’s AverageExpression function (layer="data", assays ="RNA"). The resulted values are in a non-log space, therefore the expression was then normalized by log2 in the heatmaps displayed in Supplementary Figure 5-2.

We used UCell package (Andreatta & Carmona, 2021) to calculate the gene set enrichment scores at single-cell level for the following groups of genes: Ig-SF+LRR, Ig-SF, LRR, TF, and GPCR. Gene lists are included in the supplemental data. The ranking in UCell uses the Mann-Whitney U statistic.

### Re-analysis of published data: olfactory projection neurons

ScRNA-sequencing data for olfactory projection neurons featured in Figure 7 was acquired from Xie and colleagues (Xie et al., 2021). We performed differential expression analysis (FindMarkers function in Seurat, average log2-fold change > 5, adjusted p-value < 0.05) between MZ19-labeled PNs and other PNs to identify molecules uniquely and highly expressed in MZ19^+^ PNs across developmental time. Mz19 PNs were selected as clusters labeled DA1, VA1d, and DC3.

### Re-analysis of published data: fruitless neurons

The analysis of *fruitless* neurons was based on the publicly available dataset generated by the Arbeitman lab (Palmateer et al., 2023): *fru*+ neurons were purified at 48h APF and subjected to scRNAseq profiling using the 10X Genomics platform. We downloaded the processed dataset from NCBI GEO (GSE160370) and performed a re-analysis of the dataset starting from raw expression counts. The count matrix and metadata were extracted from the Seurat-object for the full dataset, including information on replicates, sex, percentage of mitochondrial transcripts, and cell type annotations. We cross-referenced different names for the same genes to build a consensus set of features for downstream analyses.

The reanalysis was carried out using Seurat v5.0.1 (Hao et al., 2024). We kept 24,478 cells with more than 1000 transcripts. The preprocessing and clustering were performed using the standard Seurat workflow: gene counts were normalized (function: NormalizeData); 2000 highly-variable genes (FindVariableFeatures) were scaled and used for PCA (functions: ScaleData/RunPCA); first 100 principle components were used for clustering (functions: FindNeighbors/FindClusters, resolution = 2) and tSNE projections (function: RunTSNE).

The initial analysis revealed an unexpected heterogeneity within many clusters that correspond to known cell types (e.g., KCs and visual system neurons). It was driven by activity-regulated genes (ARG), heat-shock proteins (HSP), and components of mitochondrial oxidation phosphorylation complexes (OXPHOS). These signatures are likely to represent unknown sources of biological and technical variability in this dataset (e.g., developmental pseudotime). We used UCell package (Andreatta & Carmona, 2021) to calculate the gene set enrichment scores for the following groups of genes: ARG (Hr38, sr, CG14186), HSP (Hsp67Ba, Hsp67Bc), OXPHOS (ATPsynbeta, ATPsyngamma). We regressed out these UCell scores together with the numbers of transcripts per cell and replicates (function: ScaleData) and repeated PCA and clustering analysis as described above.

The analysis revealed 132 clusters of neurons. The large transcriptionally distinct clusters are expected to represent abundant cell types with dozens to hundreds of copies per brain. The most abundant *fru+* cell types include γ KCs and cell types in the visual system. KC clusters were annotated based on the expression of known marker genes (Supplementary Figure 5-4). The visual system cell types were annotated using a published atlas of the visual system (Kurmangaliyev et al., 2020; Yoo et al., 2023). Normalized expression values were averaged for each cluster in *fru+* dataset and visual system neurons at 48h APF (atlas V1.1; https://zenodo.org/records/8111612). Next, we identified markers genes enriched in each fru+ cluster (function: FindAllMarkers, adjusted p-value < 0.01, fold-change > 8, detected in more than 50% of cells in the cluster). We used log-transformed expression values of the union of the top 10 marker genes (400 genes) to calculate Pearson’s correlation coefficients between clusters across datasets. The clusters with the best mutual match between datasets and Pearson’s r > 0.9 were considered the same cell types and used for label transfer (Supplementary Figure 5-3). We also verified matched clusters based on the expression patterns of marker genes (Supplementary Figure 5-4). In total, we annotated 3 KC clusters and 23 visual system clusters. Of those, 7 clusters are matched to neurons in the connectome (Dm3, Tm9, LC10a, LPC1, LLPC1, LLPC2, LLPC3); other clusters with the prefix “N” are yet to be mapped to morphological cell types. Clusters with the prefix “X” did not match clusters in the visual system atlas and we infer these are non-optic-lobe types. While the Palmateer dataset includes the ventral nerve cord, individual *fruitless* neuron types from the VNC are relatively rare, thus larger "X" clusters likely derive from the central brain.

## Selection of gene sets

### Cell Surface and Secreted Molecules (CSS’s)

This is a published list of 932 cell surface and secreted molecules (CSSs) with cell recognition properties (Kurusu et al., 2008).

### Ig-SF and LRR specificity molecules

To define a category of cell adhesion molecules from the much bigger list of CSS’s, we used gene classifications based on extracellular domain information for protein-coding genes from the *Drosophila melanogaster* extracellular domain database (Pei et al., 2018). This database has been used in work describing the transcriptional programs of synaptic specificity such as in the fly visual system (Kurmangaliyev et al., 2020). Cell adhesion molecules were defined from protein domains of Ig (132 genes) and LRR (72 genes).

### GPCRs

This is a list of 112 genes from Flybase; 111 of these were expressed in the Kenyon cell sequencing data, and 108 in the Palmateer data. 5 GPCRs that were also LRRs were excluded from this list: CG44153, Lgr1, Lgr3, Lgr4, rk.

### Transcription factors

This is a list of 628 transcription factors from Flybase. 613 of these were expressed in the Kenyon cell sequencing data, while 600 were expressed in the Palmateer data.

## MiMIC reporter validation of sequencing data

Expression pattern of CSS’s in the bulk- and single-cell sequencing datasets was cross-checked with expression patterns of MiMIC reporter lines for an immunoglobulin superfamily of proteins called dprs (defective proboscis response) and their binding partners, DIPs (dpr-interacting proteins). This group of genes was selected as they have known roles in synaptic matching and MiMIC lines were available for 14 of these genes (Barish et al., 2018; Bornstein et al., 2021; Carrillo et al., 2015; C. Xu et al., 2019). As a readout for reporter expression, we used the mushroom body pedunculus, a region where the Kenyon cell axons converge forming a stereotyped pattern with easily identifiable KC types, with early-born γ axons at the margin and latest-born αβ axons in the center of the pedunculus (transverse section) (Zhan et al., 2004).

## Brain dissection, immunostaining, and confocal imaging

For immunostaining, brains were dissected in external saline (108 mM NaCl, 5 mM KCl, 2 mM CaCl2, 8.2 mM MgCl2, 4 mM NaHCO3, 1 mM NaH2PO4, 5 mM trehalose, 10 mM sucrose, 5 mM HEPES pH7.5, osmolarity adjusted to 265 mOsm) for up to thirty minutes before being transferred to paraformaldehyde in PBS. All steps were performed in cell strainer baskets (caps of FACS tubes) in 24 well plates, with the brains in the baskets lifted from well to well to change solutions. Brains were fixed overnight at 4C in 1% PFA in PBS or for 25 minutes at room temperature in 4% PFA. In general, the 4% PFA method preserves neuronal structures better, while the 1% PFA method allows better antibody penetration. Some antibodies also work better with the 1% method. After same-day (4%) or overnight (1%) fixation, brains were washed 3x10’ in PBS supplemented with 0.1% triton-x-100 on a shaker at room temperature, blocked 1 hour in PBS, 0.1% triton, 4% Normal Goat Serum, and then incubated for at least two overnights in primary antibody solution, diluted in PBS, 0.1% triton, 4% Normal Goat Serum. Primary antibody was washed 3x10’ in PBS supplemented with 0.1% triton-x-100 on a shaker at room temperature, then brains were incubated in secondary antibodies for at least two overnights, diluted in PBS, 0.1% triton, 4% Normal Goat Serum. DAPI (1 microgram/mL) was included in secondary antibody mixes. Antibodies and concentrations can be found in the resources table.

Brains were mounted in 1x PBS, 90% glycerol supplemented with propyl gallate in binder reinforcement stickers sandwiched between two coverslips. Samples were stored at 4C in the dark prior to imaging. The coverslip sandwiches were taped to slides, allowing us to perform confocal imaging on one side of the brain and then flip over the sandwich to allow a clear view of the other side of the brain. Scanning confocal stacks were collected along the anterior-posterior axis on a Leica SP8 with 1 micrometer spacing in Z and ∼150nm axial pixel size, using a 40x 1.3 NA objective.

## Image analysis

### Analysis considerations

We used mixed-sex populations. Sex differences in the fly are well-documented, and anatomic and physiologic sex differences have not been observed in the mushroom body. Any brains that appeared damaged from dissections, or those with the mushroom body region obscured due to insufficient tracheal removal, were not included in the analysis.

Analysis was performed blind to genotype, and also blind to the goals of the experiment when possible, and quantitation of features on the anterior and posterior sides of the brain were recorded independent of one another and merged after all quantifications were completed.

### Calyx Volume

To measure the size of the mushroom body calyx, we used markers such as ChAT or brp to visualize the structure. We then measured the area of each slice of the calyx in Z by outlining in FIJI and using the ‘Measure’ command. Calyx volume was calculated as the sum of areas multiplied by the Z step size.

### Projection neuron bouton numbers

To count aggregate boutons, we used ChAT signal as previously described (Elkahlah et al., 2020). We counted as separate structures ChAT signals that were compact and appeared distinct from one another and that were 2+ micrometers in diameter. We found that boutons in slice 0 often appeared in slices −1 and +1 as well, but never in slices −2 or +2. To avoid over counting, we began at the most superficial slice in the stack where boutons were visible, and counted every other slice, i.e. every second micron.

To count boutons of subsets of PNs, we drove a fluorescent reporter under the control of VT033008, VT033006, or MZ19 and counted coherent and compact fluorescent signals, that overlapped with ChAT signals in order to distinguish true signal from background noise.

### Projection neuron cell numbers

To count PNs, we used genetically-encoded fluorescence in that group of cells. We counted labeled somata in every third slice in the stack (every third micron along the A-P axis), with reference to DAPI to distinguish individual cells from one another. As we did for boutons, in analyzing somata we initially determined that somata in slice 0 could also be seen in slices −2, –1, +1, and +2 but not in slice −3 or +3. To avoid double-counting, we therefore counted every third micron.

### Normalized Bruchpilot intensity

We measured Bruchpilot (Brp) intensity in the calyx as a readout of synaptic density. First, we identified the Z plane with the largest extent of the calyx in the A-P axis, and measured the mean flourescence value in a defined ROI region of 250^2^ pixels. The defined ROI was placed in the center of the calyx. For normalization, the calyx Brp signal was divided by Brp signal measured in a 75^2^ pixel ROI in an unmanipulated brain region, the protocerebral bridge. We chose this brain region as it was the closest to the calyx and was in the field of view in all calyx images taken.

### Normalized Kenyon cell fluorescence in bouton ROIs

We first defined ROIs in Fiji for each projection neuron bouton labeled by MZ19. Boutons were defined by using genetically-encoded fluorescence in that group of cells, overlapped with ChAT signal to distinguish true bouton signal from any background noise. In line with PN bouton counting methods described earlier, we only made bouton ROIs in every second slice (z step size was 1 micron). As a control we also made bouton ROIs not labeled by MZ19. These non-MZ19 boutons were nearby MZ19+ boutons. We also matched the number of non-MZ19 bouton ROIs to the number of MZ19+ bouton ROIs for each z-slice of the calyx. For each bouton ROI, we then measured the mean fluorescence value in the Kenyon cell channel and normalized it to the mean fluorescence in the Kenyon cell channel in the calyx ROI defined by ChAT immunoreactivity.

### Statistical considerations

Brains were prepared for imaging in batches of 5–10. In initial batches, we assessed the variability of the manipulation, and used this variability to determine how many batches to analyze so as to obtain enough informative samples. Genotypes or conditions being compared with one another were always prepared for staining together and imaged interspersed with one another to equalize batch effects. We excluded from analysis samples with overt physical damage to the cells or structures being measured.

Statistical tests applied are mentioned in each figure legend along with the p-value significance. Normality of data was assessed with Shapiro test. When both distributions were normal (p>0.05) we used two-sided Student’s T-test. When one or both distributions were non-normal (p<=0.05) we used the two-sided Wilcoxon rank sum test. P values are also included in the figure.

To communicate our findings in the simplest and most complete way, we have displayed each data point for each sample to allow readers to assess effect size and significance directly. Effect size was also calculated using Cohen’s D using a pooled standard deviation. Statistical analyses were performed in R.

## Data availability

We used data from the Hemibrain connectome that is available to the public under a Creative Commons license at https://neuprint.janelia.org/. Annotated tables of synapses, projection neurons, projection neuron boutons and branch points, Kenyon cells, and Kenyon cell claws are provided in Supplementary Information. R scripts used to annotate and analyze Hemibrain connectome data can be found at https://github.com/emmamtk/ThorntonKolbeAhmed2024 which will be made public on publication. Bulk RNA sequencing of Kenyon cells and projection neurons will be deposited into the Gene Expression Omnibus database by the time of publication. scRNAseq of Kenyon cells at developmental time points will be deposited into the Gene Expression Omnibus database by the time of publication. scRNAseq of adult Kenyon cells was shared prior to publication by author BS and is being prepared for publication separately.

---

**Supplementary Figure 1.**
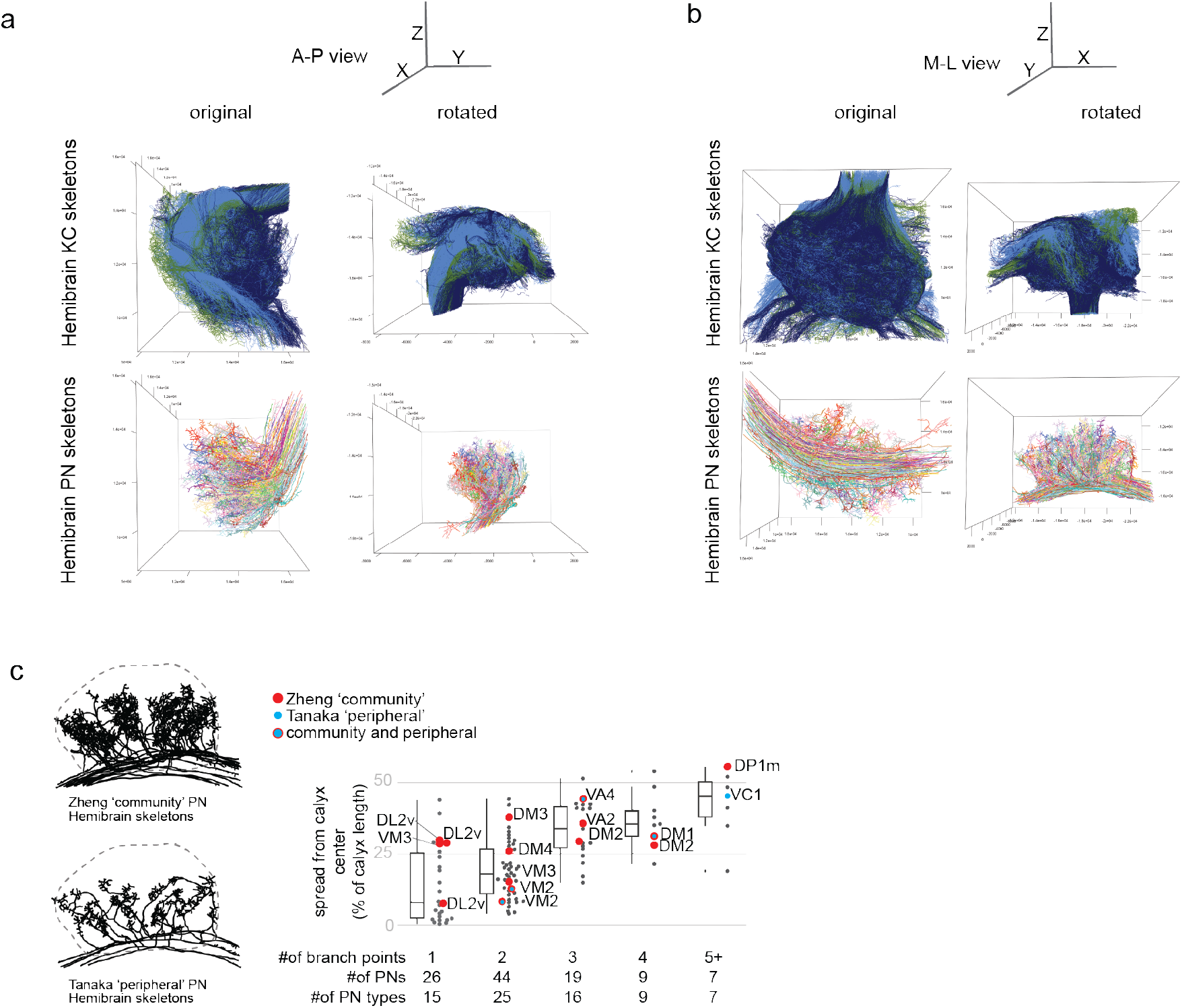
**(a-b)** A-P view (a) and M-L view (b) of KCs (top) and PNs (bottom) before (left) and after (right) 285 degree rotation around the X axis, and 200 degree rotation about the Y axis such that the calyx axons of the mALT were perpendicular to the X axis. **(c)**left: The 15 Hemibrain skeletons of PNs in the overconvergent community in Zheng 2022 (DL2v, VM3,DM3, DM4, VM2,VA4,VA2, DM2,DM1,DP1m)(top) and the 8 Hemibrain skeletons of the overlapping set of PNs labeled as peripheral calyx innervators by Tanaka 2004 (VM2,VA4,DM1,VC1). Skeletons are displayed M-L with a dotted outline of the calyx. Plot from 1j showing maximum spread from calyx center by number of collaterals with Zheng community PNs highlighted in red, Tanaka peripheral PNs highlighted in blue, and those in both groups highlighted in blue surrounded by red. These PNs mostly make several collaterals and thus spread into the periphery or are notably peripheralized for PNs with the same number of collaterals with the exception of VM2.

**Supplementary Figure 2.**
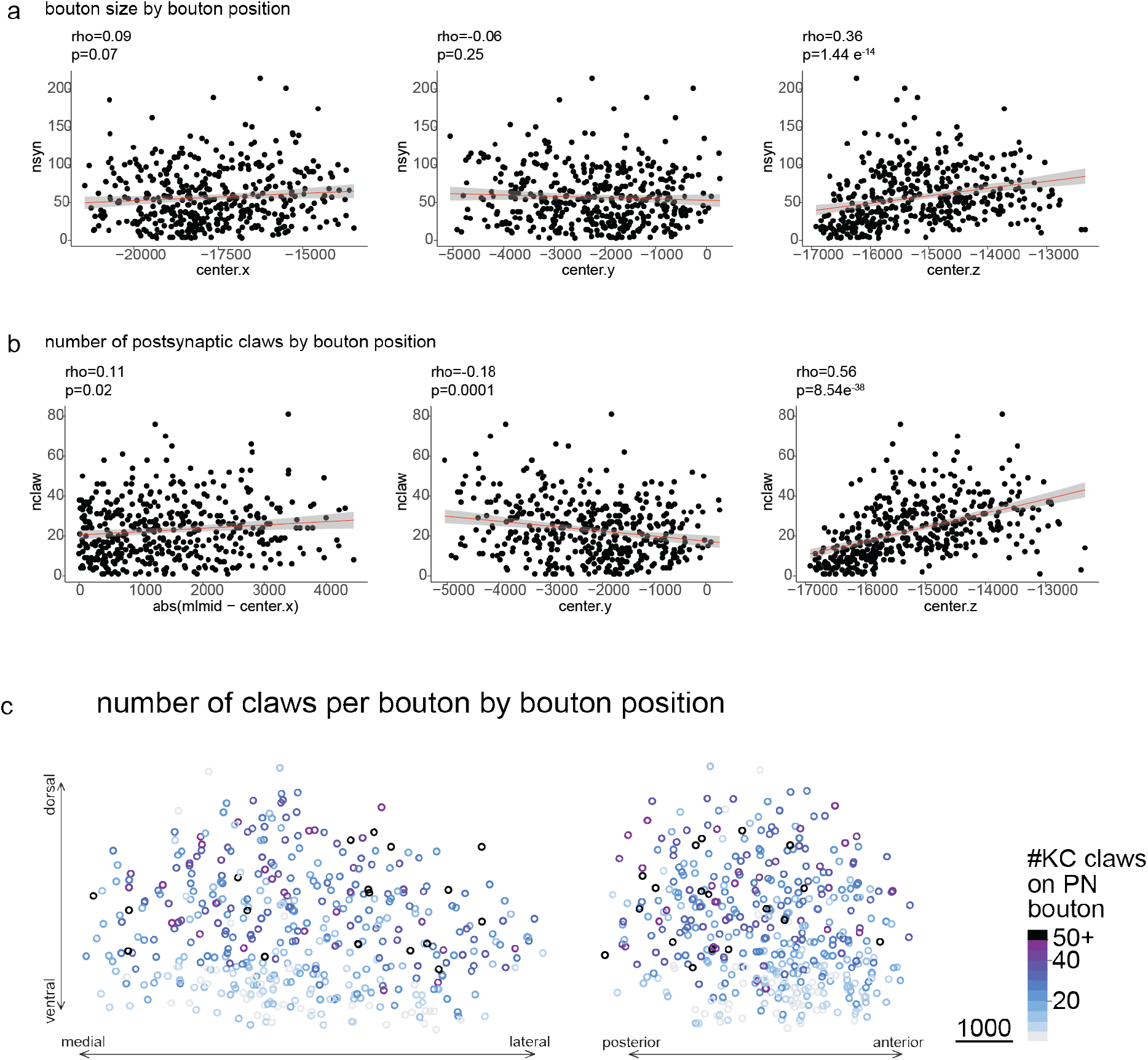
**(a)**Bouton size as measured by number of presynapses is invariable over the M-L and A-P axis and weakly correlated with D-V position. M-L(Spearman’s rank correlation, rho=0.09, p=0.07), A-P(Spearman’s rank correlation, rho=0.06, p=0.25),D-V(Spearman’s rank correlation, rho=0.36, p=1.44e-14) **(b)**Number of claws per bouton is weakly correlated with bouton position along the M-L (Spearman’s rank correlation, rho=0.11, p=0.02) and A-P (Spearman’s rank correlation, rho=- 0.18, p=0.0001) axes but more strongly correlated with bouton D-V position(Spearman’s rank correlation, rho=0.56, p=8.54e-38). **(c)** bouton centers colored by number of claws postsynaptic to that bouton.

**Supplementary Figure 4.**
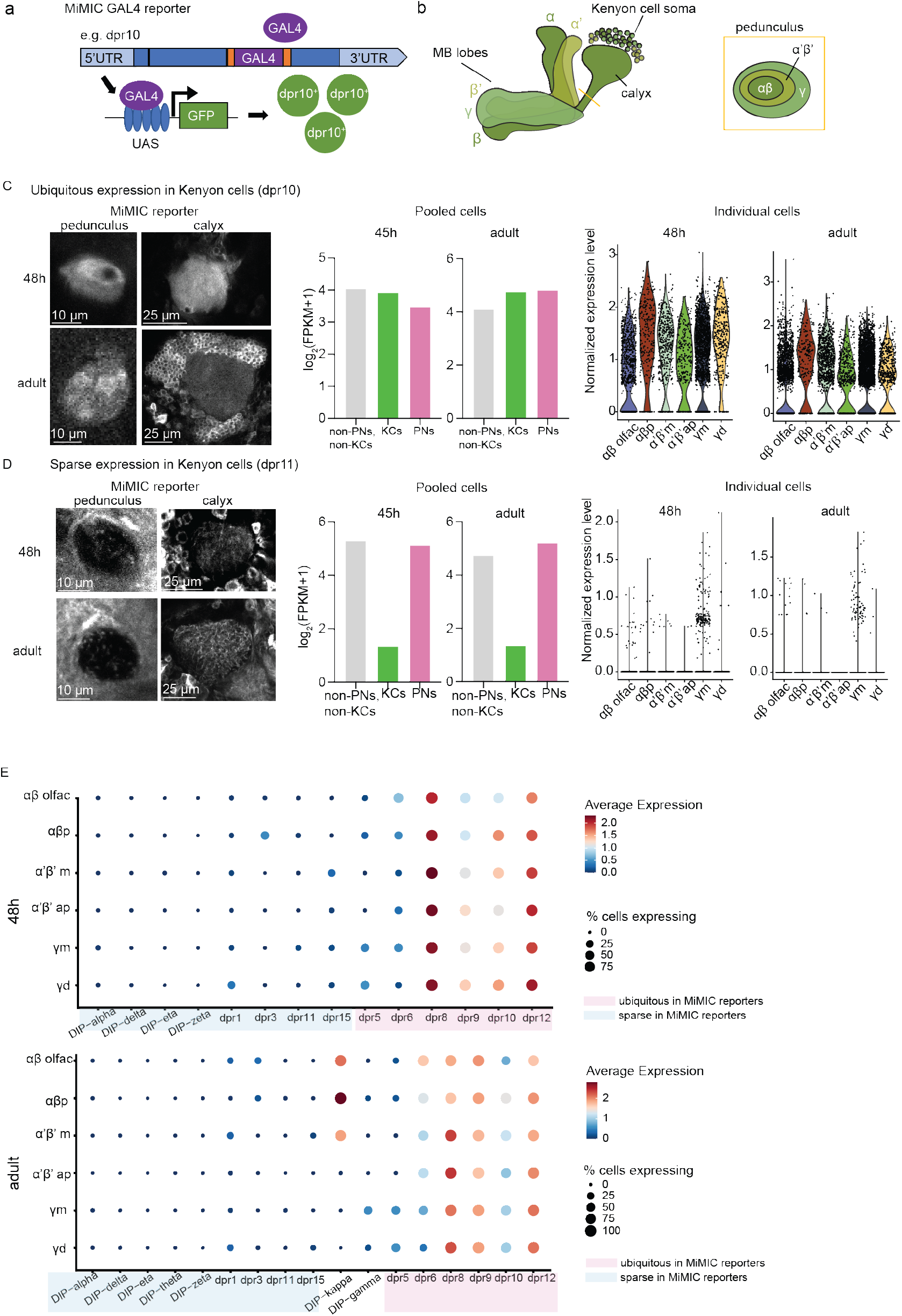
**(a)** Schematic of MiMIC-Gal4 reporter inserted into the coding intron of the gene of interest e.g. dpr10, resulting in dpr10 expressing cells to become GFP-positive. **(b)** Right: Mushroom body schematic with the 3 major KC classes labeled. A transverse section where axons of the 3 classes converge (yellow line) shows the pedunculus(boxed inset); readout for gene expression in KCs. **(c)** Expression of a gene (dpr10) ubiquitously expressed in 48hAPF and adult Kenyon cells in a MiMIC- Gal4 reporter line (left), bulk RNA-sequencing data (middle), and single-cell RNA-sequencing (right). Bulk RNA-seq: normalized FPKM values, scRNA-seq: log-normalized expression where expression of 3 corresponds to ∼20 reads per cell. **(d)** Expression of a gene (dpr11) sparsely expressed in 48hAPF and adult Kenyon cells in a MiMIC-Gal4 reporter line (left), bulk RNA-sequencing data (middle), and single-cell RNA-sequencing (right). Bulk RNA-seq: normalized FPKM values, scRNA-seq: log-normalized expression where expression of 3 corresponds to ∼20 reads per cell. **(e)** Dot plot showing average expression and percentage of cells expressing the gene (expression > 0) in scRNA-seq data of all 14 MiMIC-Gal4 reporters tested from the DIP/dpr gene family, at both the 48h APF and adult stages. Dot size represents percentage of cells the gene is expressed in, and the color represents log-normalized average expression. Pink: genes that were ubiquitous in MiMIC-Gal4 reporters, blue: sparse in reporters. The two unhighlighted genes (DIP- kappa, DIP-γ) were only tested in adults and were specific to αβ and γ KCs respectively.

**Supplementary Figure 5-1.**
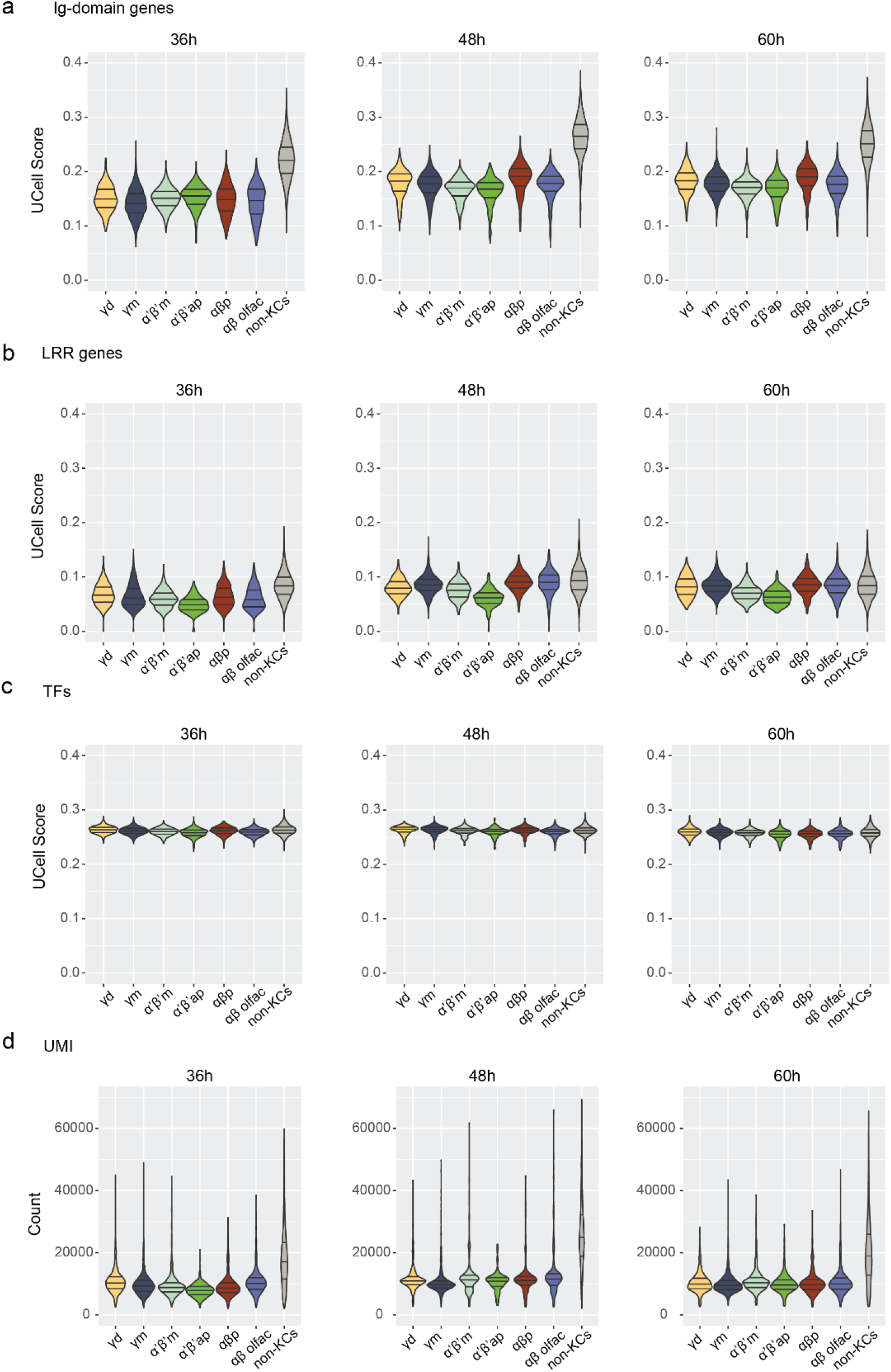
**(a-c)** UCell scores of Ig-domain genes, LRR-domain genes, and transcription factors (TF) in our dataset. Violins show distributions of UCell values for individual cells. Solid lines: 25%, 50% (median), and 75% quartiles for each violin. **(d)** Distributions of number of UMIs per cell. Lower UMIs in Kenyon cells reflect lower total RNA content per cell.

**Supplementary Figure 5-2.**
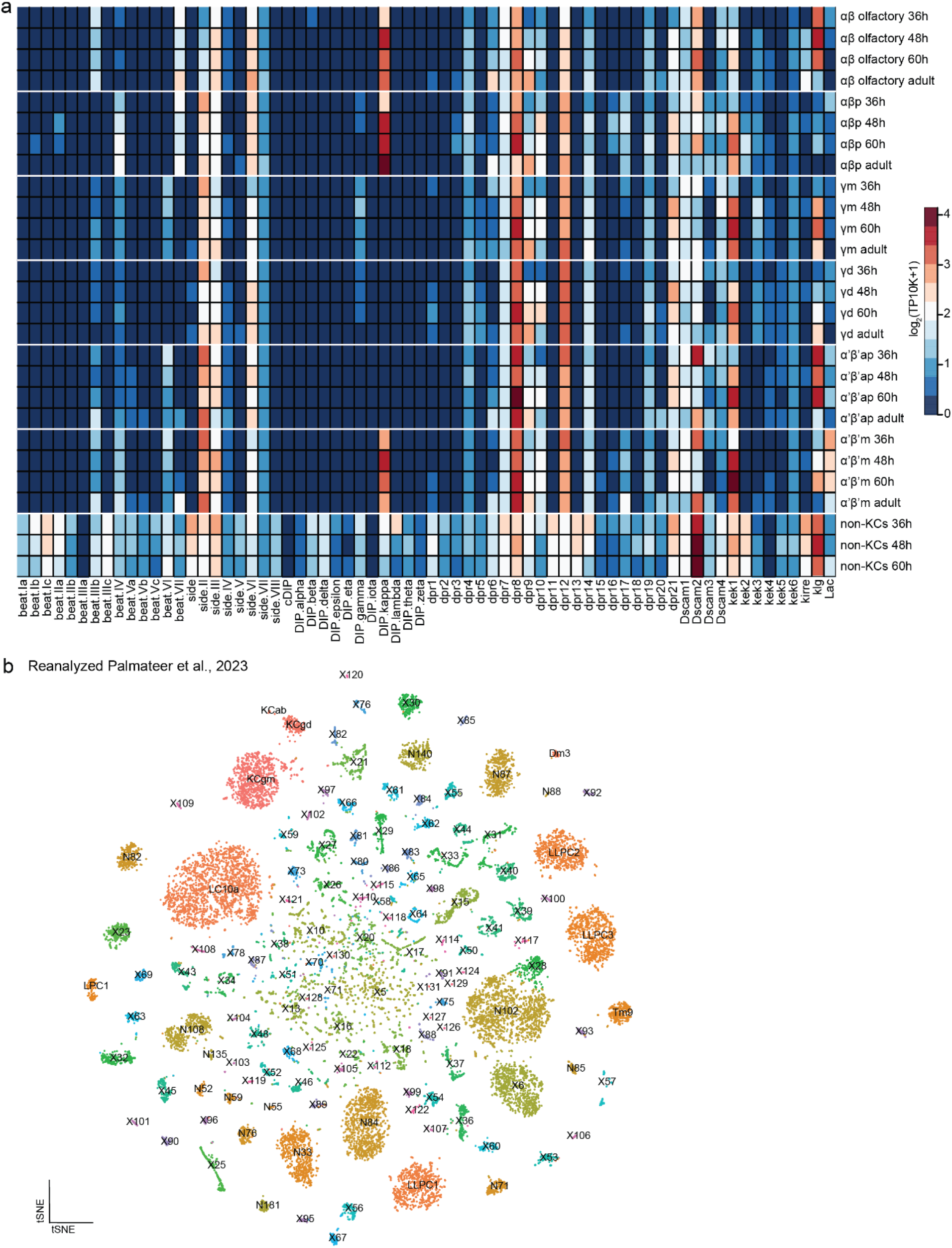
**(a)** Average expression of Ig-SF molecules in individual Kenyon cell types across developmental time, as well as in a mélange of other brain cells. **(b)** tSNE of Palmateer et al., 2023 dataset of *fruitless*-expressing neurons. The visual system neurons were annotated based on cross-correlation analysis with the visual system atlas from Yoo et al (see Supplementary Figure 5-3). Kenyon cells are annotated based on the expression of dac, ey, and Mef2. Clusters tagged “N” have one-to-one matches with transcriptional cell types in the visual system atlas, but have not been matched yet to cell types in the connectome. We infer that the remaining clusters, tagged “X,” are from the central brain or ventral nerve cord, as these do not find a match in the visual system.

**Supplementary Figure 5-3.**
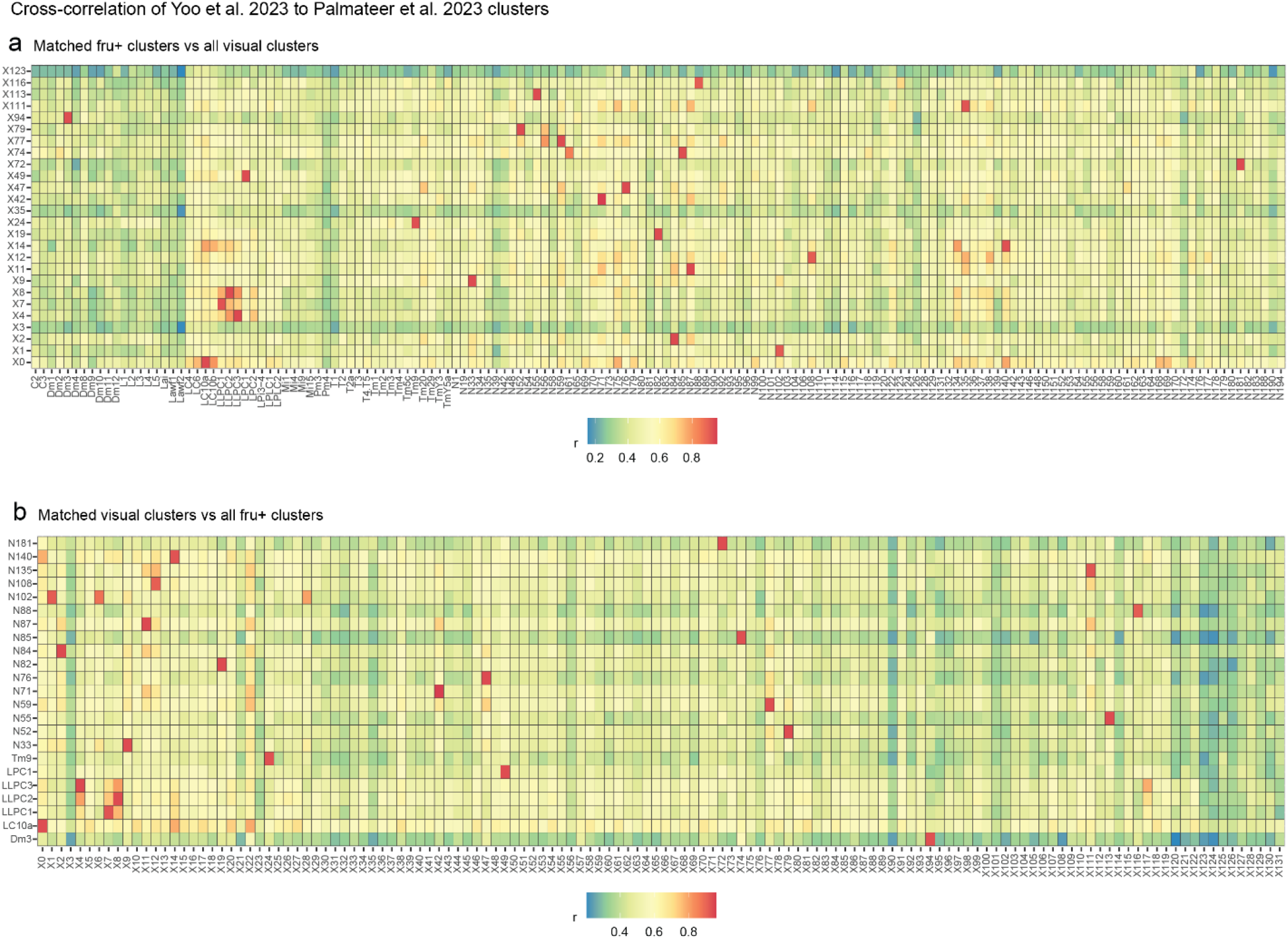
**(a,b)** Cross-correlation analysis of Palmateer clusters and our optic lobe atlas from Yoo et al., 2023. Heatmaps show Pearson’s correlation coefficients (r) between expression profiles of marker genes in clusters from Palmateer and Yoo datasets. Data is shown for 23 clusters with the best mutual matches between two datasets.

**Supplementary Figure 5-4.**
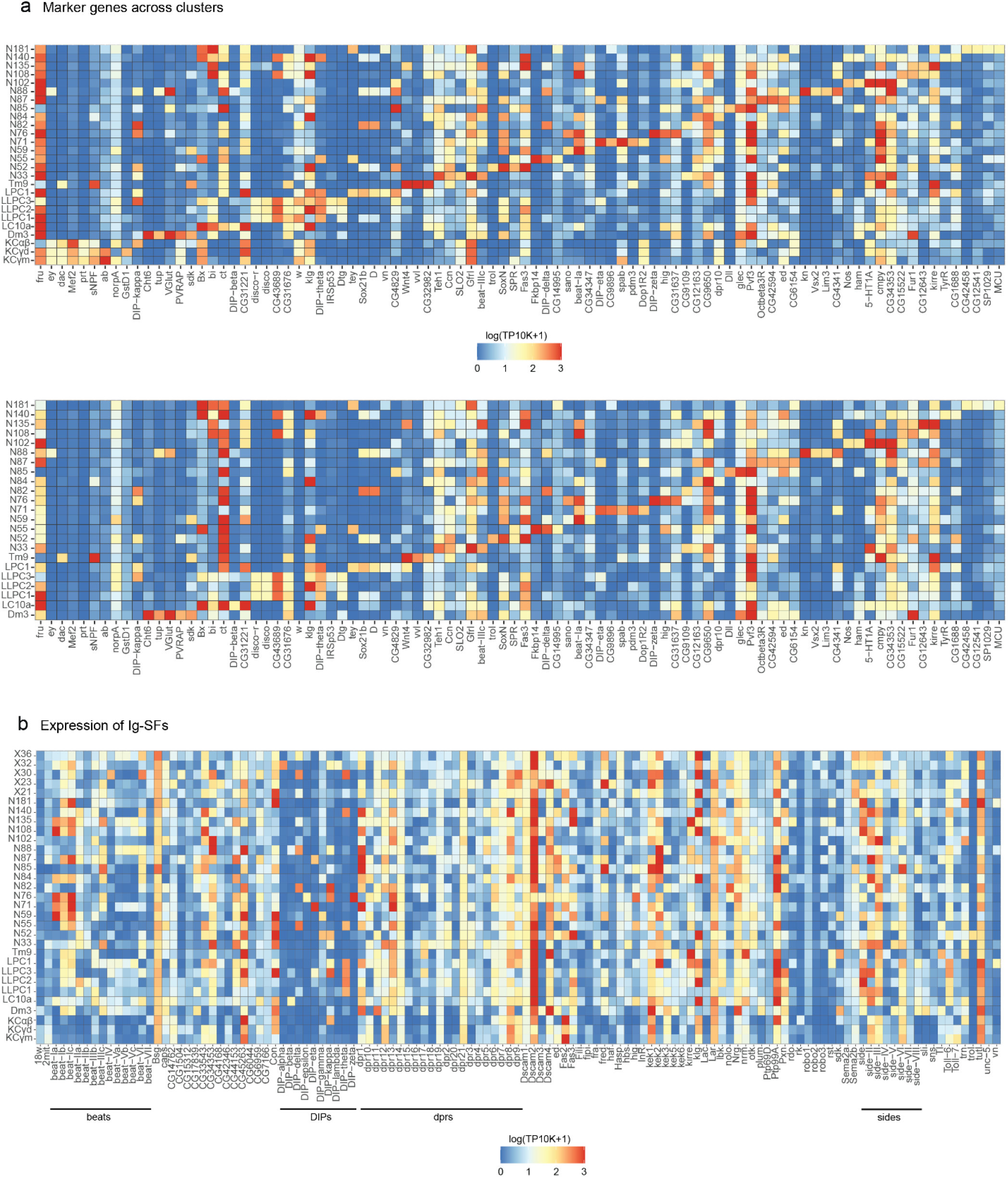
**(a)** Discriminative markers for cell types with one-to-one matches between Yoo et al., and Palmateer et al. Markers for Kenyon cell clusters are also shown. **(b)** Average expression of Ig-SF molecules in select cell types and clusters in Palmateer et al. dataset.

**Supplementary Figure 5-5.**
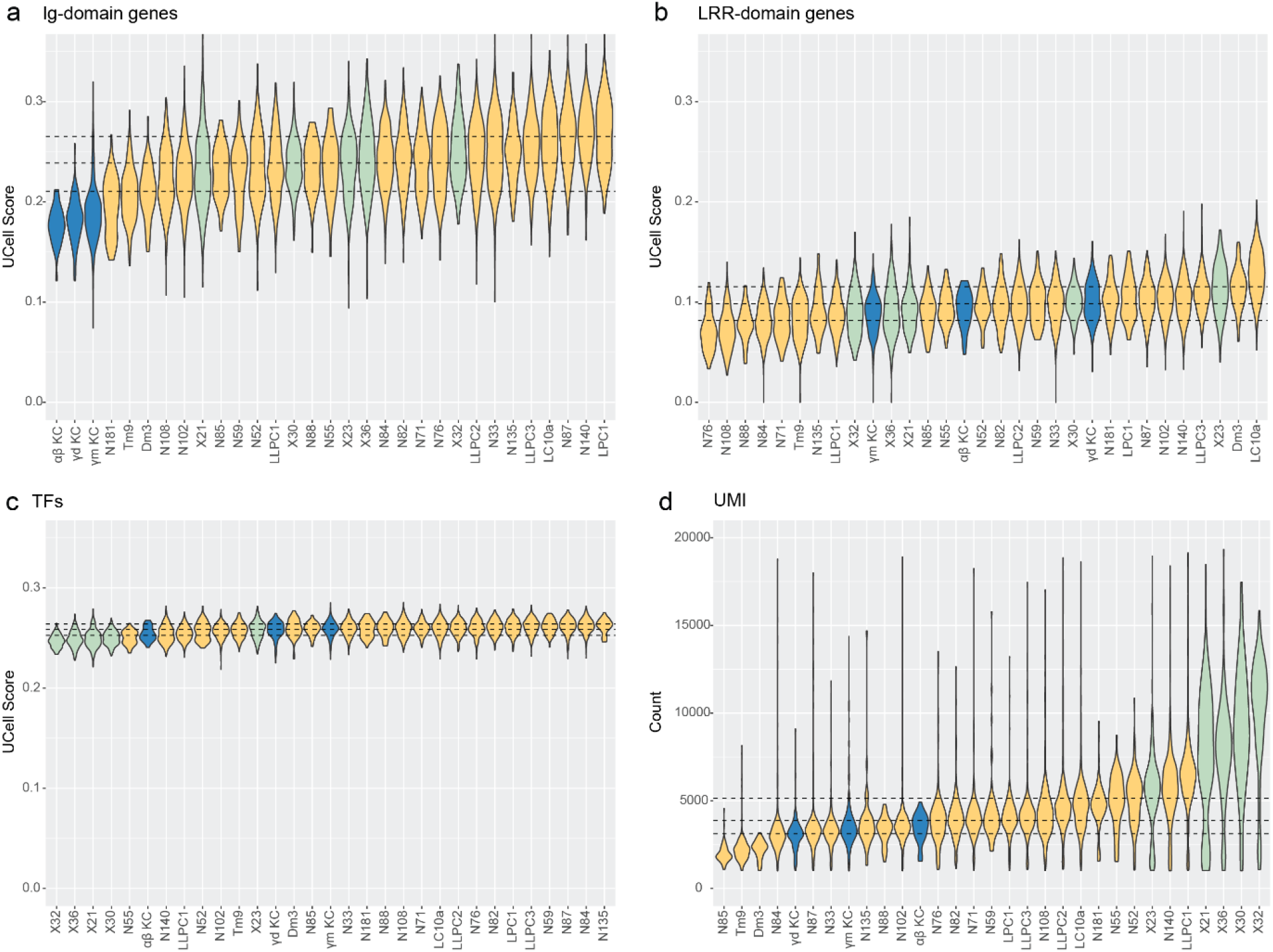
**(a-c)** UCell scores of Ig-domain, LRR-domain and TF gene sets for select cell types and clusters from Figure 5d. Violins show distributions of UCell values for individual cells. Dashed lines: 25%, 50% (median), and 75% quartiles across the entire dataset. Clusters are ordered by UCell median. Additional gene sets are shown in Figure 5e,f. **(d)** Distributions of number of UMIs per cell. Clusters are ordered by median UMIs. Central brain neurons (“X” clusters, green) are generally larger and have higher RNA content, while Kenyon cells and optic lobe neurons (blue, yellow) are smaller and have lower RNA content.

## References

Ahmed, M., Rajagopalan, A. E., Pan, Y., Li, Y., Williams, D. L., Pedersen, E. A., Thakral, M., Previero, A., Close, K. C., Christoforou, C. P., Cai, D., Turner, G. C., & Clowney, E. J. (2023). Input density tunes Kenyon cell sensory responses in the Drosophila mushroom body. Current Biology: CB, 33(13), 2742–2760.e12.

Albus, J. S. (1971). A theory of cerebellar function. Mathematical Biosciences, 10(1), 25–61.

Andreatta, M., & Carmona, S. J. (2021). UCell: Robust and scalable single-cell gene signature scoring. Computational and Structural Biotechnology Journal, 19, 3796–3798.

Aresta-Branco, F., Erben, E., Papavasiliou, F. N., & Stebbins, C. E. (2019). Mechanistic Similarities between Antigenic Variation and Antibody Diversification during Trypanosoma brucei Infection. Trends in Parasitology, 35(4), 302–315.

Aso, Y., Grübel, K., Busch, S., Friedrich, A. B., Siwanowicz, I., & Tanimoto, H. (2009). The mushroom body of adult Drosophila characterized by GAL4 drivers. Journal of Neurogenetics, 23(1–2), 156–172.

Aso, Y., Hattori, D., Yu, Y., Johnston, R. M., Iyer, N. A., Ngo, T. T. B., Dionne, H., Abbott, L. F., Axel, R., Tanimoto, H., & Rubin, G. M. (2014). The neuronal architecture of the mushroom body provides a logic for associative learning. ELife, 3, e04577.

Barak, O., Rigotti, M., & Fusi, S. (2013). The sparseness of mixed selectivity neurons controls the generalization-discrimination trade-off. The Journal of Neuroscience: The Official Journal of the Society for Neuroscience, 33(9), 3844–3856.

Barish, S., Nuss, S., Strunilin, I., Bao, S., Mukherjee, S., Jones, C. D., & Volkan, P. C. (2018). Combinations of DIPs and Dprs control organization of olfactory receptor neuron terminals in Drosophila. PLOS Genetics, 14, 1–33.

Bates, Alexander S., Schlegel, P., Roberts, R. J. V., Drummond, N., Tamimi, I. F. M., Turnbull, R., Zhao, X., Marin, E. C., Popovici, P. D., Dhawan, S., Jamasb, A., Javier, A., Serratosa Capdevila, L., Li, F., Rubin, G. M., Waddell, S., Bock, D. D., Costa, M., & Jefferis, G. S. X. E. (2020). Complete Connectomic Reconstruction of Olfactory Projection Neurons in the Fly Brain. Current Biology: CB, 30(16), 3183–3199.e6.

Bates, Alexander Shakeel, Manton, J. D., Jagannathan, S. R., Costa, M., Schlegel, P., Rohlfing, T., & Jefferis, G. S. (2020). The natverse, a versatile toolbox for combining and analysing neuroanatomical data. ELife, 9. 10.7554/eLife.53350

Bekkers, J. M., & Stevens, C. F. (1991). Excitatory and inhibitory autaptic currents in isolated hippocampal neurons maintained in cell culture. Proceedings of the National Academy of Sciences of the United States of America, 88(17), 7834–7838.

Bengtsson, F., & Jörntell, H. (2009). Sensory transmission in cerebellar granule cells relies on similarly coded mossy fiber inputs. Proceedings of the National Academy of Sciences of the United States of America, 106(7), 2389–2394.

Bischof, J., Björklund, M., Furger, E., Schertel, C., Taipale, J., & Basler, K. (2013). A versatile platform for creating a comprehensive UAS-ORFeome library in Drosophila. *Development (Cambridge*, England), 140(11), 2434–2442.

Bornstein, B., Meltzer, H., Adler, R., Alyagor, I., Berkun, V., Cummings, G., Reh, F., Keren-Shaul, H., David, E., Riemensperger, T., & Schuldiner, O. (2021). Transneuronal Dpr12/DIP-δ interactions facilitate compartmentalized dopaminergic innervation of Drosophila mushroom body axons. The EMBO Journal, 40(12), e105763.

Brovero, S. G., Fortier, J. C., Hu, H., Lovejoy, P. C., Newell, N. R., Palmateer, C. M., Tzeng, R.-Y., Lee, P.-T., Zinn, K., & Arbeitman, M. N. (2021). Investigation of Drosophila fruitless neurons that express Dpr/DIP cell adhesion molecules. ELife, 10. 10.7554/eLife.63101

Brovkina, M. V., Duffié, R., Burtis, A. E. C., & Clowney, E. J. (2021). Fruitless decommissions regulatory elements to implement cell-type-specific neuronal masculinization. PLoS Genetics, 17(2), e1009338.

Cachero, S., Gkantia, M., Bates, A. S., Frechter, S., Blackie, L., McCarthy, A., Sutcliffe, B., Strano, A., Aso, Y., & Jefferis, G. S. X. E. (2020). BAcTrace, a tool for retrograde tracing of neuronal circuits in Drosophila. Nature Methods, 17(12), 1254–1261.

Caron, S. J. C., Ruta, V., Abbott, L. F., & Axel, R. (2013). Random convergence of olfactory inputs in the Drosophila mushroom body. Nature, 497(7447), 113–117.

Carrillo, R. A., Özkan, E., Menon, K. P., Nagarkar-Jaiswal, S., Lee, P. T., Jeon, M., Birnbaum, M. E., Bellen, H. J., Garcia, K. C., & Zinn, K. (2015). Control of Synaptic Connectivity by a Network of Drosophila IgSF Cell Surface Proteins. Cell, 163(7), 1770–1782.

Cayco-Gajic, N. A., Clopath, C., & Silver, R. A. (2017). Sparse synaptic connectivity is required for decorrelation and pattern separation in feedforward networks. Nature Communications, 8(1), 1116.

Cayco-Gajic, N. A., & Silver, R. A. (2019). Re-evaluating Circuit Mechanisms Underlying Pattern Separation. Neuron, 101(4), 584–602.

Chabrol, F. P., Arenz, A., Wiechert, M. T., Margrie, T. W., & DiGregorio, D. A. (2015). Synaptic diversity enables temporal coding of coincident multisensory inputs in single neurons. Nature Neuroscience, 18(5), 718–727.

Clements, J., Dolafi, T., Umayam, L., Neubarth, N. L., Berg, S., Scheffer, L. K., & Plaza, S. M. (2020). neuPrint: Analysis Tools for EM Connectomics. In bioRxiv (p. 2020.01.16.909465). 10.1101/2020.01.16.909465

Connolly, J. B., Roberts, I. J., Armstrong, J. D., Kaiser, K., Forte, M., Tully, T., & O’Kane, C. J. (1996). Associative learning disrupted by impaired Gs signaling in Drosophila mushroom bodies. Science (New York, N.Y.), 274(5295), 2104–2107.

Croset, V., Treiber, C. D., & Waddell, S. (2018). Cellular diversity in the Drosophila midbrain revealed by single-cell transcriptomics. ELife, 7, 1–31.

De Belle, J. S., & Heisenberg, M. (1994). Associative odor learning in Drosophila abolished by chemical ablation of mushroom bodies. Science, 263(5147), 692–695.

Di Bella, D. J., Habibi, E., Stickels, R. R., Scalia, G., Brown, J., Yadollahpour, P., Yang, S. M., Abbate, C., Biancalani, T., Macosko, E. Z., Chen, F., Regev, A., & Arlotta, P. (2021). Molecular logic of cellular diversification in the mouse cerebral cortex. Nature, 595(7868), 554–559.

Drotos, A. C., & Roberts, M. T. (2024). Identifying neuron types and circuit mechanisms in the auditory midbrain. Hearing Research, 442, 108938.

Edelman, G. M. (1989). The remembered present: A biological theory of consciousness. The Remembered Present: A Biological Theory of Consciousness. https://psycnet.apa.org/record/1990-97063-000

Eichler, K., Li, F., Litwin-Kumar, A., Park, Y., Andrade, I., Schneider-Mizell, C. M., Saumweber, T., Huser, A., Eschbach, C., Gerber, B., Fetter, R. D., Truman, J. W., Priebe, C. E., Abbott, L. F., Thum, A. S., Zlatic, M., & Cardona, A. (2017). The complete connectome of a learning and memory centre in an insect brain. Nature, 548(7666), 175–182.

Elkahlah, N. A., Rogow, J. A., Ahmed, M., & Clowney, E. J. (2020). Presynaptic developmental plasticity allows robust sparse wiring of the drosophila mushroom body. ELife, 9. 10.7554/eLife.52278

Ellis, K. E., Smihula, H., Ganguly, I., Vigato, E., Bervoets, S., Auer, T. O., Benton, R., Litwin-Kumar, A., & Caron, S. J. C. (2023). Evolution of connectivity architecture in the Drosophila mushroom body. In bioRxiv (p. 2023.02.10.528036). 10.1101/2023.02.10.528036

Farris, S. M. (2011). Are mushroom bodies cerebellum-like structures? Arthropod Structure & Development, 40(4), 368–379.

Farris, S. M. (2015). Evolution of brain elaboration. Philosophical Transactions of the Royal Society of London. Series B, Biological Sciences, 370(1684), 20150054.

Ganguly, I., Heckman, E. L., Litwin-Kumar, A., Clowney, E. J., & Behnia, R. (2024). Diversity of visual inputs to Kenyon cells of the Drosophila mushroom body. Nature Communications, 15, 5698.

Gayoso, A., Lopez, R., Xing, G., Boyeau, P., Valiollah Pour Amiri, V., Hong, J., Wu, K., Jayasuriya, M., Mehlman, E., Langevin, M., Liu, Y., Samaran, J., Misrachi, G., Nazaret, A., Clivio, O., Xu, C., Ashuach, T., Gabitto, M., Lotfollahi, M., … Yosef, N. (2022). A Python library for probabilistic analysis of single-cell omics data. Nature Biotechnology, 40(2), 163–166.

Gilmer, J. I., & Person, A. L. (2017). Morphological Constraints on Cerebellar Granule Cell Combinatorial Diversity. The Journal of Neuroscience: The Official Journal of the Society for Neuroscience, 37(50), 12153–12166.

Grabe, V., Baschwitz, A., Dweck, H. K. M., Lavista-Llanos, S., Hansson, B. S., & Sachse, S. (2016). Elucidating the Neuronal Architecture of Olfactory Glomeruli in the Drosophila Antennal Lobe. Cell Reports, 16(12), 3401–3413.

Grueber, W. B., & Sagasti, A. (2010). Self-avoidance and tiling: Mechanisms of dendrite and axon spacing. Cold Spring Harbor Perspectives in Biology, 2(9), a001750.

Gruntman, E., & Turner, G. C. (2013). Integration of the olfactory code across dendritic claws of single mushroom body neurons. Nature Neuroscience, 16(12), 1821–1829.

Han, D. D., Stein, D., & Stevens, L. M. (2000). Investigating the function of follicular subpopulations during Drosophila oogenesis through hormone-dependent enhancer-targeted cell ablation. Development (Cambridge, England), 127(3), 573–583.

Hao, Y., Hao, S., Andersen-Nissen, E., Mauck, W. M., 3rd, Zheng, S., Butler, A., Lee, M. J., Wilk, A. J., Darby, C., Zager, M., Hoffman, P., Stoeckius, M., Papalexi, E., Mimitou, E. P., Jain, J., Srivastava, A., Stuart, T., Fleming, L. M., Yeung, B., … Satija, R. (2021). Integrated analysis of multimodal single-cell data. Cell, 184(13), 3573–3587.e29.

Hao, Y., Stuart, T., Kowalski, M. H., Choudhary, S., Hoffman, P., Hartman, A., Srivastava, A., Molla, G., Madad, S., Fernandez-Granda, C., & Satija, R. (2024). Dictionary learning for integrative, multimodal and scalable single-cell analysis. Nature Biotechnology, 42(2), 293–304.

Hassan, B. A., & Hiesinger, P. R. (2015). Beyond Molecular Codes: Simple Rules to Wire Complex Brains. Cell, 163(2), 285–291.

Hayashi, T. T., MacKenzie, A. J., Ganguly, I., Ellis, K. E., Smihula, H. M., Jacob, M. S., Litwin-Kumar, A., & Caron, S. J. C. (2022). Mushroom body input connections form independently of sensory activity in Drosophila melanogaster. Current Biology: CB, 0(0). 10.1016/j.cub.2022.07.055

Hiesinger, P. R. (2021). Brain wiring with composite instructions. *BioEssays: News and Reviews in Molecular*, Cellular and Developmental Biology, 43(1), e2000166.

Hobbs, M. J., & Young, J. Z. (1973). A cephalopod cerebellum. Brain Research, 55(2), 424–430.

Hong, W., Mosca, T. J., & Luo, L. (2012). Teneurins instruct synaptic partner matching in an olfactory map. Nature, 484(7393), 201–207.

Huang, C.-C., Sugino, K., Shima, Y., Guo, C., Bai, S., Mensh, B. D., Nelson, S. B., & Hantman, A. W. (2013). Convergence of pontine and proprioceptive streams onto multimodal cerebellar granule cells. ELife, 2, e00400.

Ishikawa, T., Shimuta, M., & Häusser, M. (2015). Multimodal sensory integration in single cerebellar granule cells in vivo. ELife, 4. 10.7554/eLife.12916

Ito, K., Awano, W., Suzuki, K., Hiromi, Y., & Yamamoto, D. (1997). The Drosophila mushroom body is a quadruple structure of clonal units each of which contains a virtually identical set of neurones and glial cells. Development (Cambridge, England), 124(4), 761–771.

Ito, M. (1970). Neurophysiological aspects of the cerebellar motor control system. International Journal of Neurology, 7(2), 162–176.

Jefferis, G. S. X. E., Potter, C. J., Chan, A. M., Marin, E. C., Rohlfing, T., Maurer, C. R., Jr, & Luo, L. (2007). Comprehensive maps of Drosophila higher olfactory centers: spatially segregated fruit and pheromone representation. Cell, 128(6), 1187–1203.

Jefferis, G. S. X. E., Vyas, R. M., Berdnik, D., Ramaekers, A., Stocker, R. F., Tanaka, N. K., Ito, K., & Luo, L. (2004). Developmental origin of wiring specificity in the olfactory system of Drosophila. Development, 131(1), 117–130.

Jenett, A., Rubin, G. M., Ngo, T.-T. B., Shepherd, D., Murphy, C., Dionne, H., Pfeiffer, B. D., Cavallaro, A., Hall, D., Jeter, J., Iyer, N., Fetter, D., Hausenfluck, J. H., Peng, H., Trautman, E. T., Svirskas, R. R., Myers, E. W., Iwinski, Z. R., Aso, Y., … Zugates, C. T. (2012). A GAL4-driver line resource for Drosophila neurobiology. Cell Reports, 2(4), 991–1001.

Jortner, R. A., Farivar, S. S., & Laurent, G. (2007). A simple connectivity scheme for sparse coding in an olfactory system. The Journal of Neuroscience: The Official Journal of the Society for Neuroscience, 27(7), 1659–1669.

Kasthuri, N., Hayworth, K. J., Berger, D. R., Schalek, R. L., Conchello, J. A., Knowles-Barley, S., Lee, D., Vázquez-Reina, A., Kaynig, V., Jones, T. R., Roberts, M., Morgan, J. L., Tapia, J. C., Seung, H. S., Roncal, W. G., Vogelstein, J. T., Burns, R., Sussman, D. L., Priebe, C. E., … Lichtman, J. W. (2015). Saturated Reconstruction of a Volume of Neocortex. Cell, 162(3), 648–661.

Kennedy, A., Wayne, G., Kaifosh, P., Alviña, K., Abbott, L. F., & Sawtell, N. B. (2014). A temporal basis for predicting the sensory consequences of motor commands in an electric fish. Nature Neuroscience, 17(3), 416–422.

Kiral, F. R., Linneweber, G. A., Mathejczyk, T., Georgiev, S. V., Wernet, M. F., Hassan, B. A., von Kleist, M., & Hiesinger, P. R. (2020). Autophagy-dependent filopodial kinetics restrict synaptic partner choice during Drosophila brain wiring. Nature Communications, 11(1), 1325.

Kunz, T., Kraft, K. F., Technau, G. M., & Urbach, R. (2012). Origin of Drosophila mushroom body neuroblasts and generation of divergent embryonic lineages. *Development (Cambridge*, England), 139(14), 2510–2522.

Kurmangaliyev, Y. Z., Yoo, J., Valdes-Aleman, J., Sanfilippo, P., & Zipursky, S. L. (2020). Transcriptional Programs of Circuit Assembly in the Drosophila Visual System. Neuron, 108(6), 1045–1057.e6.

Kurusu, M., Cording, A., Taniguchi, M., Menon, K., Suzuki, E., & Zinn, K. (2008). A screen of cell-surface molecules identifies leucine-rich repeat proteins as key mediators of synaptic target selection. Neuron, 59(6), 972–985.

Kvon, E. Z., Kazmar, T., Stampfel, G., Yáñez-Cuna, J. O., Pagani, M., Schernhuber, K., Dickson, B. J., & Stark, A. (2014). Genome-scale functional characterization of Drosophila developmental enhancers in vivo. Nature, 512(7512), 91–95.

Lai, S.-L., & Lee, T. (2006). Genetic mosaic with dual binary transcriptional systems in Drosophila. Nature Neuroscience, 9(5), 703–709.

Laurent, G., & Naraghi, M. (1994). Odorant-induced oscillations in the mushroom bodies of the locust. The Journal of Neuroscience: The Official Journal of the Society for Neuroscience, 14(5), 2993– 3004.

Lee, G., Foss, M., Goodwin, S. F., Carlo, T., Taylor, B. J., & Hall, J. C. (2000). Spatial, temporal, and sexually dimorphic expression patterns of the fruitless gene in the Drosophila central nervous system. Journal of Neurobiology, 43(4), 404–426.

Lee, T., Lee, A., & Luo, L. (1999). Development of the Drosophila mushroom bodies: Sequential generation of three distinct types of neurons from a neuroblast. Development, 126(18), 4065– 4076.

Leiss, F., Groh, C., Butcher, N. J., Meinertzhagen, I. A., & Tavosanis, G. (2009). Synaptic organization in the adult Drosophila mushroom body calyx. The Journal of Comparative Neurology, 517(6), 808–824.

Leutgeb, J. K., Leutgeb, S., Moser, M.-B., & Moser, E. I. (2007). Pattern separation in the dentate gyrus and CA3 of the hippocampus. Science, 315(5814), 961–966.

Li, F., Lindsey, J. W., Marin, E. C., Otto, N., Dreher, M., Dempsey, G., Stark, I., Bates, A. S., Pleijzier, M. W., Schlegel, P., Nern, A., Takemura, S.-Y., Eckstein, N., Yang, T., Francis, A., Braun, A., Parekh, R., Costa, M., Scheffer, L. K., … Rubin, G. M. (2020). The connectome of the adult Drosophila mushroom body provides insights into function. ELife, 9. 10.7554/eLife.62576

Li, Hanqing, Watson, A., Olechwier, A., Anaya, M., Sorooshyari, S. K., Harnett, D. P., Lee, H.-K. P., Vielmetter, J., Fares, M. A., Garcia, K. C., Özkan, E., Labrador, J.-P., & Zinn, K. (2017). Deconstruction of the beaten Path-Sidestep interaction network provides insights into neuromuscular system development. ELife, 6. 10.7554/eLife.28111

Li, Hongjie, Horns, F., Xie, Q., Xie, Q., Li, T., Luginbuhl, D. J., Luo, L., & Quake, S. R. (2017). Classifying Drosophila Olfactory Projection Neuron Subtypes by Single-Cell RNA Sequencing. Cell, 171(5), 1206.e22-1220.e22.

Li, Y., Lu, H., Cheng, P.-L., Ge, S., Xu, H., Shi, S.-H., & Dan, Y. (2012). Clonally related visual cortical neurons show similar stimulus feature selectivity. Nature, 486(7401), 118–121.

Lieberman, O. J., McGuirt, A. F., Tang, G., & Sulzer, D. (2019). Roles for neuronal and glial autophagy in synaptic pruning during development. Neurobiology of Disease, 122, 49–63.

Lin, A. C., Bygrave, A. M., De Calignon, A., Lee, T., & Miesenböck, G. (2014). Sparse, Decorrelated Odor Coding in the Mushroom Body Enhances Learned Odor Discrimination Andrew. Nature Neuroscience, 17(4), 559–568.

Lin, H.-H., Lai, J. S.-Y., Chin, A.-L., Chen, Y.-C., & Chiang, A.-S. (2007). A map of olfactory representation in the Drosophila mushroom body. Cell, 128(6), 1205–1217.

Lin, S., Kao, C.-F., Yu, H.-H., Huang, Y., & Lee, T. (2012). Lineage analysis of Drosophila lateral antennal lobe neurons reveals notch-dependent binary temporal fate decisions. PLoS Biology, 10(11), e1001425.

Litwin-Kumar, A., Harris, K. D., Axel, R., Sompolinsky, H., & Abbott, L. F. (2017). Optimal Degrees of Synaptic Connectivity. Neuron, 93(5), 1153–1164.e7.

Magklara, A., & Lomvardas, S. (2013). Stochastic gene expression in mammals: lessons from olfaction. Trends in Cell Biology, 23(9), 449–456.

Marin, E. C., Jefferis, G. S. X. E., Komiyama, T., Zhu, H., & Luo, L. (2002). Representation of the glomerular olfactory map in the Drosophila brain. Cell, 109(2), 243–255.

Markstein, M., Pitsouli, C., Villalta, C., Celniker, S. E., & Perrimon, N. (2008). Exploiting position effects and the gypsy retrovirus insulator to engineer precisely expressed transgenes. Nature Genetics, 40(4), 476–483.

Marr, D. (1969). A theory of cerebellar cortex. The Journal of Physiology, 202(2), 437–470.

Masse, N. Y., Turner, G. C., & Jefferis, G. S. X. E. (2009). Olfactory Information Processing in Drosophila. Current Biology: CB, 19(16), R700–R713.

Minsky, M. (1952). A Neural-Analogue Calculator Based upon a Probability Model of Reinforcement.

Minsky, M. (2016). Building my randomly wired neural network machine. https://www.webofstories.com/people/marvin.minsky/136?o=SH

Morano, N. C., Lopez, D. H., Meltzer, H., Sergeeva, A. P., Katsamba, P. S., Rostam, K. D., Gupta, H. P., Becker, J. E., Bornstein, B., Cosmanescu, F., Schuldiner, O., Honig, B., Mann, R. S., & Shapiro, L. (2024). *Cis*inhibition of co-expressed DIPs and Dprs shapes neural development. In bioRxiv (p. 2024.03.04.583391). 10.1101/2024.03.04.583391

Mugnaini, E., Osen, K. K., Dahl, A. L., Friedrich, V. L., Jr, & Korte, G. (1980). Fine structure of granule cells and related interneurons (termed Golgi cells) in the cochlear nuclear complex of cat, rat and mouse. Journal of Neurocytology, 9(4), 537–570.

Murthy, M., Fiete, I., & Laurent, G. (2008). Testing odor response stereotypy in the Drosophila mushroom body. Neuron, 59(6), 1009–1023.

Muthukumar, A. K., Stork, T., & Freeman, M. R. (2014). Activity-dependent regulation of astrocyte GAT levels during synaptogenesis. Nature Neuroscience, 17(10), 1340–1350.

Neuwirth, E. (2022). RColorBrewer: ColorBrewer Palettes (1.1-3) [Computer software]. https://CRAN.R-project.org/package=RColorBrewer

Osaka, J., Ishii, A., Wang, X., Iwanaga, R., Kawamura, H., Akino, S., Sugie, A., Hakeda-Suzuki, S., & Suzuki, T. (2024). Complex formation of immunoglobulin superfamily molecules Side-IV and Beat-IIb regulates synaptic specificity. Cell Reports, 43(2), 113798.

Özel, M. N., Simon, F., Jafari, S., Holguera, I., Chen, Y.-C., Benhra, N., El-Danaf, R. N., Kapuralin, K., Malin, J. A., Konstantinides, N., & Desplan, C. (2021). Neuronal diversity and convergence in a visual system developmental atlas. Nature, 589(7840), 88–95.

Özkan, E., Carrillo, R. A., Eastman, C. L., Weiszmann, R., Waghray, D., Johnson, K. G., Zinn, K., Celniker, S. E., & Garcia, K. C. (2013). An extracellular interactome of immunoglobulin and LRR proteins reveals receptor-ligand networks. Cell, 154(1), 228.

Palmateer, C. M., Artikis, C., Brovero, S. G., Friedman, B., Gresham, A., & Arbeitman, M. N. (2023). Single-cell transcriptome profiles of Drosophila fruitless-expressing neurons from both sexes. ELife, 12. 10.7554/eLife.78511

Pei, J., Kinch, L. N., & Grishin, N. V. (2018). FlyXCDB—A Resource for Drosophila Cell Surface and Secreted Proteins and Their Extracellular Domains. Journal of Molecular Biology, 430(18, Part B), 3353–3411.

Peters, A., & Feldman, M. L. (1976). The projection of the lateral geniculate nucleus to area 17 of the rat cerebral cortex. I. General description. Journal of Neurocytology, 5(1), 63–84.

Pfeiffer, B. D., Truman, J. W., & Rubin, G. M. (2012). Using translational enhancers to increase transgene expression in Drosophila. Proceedings of the National Academy of Sciences of the United States of America, 109(17), 6626–6631.

Pitman, J. L., Huetteroth, W., Burke, C. J., Krashes, M. J., Lai, S.-L., Lee, T., & Waddell, S. (2011). A pair of inhibitory neurons are required to sustain labile memory in the Drosophila mushroom body. Current Biology: CB, 21(10), 855–861.

Potter, C. J., Tasic, B., Russler, E. V., Liang, L., & Luo, L. (2010). The Q system: a repressible binary system for transgene expression, lineage tracing, and mosaic analysis. Cell, 141(3), 536–548.

R Foundation for Statistical Computing. (2023). R: A language and Environment for Statistical Computing (4.3.1) [Computer software]. https://www.R-project.org/

Rosenblatt, F. (1958). THE PERCEPTRON: A PROBABILISTIC MODEL FOR INFORMATION STORAGE AND ORGANIZATION IN THE BRAIN. Psychological Review, 65(6), 386–408.

Sanes, J. R., & Zipursky, S. L. (2020). Synaptic Specificity, Recognition Molecules, and Assembly of Neural Circuits. Cell, 181(3), 536–556.

Satija, R., Farrell, J. A., Gennert, D., Schier, A. F., & Regev, A. (2015). Spatial reconstruction of single-cell gene expression data. Nature Biotechnology, 33(5), 495–502.

Sawtell, N. B. (2010). Multimodal integration in granule cells as a basis for associative plasticity and sensory prediction in a cerebellum-like circuit. Neuron, 66(4), 573–584.

Scheffer, L. K., Xu, C. S., Januszewski, M., Lu, Z., Takemura, S.-Y., Hayworth, K. J., Huang, G. B., Shinomiya, K., Maitlin-Shepard, J., Berg, S., Clements, J., Hubbard, P. M., Katz, W. T., Umayam, L., Zhao, T., Ackerman, D., Blakely, T., Bogovic, J., Dolafi, T., … Plaza, S. M. (2020). A connectome and analysis of the adult Drosophila central brain. ELife, 9. 10.7554/eLife.57443

Schindelin, J., Arganda-Carreras, I., Frise, E., Kaynig, V., Longair, M., Pietzsch, T., Preibisch, S., Rueden, C., Saalfeld, S., Schmid, B., Tinevez, J.-Y., White, D. J., Hartenstein, V., Eliceiri, K., Tomancak, P., & Cardona, A. (2012). Fiji: an open-source platform for biological-image analysis. Nature Methods, 9(7), 676–682.

Shih, M. F. M., Davis, F. P., Henry, G. L., & Dubnau, J. (2019). Nuclear transcriptomes of the seven neuronal cell types that constitute the drosophila mushroom bodies. G3: Genes, Genomes, Genetics, 9(1), 81–94.

Shuster, S. A., Wagner, M. J., Pan-Doh, N., Ren, J., Grutzner, S. M., Beier, K. T., Kim, T. H., Schnitzer, M. J., & Luo, L. (2021). The relationship between birth timing, circuit wiring, and physiological response properties of cerebellar granule cells. Proceedings of the National Academy of Sciences of the United States of America, 118(23). 10.1073/pnas.2101826118

Sperry, R. W. (1963). Chemoaffinity in the orderly growth of nerve fiber patterns and connections. Proceedings of the National Academy of Sciences of the United States of America, 50(4), 703–710.

Srinivasan, S., & Stevens, C. F. (2018). The distributed circuit within the piriform cortex makes odor discrimination robust. The Journal of Comparative Neurology, 526(17), 2725–2743.

Stocker, R. F., Heimbeck, G., Gendre, N., & de Belle, J. S. (1997). Neuroblast ablation in Drosophila P[GAL4] lines reveals origins of olfactory interneurons. Journal of Neurobiology, 32(5), 443– 456.

Stuart, T., Butler, A., Hoffman, P., Hafemeister, C., Papalexi, E., Mauck, W. M., 3rd, Hao, Y., Stoeckius, M., Smibert, P., & Satija, R. (2019). Comprehensive integration of single-cell data. Cell, 177(7), 1888–1902.e21.

Tan, L., Zhang, K. X., Pecot, M. Y., Nagarkar-Jaiswal, S., Lee, P. T., Takemura, S. Y., McEwen, J. M., Nern, A., Xu, S., Tadros, W., Chen, Z., Zinn, K., Bellen, H. J., Morey, M., & Zipursky, S. L. (2015). Ig Superfamily Ligand and Receptor Pairs Expressed in Synaptic Partners in Drosophila. Cell, 163(7), 1756–1769.

Tanaka, N. K., Awasaki, T., Shimada, T., & Ito, K. (2004). Integration of chemosensory pathways in the Drosophila second-order olfactory centers. Current Biology: CB, 14(6), 449–457.

Tirian, L., & Dickson, B. J. (2017). The VT GAL4, LexA, and split-GAL4 driver line collections for targeted expression in the *Drosophilanervous* system. In medRxiv (p. 198648). bioRxiv. 10.1101/198648

Van der Loos, H., & Glaser, E. M. (1972). Autapses in neocortex cerebri: synapses between a pyramidal cell’s axon and its own dendrites. Brain Research, 48, 355–360.

Vogt, K., Aso, Y., Hige, T., Knapek, S., Ichinose, T., Friedrich, A. B., Turner, G. C., Rubin, G. M., & Tanimoto, H. (2016). Direct neural pathways convey distinct visual information to drosophila mushroom bodies. ELife, 5(APRIL2016), 1–13.

Wang, J., Zugates, C. T., Liang, I. H., Lee, C.-H. J., & Lee, T. (2002). Drosophila Dscam is required for divergent segregation of sister branches and suppresses ectopic bifurcation of axons. Neuron, 33(4), 559–571.

Williams, D. L., Sikora, V. M., Hammer, M. A., Amin, S., Brinjikji, T., Brumley, E. K., Burrows, C. J., Carrillo, P. M., Cromer, K., Edwards, S. J., Emri, O., Fergle, D., Jenkins, M. J., Kaushik, K., Maydan, D. D., Woodard, W., & Clowney, E. J. (2021). May the odds be ever in your favor: Non-deterministic mechanisms diversifying cell surface molecule expression. Frontiers in Cell and Developmental Biology, 9, 720798.

Wilton, D. K., Dissing-Olesen, L., & Stevens, B. (2019). Neuron-Glia signaling in synapse elimination. Annual Review of Neuroscience, 42(1), 107–127.

Wolf, F. A., Angerer, P., & Theis, F. J. (2018). SCANPY: large-scale single-cell gene expression data analysis. Genome Biology, 19(1), 15.

Wolterhoff, N., & Hiesinger, P. R. (2024). Synaptic promiscuity in brain development. Current Biology: CB, 34(3), R102–R116.

Wong, A. M., Wang, J. W., & Axel, R. (2002). Spatial representation of the glomerular map in the Drosophila protocerebrum. Cell, 109(2), 229–241.

Xie, Q., Brbic, M., Horns, F., Kolluru, S. S., Jones, R. C., Li, J., Reddy, A. R., Xie, A., Kohani, S., Li, Z., McLaughlin, C. N., Li, T., Xu, C., Vacek, D., Luginbuhl, D. J., Leskovec, J., Quake, S. R., Luo, L., & Li, H. (2021). Temporal evolution of single-cell transcriptomes of Drosophila olfactory projection neurons. ELife, 10. 10.7554/eLife.63450

Xie, Q., Wu, B., Li, J., Xu, C., Li, H., Luginbuhl, D. J., Wang, X., Ward, A., & Luo, L. (2019). Transsynaptic Fish-lips signaling prevents misconnections between nonsynaptic partner olfactory neurons. Proceedings of the National Academy of Sciences of the United States of America, 116(32), 16068–16073.

Xu, C., Theisen, E., Maloney, R., Peng, J., Santiago, I., Yapp, C., Werkhoven, Z., Rumbaut, E., Shum, B., Tarnogorska, D., Borycz, J., Tan, L., Courgeon, M., Griffin, T., Levin, R., Meinertzhagen, I. A., de Bivort, B., Drugowitsch, J., & Pecot, M. Y. (2019). Control of Synaptic Specificity by Establishing a Relative Preference for Synaptic Partners. Neuron, 103(5), 865–877.e7.

Xu, H.-T., Han, Z., Gao, P., He, S., Li, Z., Shi, W., Kodish, O., Shao, W., Brown, K. N., Huang, K., & Shi, S.-H. (2014). Distinct lineage-dependent structural and functional organization of the hippocampus. Cell, 157(7), 1552–1564.

Yagi, R., Mabuchi, Y., Mizunami, M., & Tanaka, N. K. (2016). Convergence of multimodal sensory pathways to the mushroom body calyx in Drosophila melanogaster. Scientific Reports, 6(July), 1–8.

Yang, J.-Y., O’Connell, T. F., Hsu, W.-M. M., Bauer, M. S., Dylla, K. V., Sharpee, T. O., & Hong, E. J. (2023). Restructuring of olfactory representations in the fly brain around odor relationships in natural sources. In bioRxiv (p. 2023.02.15.528627). 10.1101/2023.02.15.528627

Yang, K., Liu, T., Wang, Z., Liu, J., Shen, Y., Pan, X., Wen, R., Xie, H., Ruan, Z., Tan, Z., Chen, Y., Guo, A., Liu, H., Han, H., Di, Z., & Zhang, K. (2022). Classifying Drosophila olfactory projection neuron boutons by quantitative analysis of electron microscopic reconstruction. IScience, 25(5), 104180.

Yoo, J., Dombrovski, M., Mirshahidi, P., Nern, A., LoCascio, S. A., Zipursky, S. L., & Kurmangaliyev, Y. Z. (2023). Brain wiring determinants uncovered by integrating connectomes and transcriptomes. Current Biology: CB, 33(18), 3998–4005.e6.

Yu, H.-H., Kao, C.-F., He, Y., Ding, P., Kao, J.-C., & Lee, T. (2010). A complete developmental sequence of a Drosophila neuronal lineage as revealed by twin-spot MARCM. PLoS Biology, 8(8), e1000461.

Yu, Y.-C., Bultje, R. S., Wang, X., & Shi, S.-H. (2009). Specific synapses develop preferentially among sister excitatory neurons in the neocortex. Nature, 458(7237), 501–504.

Yusuyama, K., Meinertzhagen, I. A., & Schürmann, F. W. (2002). Synaptic organization of the mushroom body calyx in Drosophila melanogaster. The Journal of Comparative Neurology, 445(3), 211–226.

Zappia, L., & Oshlack, A. (2018). Clustering trees: a visualization for evaluating clusterings at multiple resolutions. GigaScience, 7(7), giy083.

Zhan, X.-L., Clemens, J. C., Neves, G., Hattori, D., Flanagan, J. J., Hummel, T., Vasconcelos, M. L., Chess, A., & Zipursky, S. L. (2004). Analysis of Dscam diversity in regulating axon guidance in Drosophila mushroom bodies. Neuron, 43(5), 673–686.

Zheng, Z., Lauritzen, J. S., Perlman, E., Robinson, C. G., Nichols, M., Milkie, D., Torrens, O., Price, J., Fisher, C. B., Sharifi, N., Calle-Schuler, S. A., Kmecova, L., Ali, I. J., Karsh, B., Trautman, E. T., Bogovic, J. A., Hanslovsky, P., Jefferis, G. S. X. E., Kazhdan, M., … Bock, D. D. (2018). A Complete Electron Microscopy Volume of the Brain of Adult Drosophila melanogaster. Cell, 174(3), 730–743.e22.

Zheng, Z., Li, F., Fisher, C., Ali, I. J., Sharifi, N., Calle-Schuler, S., Hsu, J., Masoodpanah, N., Kmecova, L., Kazimiers, T., Perlman, E., Nichols, M., Li, P. H., Jain, V., & Bock, D. D. (2022). Structured sampling of olfactory input by the fly mushroom body. Current Biology: CB, 32(15), 3334–3349.e6.

Zhu, S., Chiang, A.-S., & Lee, T. (2003). Development of the Drosophila mushroom bodies: elaboration, remodeling and spatial organization of dendrites in the calyx. Development, 130(12), 2603–2610.

Zirin, J., Hu, Y., Liu, L., Yang-Zhou, D., Colbeth, R., Yan, D., Ewen-Campen, B., Tao, R., Vogt, E., VanNest, S., Cavers, C., Villalta, C., Comjean, A., Sun, J., Wang, X., Jia, Y., Zhu, R., Peng, P., Yu, J., … Perrimon, N. (2020). Large-scale transgenic Drosophila resource collections for loss- and gain-of-function studies. Genetics, 214(4), 755–767.

